# APOE Christchurch enhances a disease-associated microglial response to plaque but suppresses response to tau pathology

**DOI:** 10.1101/2024.06.03.597211

**Authors:** Kristine M. Tran, Nellie Kwang, Angela Gomez-Arboledas, Shimako Kawauchi, Cassandra Mar, Donna Chao, Celia Da Cunha, Shuling Wang, Sherilyn Collins, Amber Walker, Kai-Xuan Shi, Joshua A. Alcantara, Jonathan Neumann, Andrea J. Tenner, Frank M. LaFerla, Lindsay A. Hohsfield, Vivek Swarup, Grant R. MacGregor, Kim N. Green

**Affiliations:** Department of Neurobiology and Behavior, University of California, Irvine, CA 92697, USA; Institute for Memory Impairments and Neurological Disorders, University of California, Irvine, CA 92697, USA; Transgenic Mouse Facility, ULAR, Office of Research, University of California, Irvine, CA 92697, USA; Department of Developmental and Cell Biology, University of California, Irvine, CA 92697, USA; Center for Complex Biological Systems, University of California, Irvine, CA 92697, USA; Department of Pharmaceutical Sciences, University of California, Irvine, CA 92697, USA; Department of Molecular Biology & Biochemistry, University of California, Irvine, CA 92697, USA; Department of Pathology and Laboratory Medicine, University of California, Irvine, CA 92697, USA

**Keywords:** APOE Christchurch, PS19, 5xFAD, microglia, DAM, Amyloid, Tau, resilience

## Abstract

**Background:** Apolipoprotein E ε4 (APOE4) is the strongest genetic risk factor for late-onset Alzheimer’s disease (LOAD). A recent case report identified a rare variant in APOE, APOE3-R136S (Christchurch), proposed to confer resistance to autosomal dominant Alzheimer’s Disease (AD). However, it remains unclear whether and how this variant exerts its protective effects.

**Methods:** We introduced the R136S variant into mouse *Apoe* (*ApoeCh*) and investigated its effect on the development of AD-related pathology using the 5xFAD model of amyloidosis and the PS19 model of tauopathy. We used immunohistochemical and biochemical analysis along with single-cell spatial transcriptomics and proteomics to explore the impact of the *ApoeCh* variant on AD pathological development and the brain’s response to plaques and tau.

**Results:** In 5xFAD mice, *ApoeCh* enhances a Disease-Associated Microglia (DAM) phenotype in microglia surrounding plaques, and reduces plaque load, dystrophic neurites, and plasma neurofilament light chain. By contrast, in PS19 mice, *ApoeCh* suppresses the microglial and astrocytic responses to tau-laden neurons and does not reduce tau accumulation or phosphorylation, but partially rescues tau-induced synaptic and myelin loss. We compared how microglia responses differ between the two mouse models to elucidate the distinct DAM signatures induced by *ApoeCh*. We identified upregulation of antigen presentation-related genes in the DAM response in a PS19 compared to a 5xFAD background, suggesting a differential response to amyloid versus tau pathology that is modulated by the presence of *ApoeCh*.

**Conclusions:** These findings highlight the ability of the *ApoeCh* variant to modulate microglial responses based on the type of pathology, enhancing DAM reactivity in amyloid models and dampening neuroinflammation to promote protection in tau models. This suggests that the Christchurch variant’s protective effects likely involve multiple mechanisms, including changes in receptor binding and microglial programming.

## BACKGROUND

Alzheimer’s Disease (AD), the most prevalent neurodegenerative disease, is characterized by a progressive decline in cognitive function accompanied by the development of extracellular neurotoxic amyloid-beta (Aβ) plaques and intracellular neurofibrillary tangles (NFT) [1]. AD can be classified into subtypes based on age of symptom onset: early-onset AD (EOAD) and late-onset AD (LOAD), and disease genetics: genetically determined and sporadic AD. Genetic factors can determine an individual’s susceptibility to AD, with rare autosomal dominant AD (ADAD) cases manifesting within families carrying pathogenic variants of *Amyloid Precursor Protein (APP), Presenilin 1 (PSEN1), and Presenilin 2 (PSEN2)* genes [2–7]. Polymorphisms in additional genes have been identified as major risk factors for developing both EOAD and LOAD, such as *Apolipoprotein E* (*APOE)* [8].

APOE is a soluble glycoprotein secreted by cells in both the central nervous system (CNS) and periphery. APOE is the most abundant apolipoprotein in the brain and is normally expressed at highest levels by astrocytes, and lower levels by microglia, oligodendrocytes, pericytes, endothelial cells, choroid plexus epithelial cells, and neurons. APOE mediates lipid transport and delivery between cell types in the CNS. Mature APOE is a 299 amino acid (AA) protein comprised of three domains – an N-terminal region with four alpha-helices that includes a receptor binding region (AA136-150), a central hinge domain, and a C-terminal domain that can bind lipid (AA244-272). APOE is the most robust genetic risk factor for sporadic LOAD. In humans, there are 3 different isoforms of APOE: ɛ2, ɛ3, and ɛ4, which differ in the presence of a cysteine or arginine at position 112 and 158 within the N-terminal domain. Inheritance of an APOE4 allele causes a 6 to 18-fold increase in AD risk compared to APOE2/3 [9]. Both human and animal studies suggest there are isoform-specific effects of APOE on neuroinflammation, tau pathology, and amyloid plaque dynamics, possibly due to differences in receptor binding affinity and lipid interactions [10, 11].

In 2019, a case report [12] provided striking evidence that APOE is an essential mediator of plaque-induced tau pathology and subsequent neurodegeneration. The proband was heterozygous for the ADAD-associated PSEN1 E280A variant and was anticipated to develop AD by her 40s although she remained cognitively intact until her seventies. Positron emission tomography (PET) imaging revealed significant Aβ plaque deposits in her brain, consistent with carriers of FAD mutations, but a relatively low and atypical burden of neurofibrillary tau tangles and neurodegeneration. Further investigation revealed she was homozygous for the rare APOE3-Christchurch (APOE3Ch) variant in APOE3 (APOE R136S), suggesting this might account for her cognitive preservation [12]. This key study demonstrated that the presence of plaques can be separated from the development of AD. Understanding the mechanisms by which APOE3Ch prevents the subsequent development of tau pathology and neurodegeneration is essential to stop dementia during the long plaque-laden prodromal phase of AD.

The R136S mutation resides in a region of APOE involved in binding to lipoprotein receptors, including low-density lipid receptor (LDLR), lipoprotein-related receptor 1 (LRP1), and heparan sulfate proteoglycans (HSPGs), all implicated in promoting amyloid-β aggregation and neuronal uptake of extracellular tau. Studies indicate that APOE binding is integral to some of these processes [13–17] with differences in APOE isoform-specific receptor binding affinity and lipid interactions affecting neuroinflammation, tau pathology, and amyloid plaque dynamics [10, 11, 18, 19]. Collectively, these recent findings underscore the influence of *APOE* and *APOECh* on both Aβ and tau pathology, either directly or through downstream effects such as neuroinflammation. The *APOECh* variant offers a unique opportunity to explore the interplay between Aβ and tau, given its potential association with limited tau development and resistance to cognitive decline despite increased Aβ levels. Mechanistically, APOE may modulate the immune response to plaques, preventing aspects that drive tau pathology, or it may play a pivotal role in inducing and spreading tau pathology and subsequent neurodegeneration. To test these hypotheses, we developed an *Apoe*\*R136S variant in mice, and independently introduced it into both an aggressive mouse model of amyloidosis (5xFAD mice), and of tauopathy (PS19 mice). Here we describe the effects of the *Apoe* Christchurch variant on the development of both plaques and tau pathology, as well as its impact on the brains’ response to these pathologies.

## METHODS

### Animals

All experiments involving mice were approved by the UC Irvine Institutional Animal Care and Use Committee and were conducted in compliance with all relevant ethical regulations for animal testing and research. All experiments involving mice comply with the Animal Research: Reporting of in Vivo Experiments (ARRIVE-10) guidelines.

### Environmental conditions

Animals were housed in autoclaved individual ventilated cages (SuperMouse 750, Lab Products, Seaford, DE) containing autoclaved corncob bedding (Envigo 7092BK 1/8” Teklad, Placentia, CA) and two autoclaved 2” square cotton nestlets (Ancare, Bellmore, NY) plus a LifeSpan multi-level environmental enrichment platform. Tap water (acidified to pH2.5-3.0 with HCl then autoclaved) and food (LabDiet Mouse Irr 6F; LabDiet, St. Louis, MO) were provided *ad libitum*. Cages were changed every 2 weeks with a maximum of 5 adult animals per cage. Room temperature was maintained at 72 ± 2°F, with ambient room humidity (average 40-60% RH, range 10 - 70%). Light cycle was 14h light / 10h dark, lights on at 06.30h and off at 20.30h.

### Mice

CRISPR/Cas9 endonuclease-mediated genome editing was used to introduce a CGG to TCT missense mutation resulting in an arginine to serine substitution (R128S) in the mouse apolipoprotein E (*Apoe*) gene. This models the Christchurch R136S missense variant in human *APOE* (**Supplemental Fig. 1a**). Alt-R Crispr RNA (TMF1648 – gcacagaggagatacgggcg) and tracrRNA plus CAS9 protein (HiFi Cas9 nuclease V3, Integrated DNA Technologies (IDT), Coralville, IA) as a ribonucleoprotein (RNP) was electroporated into C57BL/6J (B6J) zygotes (Jackson Lab stock # 000664, Bar Harbor, ME) along with a ssODN sequence (TMF 1341–5’-ACCATGCTGGGCCAGAGCACAGAGGAGATAAGAGCATCTCTCTCCACACACCTGCGCAAGATGCGCAAGCG —3’). G0 founder animals containing the desired DNA sequence changes were backcrossed with B6J mice and N1 heterozygous mice were sequenced to determine the mutant allele. N1 heterozygous mice were backcrossed with B6J mice to produce N2F1 heterozygotes, which were used to generate animals for subsequent analysis. The formal allele name is *Apoe^em1Aduci^* (Jackson Laboratory stock 39301) For brevity, hereafter we refer to the allele as *ApoeCh*.

5xFAD hemizygous (B6.CgTg(APPSwFlLon,PSEN1*M146L*L286V)6799Vas/Mmjax, Jackson Lab stock # 34848) mice and PS19 hemizygous mice (B6.Cg-Tg(Prnp-MAPT*P301S)PS19Vle/J; Jackson Lab stock # 24841) were obtained from The Jackson Lab. The experimental cohort for 5xFAD hemizygous (HEMI); *ApoeCh* homozygous (HO) and littermate control *ApoeCh* HO mice was derived from natural mating of 5xFAD HEMI; *ApoeCh* HO with *ApoeCh* HO mice. 5xFAD HEMI control mice were generated via *in vitro* fertilization and embryo transfer to pseudo pregnant surrogate females. Non-transgenic wildtype mice were obtained through natural mating of B6J mice. Similarly, all four groups of the PS19;*ApoeCh* cohort were generated from natural mating of B6J *ApoeCh* heterozygous (HET) mice with (C57BL/6 x C3H)F1 PS19 HEMI mice with subsequent breeding of mixed background PS19 HEMI; *ApoeCh* HET with *ApoeCh* HET. Post-weaning, mice were group-housed with littermates and aged until the designated harvest dates. All animals were bred in the Transgenic Mouse Facility at UCI.

*ApoeCh* genotyping was done using a common primer set to amplify both the *Apoe* wildtype and the *ApoeCh* alleles (For 5′-GGAGGACACTATGACGGAAGTA-3′ and Rev 5′-TGCCTTGTACACAGCTAGGC-3′). Two fluorophore labeled-hydrolysis probes were used to identify the mouse *Apoe* wildtype amplicon (5’-AGCACAGAGGAGATACGGGCGC −3’ - HEX) and the *ApoeCh* amplicon (5’-AGCACAGAGGAGATAAGAGCATC −3’-FAM). The relative fluorescence from each probe was quantified at the end point of PCR cycles to assign genotype using Allelic Discrimination function in Bio-Rad CFX Maestro software (Bio-Rad, Hercules, CA). For 5xFAD genotyping, a hydrolysis probe that hybridizes to the APP(Swe) amplicon was used (For 5′-TGGGTTCAAACAAAGGTGCAA-3′ and Rev 5′-GATGACGATCACTGTCGCTATGAC-3′: APP(Swe) probe 5′-CATTGGACTCATGGTGGGCGGTG-3′). Ct values were normalized to amplification of the *Apob* locus (For 5′-CACGTGGGCTCCAGCATT-3′ and Rev 5′-TCACCAGTCATTTCTGCCTTTG-3′: *ApoB* probe 5′-CCAATGGTCGGGCACTGCTCAA-3′).

Genotyping for the PS19 transgene was performed using two primer sets to amplify both the wildtype (For 5’-CAAATGTTGCTTGTCTGGTG-3’ and Rev 5’-GTCAGTCGAGTGCACAGTTT) and P301S (For 5’-GGCATCTCAGCAATGTCTCC-3’ and Rev 5’-GGTATTAGCCTATGGGGGACAC-3’) using cycling conditions from The Jackson Laboratory (mouse stock # 24841).

### Off-target analysis

Genomic DNA was extracted from mouse tail biopsies using DirectPCR Lysis Reagent (Viagen Biotech, Los Angeles, CA) and Proteinase K (Roche, Indianapolis, IN). Amplification was performed using a Bio-Rad CFX-96 instrument. For each amplicon, a single PCR product was confirmed by capillary electrophoresis (5200 Fragment Analyzer, Agilent, Santa Clara, CA) then subjected to Sanger sequencing (Retrogen, San Diego, CA) and analyzed using SeqMan Ultra 17.4 (DNASTAR, Madison, WI). Potential off-target sites are listed in Supplementary Table 1 while primers for PCR amplification and sequencing of potential off-target sites are listed in Supplemental Table 2.

### Behavioral assays

Noldus Ethovision software (Wageningen, Netherlands) was used for video recording and tracking of animal behavior, and subsequent analyses. Detailed protocols are openly accessible via the AD Knowledge Portal (https://adknowledgeportal.synapse.org/). The following behavioral assays were executed following established methodologies as briefly outlined below [20]:

#### Hindlimb clasping

Mice were lifted by the base of their tails while clear of all surrounding objects.

They were observed for 10 sec and scored based on their hindlimb position as described [21]. Scoring was done by observers blinded to genotype.

#### Elevated plus maze (EPM)

To evaluate anxiety, mice were positioned at the center of an elevated plus maze (arms 6.2 × 75 cm, with 20 cm side walls on two closed arms, elevated 63 cm above the ground) for 5 min. Automated scoring quantified the cumulative time spent by each mouse in the open and closed arms of the maze.

#### Open field (OF)

To assess both motor function and anxiety, mice were placed within a white box (33.7cm L × 27.3cm W × 21.6cm H) for 5 min with movement recorded via video. Videos were subsequently analyzed to quantify the percentage of time spent in the center of the arena, total distance traveled, and speed.

### Plasma lipid measurement

Blood plasma was collected and analyzed using the Piccolo® blood chemistry analyzer from Abaxis (Union City, CA) following the manufacturer’s instructions. Plasma was diluted 1:1 with distilled water (ddH_2_O), and then 100 µl of the diluted sample was loaded onto the Piccolo lipid plus panel plate (#07P0212, Abaxis). Various lipid parameters including total cholesterol (CHOL), high-density lipoprotein (HDL), non-HDL cholesterol (nHDLc), triglycerides, low-density lipoprotein (LDL), and very low-density lipoprotein (vLDL) were analyzed and plotted. Lipid and general chemistry controls (#07P0401, Abaxis) were utilized to verify accuracy and reproducibility of the measurements.

### Tissue preparation for histology

Mice were euthanized by CO_2_ inhalation (5xFAD at 4 and 12 months of age, and PS19 at 5 and 9 months of age), then transcardially perfused with 1X phosphate buffered saline (PBS). In all experiments, the brains were dissected and the hemispheres divided along the midline. One half of each brain was fresh-frozen in dry-ice cooled isopentane at −40°C for spatial single-cell transcriptomics, while the other half was fixed in 4% paraformaldehyde (PFA) in PBS (Thermo Fisher Scientific, Waltham, MA) for 24hr at 4°C for immunohistochemical analysis.

### Immunohistochemistry

PFA-fixed brain halves were sectioned into 40μm coronal slices using a Leica SM2000R freezing microtome, and slices taken for immunohistological analysis (between Bregma −2.78 mm and – 3.38 mm for 5xFAD and between Bregma −1.58 mm and −2.7 mm for PS19 according to the Allen Mouse Brain Atlas, Reference Atlas version 1, 2008). One representative brain slice from each mouse within the identical experimental cohort (comprising the same genotype, age, and sex) was subjected to simultaneous staining in the same container as described [20, 22]. The free-floating sections underwent a series of washes, at room temperature unless otherwise stated: three times in 1X phosphate-buffered saline (PBS) for 10 min, 5 min, and 5 min, successively.

For AmyloGlo staining, after the PBS washes, free-floating brain slices were washed in 70% ethanol for 5 min, followed by a rinse in deionized water for 2 min, before being immersed in Amylo-Glo RTD Amyloid Plaque Staining Reagent (diluted 1:100 in 0.9% saline solution; TR-200-AG; Biosensis, Thebarton, South Australia) for 10 min. Post-incubation, the sections were subjected to a 5-min wash in 0.9% saline solution and a rinse in deionized water for 15 sec before proceeding with a standardized indirect immunohistochemical protocol. Subsequently, the sections were immersed in blocking serum solution (5% normal goat serum with 0.2% Triton X-100 in 1X PBS) for 1 hr, followed by overnight incubation at 4°C in primary antibodies diluted in blocking serum solution. For synaptic staining, free-floating sections were washed three times in 1X PBS then incubated in blocking serum solution overnight at 4°C, followed by overnight incubation at 4°C in primary antibodies diluted in blocking serum solution. Finally, the sections were incubated with secondary antibodies for 1hr in the dark followed by 3 washes in 1X PBS, before mounting on microscope slides.

Brain sections were stained using a standard indirect technique [20, 23] with the following primary antibodies against: ionized calcium-binding adapter molecule 1 (IBA1; 1:2000; 019–19,741; Wako, Osaka, Japan), glial fibrillary acidic protein (GFAP; 1:1000; AB134436; Abcam, Cambridge, MA, United States), lysosome-associated membrane protein 1 (LAMP1; 1:200; AB25245, Abcam), CD11c (1:100; 50–112-2633; eBioscience, San Diego, CA, United States), phospho-Tau Ser202, Thr205 (AT8; 1:500; MN1020; Invitrogen, Waltham, MA, United States), phospho-MLKL Ser358 (pMLKL; 1:400; PA5-105678; Invitrogen), FOX 3 (NeuN; 1:1000; AB104225; Abcam), Bassoon (1:250; 75-491; Antibodies Inc, Davis, CA, United States), Homer1 (1:250; 160003; Synaptic Systems, Gottingen, Germany) and BODIPY (1:1000; D3922; Invitrogen).

Brain sections were imaged with a Zeiss Axio Scan Z1 Slidescanner using a 10x 0.45 NA Plan-Apo objective. Images were also obtained using a 20x, 0.75 NA objective on a Leica TCS SPE-II confocal microscope and analyzed using Bitplane Imaris Software.

For super-resolution imaging, the CA1-SR hippocampal region of WT, *ApoeCh*, PS19 and PS19;*ApoeCh* 9 months old mice were acquired with a Zeiss LSM 900 Airyscan 2 microscope and Zen image acquisition software (Zen Blue, Carl Zeiss, White Plains, NY) with identical conditions. All images were collected using a 63x 1.4 NA Plan-Apo oil objective and 3 Z-stacks (180 nm step interval, within a depth of 4-6 μm) per mouse/sex/genotype was acquired to obtain a full representation of the CA1-SR. Airyscan processing of all channels and z-stack images was performed in Zen software and Bitplane Imaris Software used for quantification.

### Biochemical analysis of Aβ, tau, and neurofilament light chain

Sample preparation and quantification of Aβ followed established protocols [20, 23]. The hippocampal and cortical regions of each mouse brain hemisphere were micro-dissected and flash-frozen. Samples were pulverized using a Bessman Tissue Pulverizer. For Aβ biochemical analysis, pulverized hippocampal tissue was homogenized in 150µL of Tissue Protein Extraction Reagent (TPER; Life Technologies, Grand Island, NY), while cortical tissue was homogenized in 1000µL/150 mg of TPER. This formulation of TPER, containing 25 mM bicine and 150 mM sodium chloride (pH 7.6), effectively solubilizes proteins within brain tissue post-homogenization. Protease (Roche) and phosphatase inhibitors (Sigma-Aldrich) were added to the homogenized samples, which were then centrifuged at 100,000g for 1 hr at 4°C to generate TPER-soluble fractions. To generate formic acid fractions, pellets from TPER-soluble fractions were homogenized in 70% formic acid: 75µL for hippocampal tissue or half of the TPER volume used for cortical tissue. Subsequently, samples were again centrifuged at 100,000g for 1 hr at 4°C. Protein in the insoluble fraction of micro-dissected hippocampal and cortical tissue was normalized to the respective brain region weight, while protein in soluble fractions was normalized to the protein concentration determined via Bradford Protein Assay. Formic acid neutralization buffer (1M TRIS base, 0.5M Na_2_HPO_4_, 10% NaN_3_) was used to adjust pH before running ELISAs.

Tau protein extraction from PS19 mice was performed as described [24, 25]. Briefly, microdissected hippocampal and cortical tissues were homogenized in 10uL high salt reassembly buffer (RAB buffer; C752K77; ThermoFisher; 100 mM MES, 1 mM EGTA, 0.5 mM MgSO_4_, 750 mM NaCl, 20 mM NaF, 1 mM Na_3_VO_4_, pH = 7.0) with protease (Roche) and phosphatase inhibitors (Sigma-Aldrich) per 1 mg sample then centrifuged at 50,000g for 20 min to obtain the supernatant as the RAB-soluble fraction. The pellet was then dissolved in 10uL lysis radioimmunoprecipitation assay (RIPA) buffer with EDTA (CAT# J61529.AP; ThermoFisher; 150 mM NaCl, 50 mM Tris, 0.5% deoxycholic acid, 1% Triton X-100, 0.1% SDS, 5 mM EDTA, 20 mM NaF, 1 mM Na_3_VO_4_, pH 8.0) with protease and phosphatase inhibitor per 1mg sample and centrifuged at 50,000g for 20 min to obtain a RIPA-soluble fraction. For formic acid fractions, pellets from RIPA-soluble fractions were homogenized in 70% formic acid: 10µL per 1mg of tissue. Samples were then centrifuged at 50,000g for 20 min. Protein in the insoluble fraction of micro-dissected hippocampal and cortical tissue was normalized to the respective brain region weight, while protein in soluble fractions was normalized to the protein concentration determined via Bradford Protein Assay.

Quantitative biochemical analyses of human Aβ soluble and insoluble fraction levels were acquired using the V-PLEX Aβ Peptide Panel 1 (6E10) (K15200G-1; Meso Scale Discovery, Rockville, MD). Tau protein levels were obtained using the Phospho (Thr231)/Total Tau Kit (K15121D; Meso Scale Discovery). Quantitative biochemical analysis of neurofilament-light chain (NfL) in plasma was performed using the R-Plex Human Neurofilament L Assay (K1517XR-2; Meso Scale Discovery).

### Imaris quantitative analysis

Confocal images of each brain region were quantified automatically using the spots and surfaces module within Imaris v9.7. Amyloid burden was assessed by measuring the total AmyloGlo+ plaque number and volume from 10x 0.45 NA Zeiss Axio Scan Z1 Slidescanner images. Similarly, volumetric measurements (i.e., AmyloGlo+ plaque, IBA1+ microglia volume, etc.) were acquired automatically utilizing the surfaces module with confocal images of each brain region. Quantitative comparisons between experimental groups were carried out in sections stained simultaneously. For synaptic quantification, the total number of Bassoon or Homer1 puncta was quantified using the spots function of Imaris v9.6. Co-localization of pre- and post-synaptic puncta (Bassoon-Homer1) was determined using the spots function on Imaris and Matlab software to determine the total number of colocalized spots (defined as ≤ 200 nm distance). Results were normalized to the total volume of each image, to correct for any difference in the depth of imaging. Quantitative comparisons between experimental groups were carried out in sections stained simultaneously with the same imaging settings.

### FIJI ImageJ Analysis

Prior to analysis, all images were converted to 8-bit gray-scale using Fiji ImageJ. The threshold feature was adjusted and used to distinguish signal from background before percent coverage was measured. To quantify AT8 percent area coverage in the dentate gyrus, the whole FOV of the 20x confocal images of the dentate gyrus were analyzed. For quantification of NEUN percent coverage in the CA3 and percent plaque load in the entire brain section, whole brain-stitched 10x slidescanned images were used with appropriate brain regions outlined to define the region of interest.

### Single-cell Spatial Transcriptomics and Proteomics

Isopentane fresh-frozen brain hemispheres were embedded in optimal cutting temperature (OCT) compound (Tissue-Tek, Sakura Fintek, Torrance, CA), and 10um thick coronal sections were prepared using a cryostat (CM1950, LeicaBiosystems, Deer Park, IL). Six hemibrains were mounted onto each VWR Superfrost Plus microscope slide (Avantor, 48311-703) and kept at - 80°C until fixation. For both 5xFAD (14 months old) and PS19 (9 months old) models, n=3 mice per genotype except for n=2 for PS19;*ApoeCh* (wild-type, *ApoeCh* HO, 5xFAD HEMI or PS19 HEMI, and 5xFAD HEMI; *ApoeCh* HO or PS19 HEMI;*ApoeCh* HO) were used for transcriptomics and proteomics. The same mice were used for both transcriptomics and proteomics. Tissues were processed according to the Nanostring CosMx fresh-frozen slide preparation manual for RNA and protein assays (NanoString University).

#### Slide preparation for spatial transcriptomics

Slides were immersed in 10% neutral buffered formalin (NBF; CAT#15740) for 2 hr at 4°C, washed three times in 1X PBS (pH 7.4) for 2 min each, then baked at 60°C for 30 min. Slides were processed as follows: three washes of 1X PBS for 5 min each, 4% sodium dodecyl sulfate (SDS; CAT#AM9822) for 2 min, three washes of 1X PBS for 5 min each, 50% ethanol for 5 min, 70% ethanol for 5 min, and two washes of 100% ethanol for 5 min each, before air drying for 10 min at room temperature. Antigen retrieval was performed in a pressure cooker at 100°C for 15 min in 1X CosMx Target Retrieval Solution (Nanostring, Seattle, WA). Slides were transferred to DEPC-treated water (CAT#AM9922) and washed for 15 sec, incubated in 100% ethanol for 3 min, then air-dried for 30 min. Each slide was incubated with digestion buffer (3 µg/mL Proteinase K in 1X PBS; Nanostring) for tissue permeabilization, then washed twice in 1X PBS for 5 min each. Fiducials for image alignment were diluted to 0.00015% in 2X SSC-T and applied to the slide, then incubated for 5 min. Tissues were then post-fixed with the following washes: 10% NBF for 1 min, two washes of NBF Stop Buffer (0.1M Tris-Glycine Buffer, CAT#15740) for 5 min each, and 1x PBS for 5 min. Next, NHS-Acetate (100 mM; CAT#26777) mixture was applied to each slide and incubated for 15 min at RT. Slides were washed twice in 2X SSC for 5 min each. Slides were incubated with a modified 1000-plex Mouse Neuro RNA panel (Nanostring) for *in situ* hybridization along with an rRNA segmentation marker in a hybridization oven at 37°C for 16-18 hr overnight. Following overnight *in situ* hybridization, slides were washed twice in a stringent wash solution (50% deionized formamide [CAT#AM9342], 2X saline-sodium citrate [SSC; CAT#AM9763]) at 37°C for 25 min each, then twice in 2X SSC for 2 min each. Slides were incubated in DAPI nuclear stain for 15 min, washed in 1X PBS for 5 min, incubated with GFAP and histone cell segmentation markers for 1 hr, then washed three times in 1X PBS for 5 min each. Flow cells were affixed to each slide to create a fluidic channel for imaging, then loaded into the CosMx instrument. Approximately 300 FOVs were selected per slide, capturing hippocampal and cortical regions for each hemibrain section. Slides were imaged for 7 days and data were uploaded to the Nanostring AtoMx platform. Pre-processed data was exported as a Seurat object for further analysis in R 4.3.1.

#### Slide preparation for spatial proteomics

Slides were fixed in 10% NBF for 2 hr at 4°C, washed three times in 1X PBS for 5 min each, baked at 60°C for 30 min, then washed three times in 1X Tris Buffered Saline with Tween (TBS-T; CAT#J77500.K2) for 5 min each. Antigen retrieval was performed in a pressure cooker at 80°C in Tris-EDTA buffer (10 mM Tris Base [CAT#10708976001], 1 mM EDTA solution, 0.05% Tween 20, pH 9.0) for 7 min. After antigen retrieval, slides were cooled to RT for 5 minutes, then washed three times in 1X TBS-T for 5 min each. Slides were then incubated with Buffer W (Nanostring) for 1 hr at RT. Next, slides were incubated with CosMx 64-plex protein panel and segmentation markers (GFAP, IBA1, NEUN, and S6) at 4°C for 16-18 hr overnight. After overnight incubation, slides were washed three times in 1X TBS-T for 10 min each, then washed in 1X PBS for 2 min.

Fiducials for image alignment were diluted to 0.00005% in 1X TBS-T and applied to the slide, then incubated for 5 min. Slides were washed in 1X PBS for 5 min, incubated in 4% PFA for 15 min, then washed three times in 1X PBS for 5 min each. Slides were incubated in DAPI nuclear stain for 10 min, then washed twice in 1X PBS for 5 min each. Slides were incubated in 100 mM NHS-Acetate and washed in 1X PBS for 5 min. Flow cells were affixed to each slide, then loaded into the CosMx instrument. Approximately 700 FOVs were selected per slide, capturing each full hemibrain section. Slides were imaged for 6 days before the data were uploaded to the Nanostring AtoMx platform for analysis. Data visualization of each exported Seurat object file was performed using R 4.3.1.

#### Spatial transcriptomics data analysis

Spatial transcriptomics datasets were filtered using the AtoMx RNA Quality Control module to flag outlier negative probes (control probes targeting non-existent sequences to quantify non-specific hybridization), lowly-expressing cells, FOVs, and target genes. Datasets were then normalized and scaled using Seurat 5.0.1 *SCTransform* to account for differences in library size across cell types [26]. Principal component analysis (PCA) and uniform manifold approximation and projection (UMAP) analysis were performed to reduce dimensionality and visualize clusters in space. Unsupervised clustering at 1.0 resolution yielded 33 clusters for the 5xFAD dataset and 40 clusters for the PS19 dataset. Clusters were manually annotated based on gene expression and spatial location. Cell proportion plots were generated by first plotting the number of cells in each major cell type and scaling to 1. Normalized percentages for each genotype were calculated by dividing the number of cells in a given cell type-genotype pair by the total number of cells in that genotype, then dividing by the sum of the proportions across the cell type, to account for differences in genotype sample sizes (i.e., in the PS19;*ApoeCh* dataset, n=3/genotype except for *ApoeCh*, where n=2/ genotype). Differential gene expression analysis per cell type between genotypes was performed on scaled expression data using MAST [27] to calculate the average difference, defined as the difference in log-scaled average expression between the two groups for each major cell type. Differential upregulation (DU) and differential downregulation (DD) scores were calculated for all upregulated and downregulated genes, respectively, by summing the product of the negative log-10 adjusted p-value and the average difference for each statistically significant gene (i.e., p_adj_ < 0.05). A DU and DD score was calculated for each cluster, then plotted in XY space to visualize spatial differences in gene expression. Directionality of the DU or DD score is reported relative to the first group of the comparison; e.g., in the PS19;*ApoeCh* vs. PS19 comparison, a negative DD score indicates downregulation in the PS19;*ApoeCh* group. However, because PS19;*ApoeCh* brains have very few DAMs, visualizing DD score in PS19;*ApoeCh* brains does not clearly show the impact of *ApoeCh* on DAMs. To visualize DAMs in space, we instead plotted DD score in a PS19 brain (**Fig. 8i**). Cells with a negative DD score in a PS19;*ApoeCh* brain are conversely upregulated in a PS19 brain. Microglia, astrocyte, and oligodendrocyte glial populations were subsetted and further subclustered for deeper analysis. Data visualizations were generated using ggplot2 3.4.4 [28].

#### Spatial proteomics data analysis

Spatial proteomics data were filtered using the AtoMx Protein Quality Control module to flag unreliable cells based on segmented cell area, negative probe expression, and overly high/low protein expression. Mean fluorescence intensity data were hyperbolic arcsine transformed with the AtoMx Protein Normalization module. Cell types were automatically annotated based on marker gene expression using the CELESTA algorithm [29]. Cell proportions were calculated and percentages normalized in the same way as for spatial transcriptomics data. Protein expression was aggregated and scaled for each cell type to show overall differences in protein expression across the four genotypes. In the PS19;*ApoeCh* dataset, aggregate expression data from the *ApoeCh* genotype (n=2) was multiplied by 1.5 to normalize to the other genotypes (n=3). Data visualizations were generated using ggplot2 [28].

### Statistics

Every reported *n* represents the number of independent biological replicates. The sample sizes are similar to those found in prior studies conducted by MODEL-AD and were not predetermined using statistical methods [20]. Electrophysiology, immunohistochemical, and biochemical data were analyzed using Student’s t-test or two-way ANOVA via Prism v.9 (GraphPad, La Jolla, CA). Tukey’s post-hoc tests were utilized to examine biologically relevant interactions from the two-way ANOVA. Outlier tests were performed via Prism v.9 where relevant and any datapoints removed from the analyses acknowledged in the relevant figure legend. \**p* ≤ 0.05, \*\**p* ≤ 0.01, \*\*\**p* ≤ 0.001, \*\*\*\**p* ≤ 0.0001. Statistical trends were accepted at *p* < 0.10 (# denotes trending significance). Data are presented as raw means and standard error of the mean (SEM).

## RESULTS

### Generation of ApoeCh mice

The three common human isoforms of APOE (e2, e3, e4) are distinguished by the presence of either arginine (R) or cysteine (C) at positions 112 and 158. In APOE4, which is now considered causative for AD in a homozygous state [30], arginine residues are found at both sites. Although mice only express one common form of APOE, murine APOE (mAPOE) also contains arginine at positions 112 and 158. However, mAPOE and hAPOE show only partial homology between these sites (**Fig. 1a**), suggesting there could be differences in how this region of APOE functions between the species. Notably, a HSPG and LDLR binding site in the APOE sequence is conserved in mice (blue boxed area), including arginine 136, which is replaced by serine in the Christchurch (Ch) variant. Given that APOE is a ligand for HSPG and LDLR and is involved in Aβ aggregation and tau uptake, we investigated the effect of introducing the APOE Ch variant (R136S) into mAPOE on development of AD-like pathology, independent of hAPOE4. To accomplish this, we generated *ApoeCh* mice by utilizing CRISPR Cas-9 to introduce an R->S substitution at amino acid 128 in mAPOE, corresponding to R136S in hAPOE (**Fig. 1a**). DNA sequence was verified by sequencing of the targeted region (**Supplemental Fig. 1a-b**).

**Figure 1:**
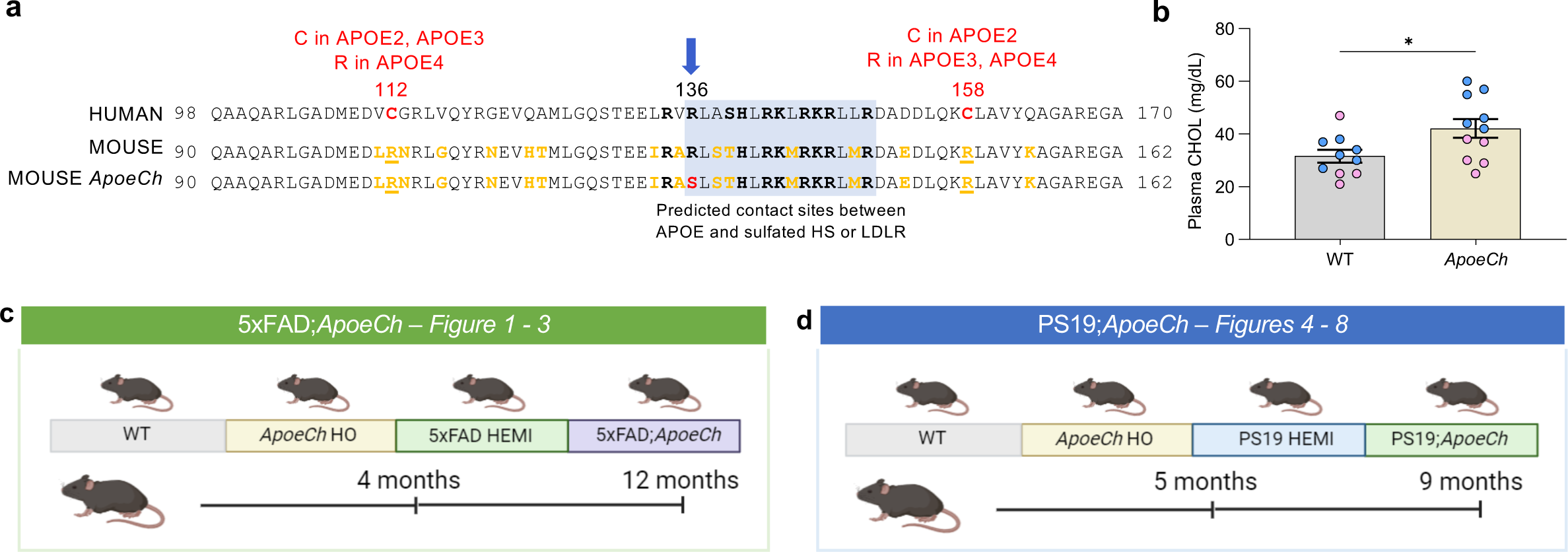
Generation of *ApoeCh* mouse model. **a** Partial amino acid (AA) sequence alignment between human and mouse APOE. The vertical blue arrow denotes the location of the R136S Christchurch variant (*rs121918393*), shown in red in the mouse *ApoeCh* sequence. Mouse has human APOE4 type sequence at positions 112 and 158. The AA differences between mouse and human APOE4 in this region are highlighted in yellow. **b** Plasma cholesterol in 4 mo WT and *ApoeCh* HO mice (p=0.0274). **c, d** Schematic showing mouse groups and study design of 5xFAD;*ApoeCh* (**c**) and PS19;*ApoeCh* (**d**) cohorts.

We conducted an off-target analysis of other likely CRISPR cut sites that might have been targeted during the generation of the mAPOE R136S allele. *ApoeCh* CRISPR G0 founder mice were backcrossed with wild-type B6J animals for three generations before being used to generate homozygous animals for this study, making it unlikely that a mutation caused by an off-target effect of CRISPR/Cas9 would be present on a chromosome other than chromosome 7, i.e., the location of *Apoe* (B6J; Chr7; 48.6 Mb, GRCm39, Ensembl release 108). Potential CRISPR/Cas9 off-target sites with up to four mismatches using crRNA TMF1648 were screened for using Cas-OFFinder (http://www.rgenome.net/cas-offinder/; [31]). Twelve potential off-target sites on mouse chromosome 7 were identified (**Supplemental Table 1**). Two potential off-target sites were within loci encoding lncRNAs, three were within introns, while one was in a 5‘UTR, with the remaining six located within intergenic regions. To screen for evidence of CRISPR/Cas9 RNP activity at non-intergenic sites, DNA from a wild-type and homozygous *ApoeCh* mouse was amplified by PCR using the primers listed (**Supplementary Table 2**) then sequenced across the potential off-target region at six of the loci (**Supplemental Fig. 1d-j**). None of the potential off-target sites showed a difference in sequence between WT and homozygous *ApoeCh* mice.

As with the human carrier of PSEN1*E280A who was homozygous for the APOE3Ch variant, and mice with a humanized APOE3*R136S allele, *ApoeCh* mice exhibited elevated plasma cholesterol, a phenomenon also observed in approximately 10% of APOE2 homozygotes [12, 32]. However, we observed no change in plasma triglyceride in our mouse model (data not shown) (**Fig. 1b**).

To investigate whether introduction of the Christchurch variant into mAPOE affects development of AD-like pathologies, while distinguishing between amyloid- and tau-dependent effects, we independently crossed *ApoeCh* mice with the 5xFAD amyloidosis mouse model [20, 33, 34] (**Fig. 1c**), and the PS19 tauopathy mouse model [35–37] (**Fig. 1d; data from Fig. 4-7**). For crosses with the 5xFAD mouse, we generated four groups: (i) WT, (ii) *ApoeCh* homozygotes (*ApoeCh*), (iii) 5xFAD hemizygotes (5xFAD) and (iv) 5xFAD; *ApoeCh*, (5xFAD;*ApoeCh*). Similarly, we generated 4 groups for the tauopathy model: (i) WT, (ii) *ApoeCh,* (iii) PS19 hemizygous (PS19), and (iv) *ApoeCh*; PS19 (PS19;*Apoe*Ch). To capture age- and disease progression-dependent changes in phenotypes, we investigated animals at 4 and 12 mo old for 5xFAD cohort and 5 and 9 mo old for the PS19 cohort, corresponding to early and late stages of disease, respectively (**Fig. 1c, d**).

### ApoeCh variant reduces dense-core Aβ plaque load and plaque-induced neuronal damage

In 5xFAD; *ApoeCh* cohort animals, we observed no significant change in body weight between genotypes at 4 mo, and a small reduction in 5xFAD;*ApoeCh* compared to 5xFAD at 12 mo (**Supplemental Fig. 2a, b**). Open field analyses revealed no consistent change in motor function between genotypes at either age (**Supplemental Fig. 2c-f**). 5xFAD mice show atypical behavior in the elevated plus maze test, with aged 5xFAD mice preferring the open arms over the closed arms, compared to WT mice. No change was detected in elevated plus maze in the 4-mo groups between any genotype, when pathology is in its early stages in 5xFAD mice. At 12 mo, both 5xFAD and 5xFAD;*ApoeCh* mice spent more time in the open arms compared to controls although no significant difference was found between 5xFAD and 5xFAD;*ApoeCh* mice (**Supplemental Fig. 2g, h**).

After behavioral testing, brains were extracted and prepared for histological, biochemical, and transcriptomic analyses. AmyloGlo staining revealed a reduced brain-wide dense core plaque load in 5xFAD;*ApoeCh* compared to 5xFAD mice at 4 and 12 mo of age, with reduced plaque number at 12 mo in males (**Fig. 2a-e**). The overall decrease in plaque load was accompanied by increased soluble and insoluble Aβ40 and Aβ42 in the cortex and hippocampus via MULTI-ARRAY assay (Meso Scale Discovery) at 4 mo of age in 5xFAD;*ApoeCh* compared to 5xFAD mice, which is partially resolved by 12 mo (**Fig. 2f-m**). Plaque-induced neuritic damage was assessed by staining for lysosomal-associated membrane protein 1 (LAMP1), which accumulates in dystrophic neurites around plaques [38]. Reduced LAMP1 volume was observed in 5xFAD;*ApoeCh* compared to 5xFAD mice at both time points, which is consistent with lowered amyloid plaque load (**Fig. 2n-p**). Plasma neurofilament light chain (NfL), an established marker for axonal degeneration that is highly correlated with AD progression [39], was elevated in 5xFAD mice at both 4 and 12 mo compared to WT mice. 5xFAD mice with *ApoeCh* mutation exhibited lower plasma NfL compared to their 5xFAD age-matched controls (**Fig. 2q-r**), suggesting a protective effect of *ApoeCh* on neurodegenerative processes. Together, these results show that the presence of the *ApoeCh* variant in the 5xFAD mice reduces plaque pathology as well as amyloid-associated downstream damage.

**Figure 2:**
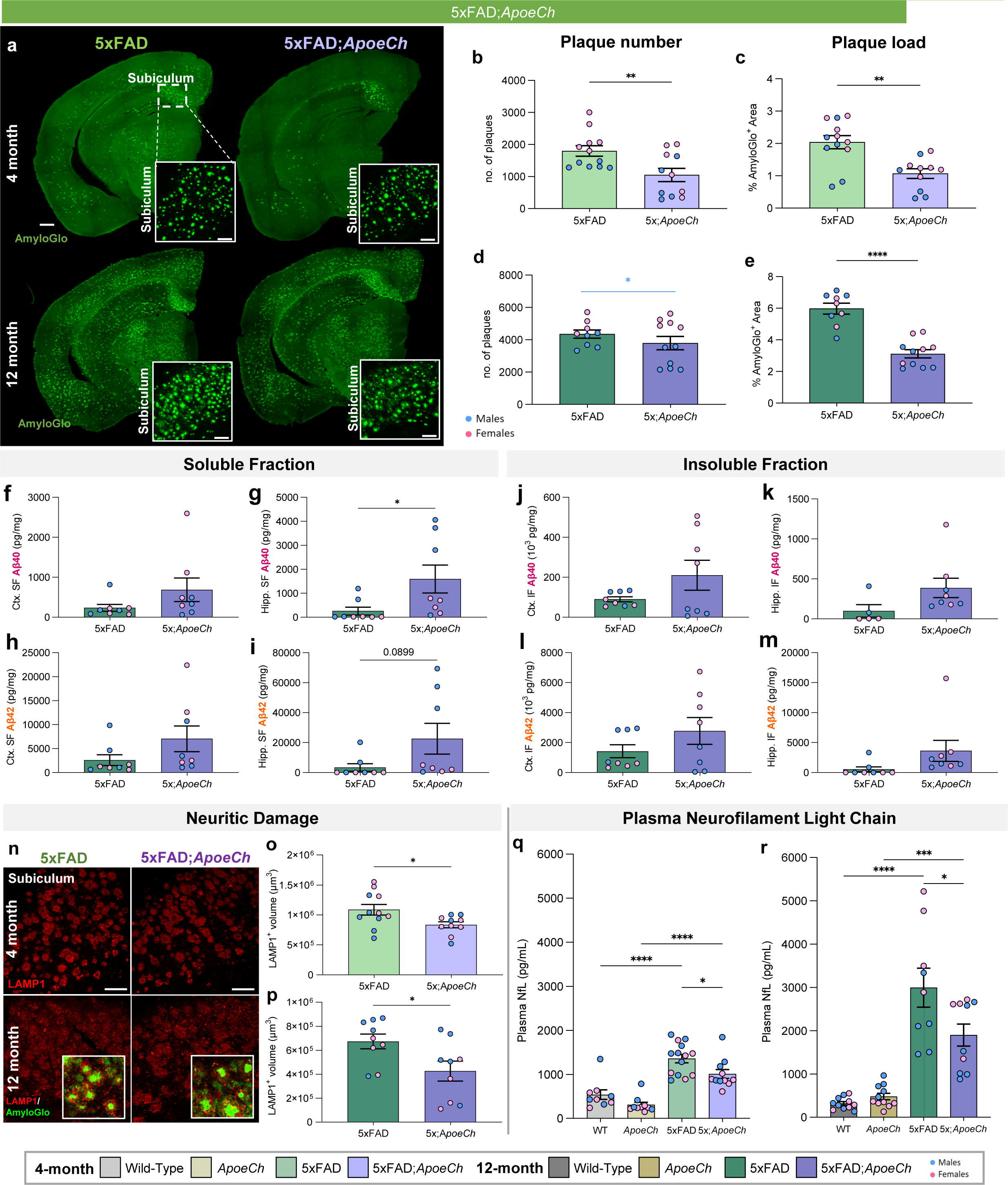
*ApoeCh* variant ameliorates Aβ plaque burden and plaque-induced damage in 5xFAD mice. **a** Representative hemispheric coronal brain images of 4-mo-old (top) and 12-mo-old (bottom) 5xFAD and 5xFAD;*ApoeCh* stained for dense-core plaques using AmyloGlo (green) Scale bar = 500µm. Insets of 20x magnification images of the subiculum. Scale bar = 100µm. **b-e** Quantification of AmyloGlo^+^ plaques (4 month - **b**; 12 month - **d**) and percent AmyloGlo^+^ area coverage (4 month - **d**; 12 month - **e**) of 5xFAD and 5xFAD;*ApoeCh* mice. Blue bar denotes statistical significance only in males. **f-m** Quantification of soluble (**f-k**) and insoluble (**j-m**) Aβ in micro-dissected cortices (**f, g, j, k**) and hippocampi (**h, i, l, m**) of 12 month-old 5xFAD and 5xFAD;*ApoeCh* mice. **n** Representative confocal images of subiculum in 4-mo-old (top) and 12-mon-old (bottom) wild-type, *ApoeCh*, 5xFAD, and 5xFAD;*ApoeCh* mice immunolabeled for LAMP1 (red) for dystrophic neurites. Insets show higher magnification images with LAMP1 (red) and AmyloGlo for dense-core plaques (green). Scale bar = 100µm. Student’s t-test. **q-r** Measurement of plasma NfL in WT, *ApoeCh*, 5xFAD, and 5xFAD;*ApoeCh* mice at 4-(**q**) and 12- mo (**r**). n=4-6 mice/sex/genotype. Data are represented as mean ± SEM. Two-way ANOVA followed by Tukey’s post hoc tests to examine biologically relevant interactions. Statistical significance is denoted by *p<0.05, **p<0.01, ***p<0.001, ****p<0.0001.

### Spatial proteomics reveal increased disease associated microglia in response to plaques with ApoeCh

As APOE proteins are produced primarily by astrocytes and reactive microglia in the CNS, we next examined *ApoeCh*-associated changes in glial cells by histology. Reactive astrocytes, detected by glial fibrillary acidic protein (GFAP), were increased in the subiculum of both 4- and 12-mo-old 5xFAD mice compared to WT (**Fig. 3a-ba; Supplemental Fig. 4a**). Consistent with lower plaque load, there was decreased astrocyte volume in 5xFAD;*ApoeCh* mice compared to 5xFAD at both ages (**Fig. 3a-b; Supplemental Fig. 4b**). Microglia were detected through immunostaining for ionized calcium binding adaptor molecule 1 (IBA1; **Fig. 3c; Supplemental Fig. 4c**) and increased IBA1+ staining was detected in the subiculum of 5xFAD mice at both 4- and 12-mo of age, where abundant plaque load develops, compared to WT. Notably, concomitant with lower plaque load, overall microglial load was significantly reduced at 12 mo in 5xFAD;*ApoeCh* mice compared to age-matched 5xFAD mice in both total IBA1+ volume (**Fig. 3d; Supplemental Fig. 4d**) or IBA1+ cell counts (**Supplemental Fig. 4f**).

**Figure 3:**
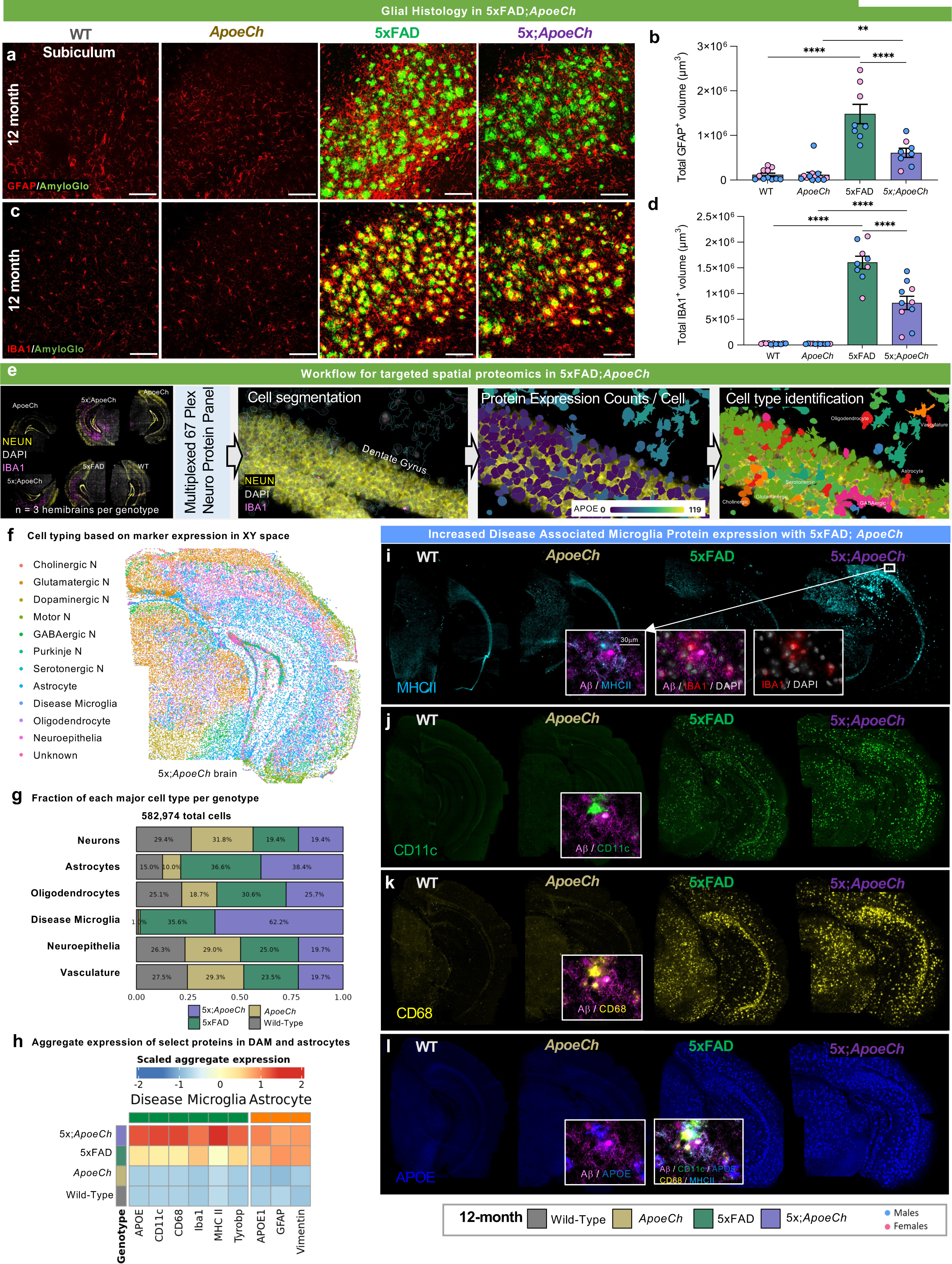
*ApoeCh* increases disease-associated microglia number in response to plaques. **a** Representative confocal images of the subiculum stained for dense-core plaques with AmyloGlo (green) and immunolabeled for GFAP (red, **a**) and IBA1 (red, **c**) of 12-mo-old WT, *ApoeCh*, 5xFAD, and 5xFAD;*ApoeCh* mice. Scale bar=100µm. **b, d** Quantification of total volume of GFAP^+^ cells (**b**) and IBA1^+^ cells (**d**). n=4-6 mice/sex/genotype. Data are represented as mean ± SEM. Two-way ANOVA followed by Tukey’s post hoc tests to examine biologically relevant interactions. Statistical significance is denoted by *p<0.05, **p<0.01, ***p<0.001, ****p<0.0001.**e** Workflow for targeted 67-plex single-cell spatial proteomics. Fields-of-view (FOVs) are first imaged with GFAP, NEUN, RPS6, and IBA1 markers for cell segmentation. Protein abundance is determined by counting the number of fluorescently-labelled oligos in each cell. Cell types are identified with the CELESTA algorithm, which classifies cells based on marker protein expression. **f** Cell types in XY space. CELESTA classifies cells into 12 different cell types, which can then be plotted in space to confirm accurate identification. Non-DAM microglia are unable to be identified using CD11b as a marker; only DAM are shown. **g** Proportions of 5xFAD;*ApoeCh*, 5xFAD, *ApoeCh*, and WT cells for each major cell type. **h** Aggregate expression of the top differentially expressed proteins in DAMs and astrocytes across the four genotypes. **i-l** Immunofluorescence images of MHCII, CD11c, CD68, and APOE for representative brains of WT, *ApoeCh*, 5xFAD, and 5xFAD;*ApoeCh* mice.

To study the effects of *ApoeCh* on the brain’s response to plaques, we next employed spatial proteomics utilizing a multi-plex 67-protein neuroscience mouse panel (Nanostring CosMx spatial molecular imager; **Figure 3e**). This technique maintains the original structure of brain tissue and allows for detailed analysis based on protein markers. The panel includes numerous markers for reactive microglia and astrocytes, enabling evaluation of their response to plaques. For each of the four genotypes, data was acquired from three 10 µm coronal slices from 14-mo-old hemibrains, yielding a dataset of 582,974 cells. Cell segmentation was automated using histone, GFAP, and DAPI markers, and cells were categorized by protein expression (**Fig. 3e**). Examples of cell segmentation are shown in **Supplemental Fig. 5a**, illustrating effective cellular identification across dense neuronal areas such as CA1 and dentate gyrus. Plotting and visualization of these cells in XY space confirmed accurate cell identification (**Fig. 3f**; **Supplemental Fig. 5b**). Using this approach, we identified astrocytes, microglia, oligodendrocytes, neuroepithelial cells, and seven neuronal subtypes. We were able to identify disease-associated microglia (DAM) based on protein expression of CD11c, but not homeostatic microglia. Quantification of cell proportion normalized by total number of cells per genotype revealed a significant increase in the number of DAMs in 5xFAD;*ApoeCh* (∼62%) compared to 5xFAD (∼36%) mice (**Fig. 3g; Supplemental Fig. 5c**), despite the reduction in plaque load and reduced overall microglial volume (**Fig. 2a-d**, **Fig. 3c-d**). Unlike DAM, the number of astrocytes was similar in both 5xFAD and 5xFAD;*ApoeCh* groups compared to WT and *ApoeCh* mice, respectively. We detected no change in neuronal, neuroepithelial, or vascular populations between WT and 5xFAD mice or 5xFAD and 5xFAD;*ApoeCh* mice (**Fig. 2g)**. These data indicate that introduction of the *ApoeCh* variant into 5xFAD mice increases the proportion of microglia that are identified as DAM.

To explore the microglial response to plaques, we next visualized and measured intensity of several of the DAM markers included in the panel, including MHCII, CD11c, CD68, and APOE (**Fig. 3i-l**; inserts show localization of these markers around plaques), and found that each marker was upregulated in microglia surrounding plaques. Aggregated expression of these proteins was plotted as a heatmap across the four genotypes, as well as IBA1 and TYROBP, showing significantly increased protein expression of microglial and DAM proteins in 5xFAD;*ApoeCh* compared to 5xFAD brains (**Fig. 2h**). Plotting protein levels of APOE, GFAP, and VIM in astrocytes shows increases in both 5xFAD genotypes but no difference between 5xFAD;*ApoeCh* and 5xFAD brains (**Fig. 2h**). Taken together, these data provide evidence that despite reducing plaque load and overall microglial numbers, the presence of *ApoeCh* robustly increases the proportion of DAMs in response to amyloid plaques, as well as upregulates DAM protein expression levels.

### Exploration of the CNS response to amyloid pathology using spatial transcriptomics

To understand the effects of *ApoeCh* on the brain’s response to plaques, we investigated transcriptional changes in microglia and other CNS cell types using spatial transcriptomics. This method enables single-cell level gene expression analysis in microglia, overcoming limitations associated with single-cell and nucleus RNA-seq, such as technique-induced expression changes [40–44]. Additionally, it provides information on genes at distinct spatial locations within the tissue, facilitating *in situ* counting and profiling of all cell populations within a brain slice. We conducted our study with single-cell resolution using multiplexed error-robust fluorescence *in situ* hybridization (MERFISH; Nanostring CosMx spatial molecular imager) and employed a 1000-plex RNA mouse neuroscience panel (Nanostring). Three coronal hemibrain slices per genotype (adjacent to the sections used for spatial proteomics) were processed capturing the hippocampus and cortical regions, resulting in a dataset encompassing 425,663 total cells from 12 brains. Cell segmentation was performed based on histone and ribosome staining (sample cell segmentation examples shown in **Supplemental Fig. 6a**), and transcript counts per cell for each of the 1000 genes were then calculated with an average of 754 transcripts per cell for a total of 426,231 cells (workflow shown in **Fig. 4a**). Cell clustering was performed using a community detection approach on a k-nearest neighbor graph, followed by dimensionality reduction via Uniform Manifold Approximation and Projection (UMAP). Clusters were manually annotated based on gene expression and spatial location. Thirty-three total clusters were identified corresponding to 11 clusters of excitatory neurons, 6 clusters of inhibitory neurons, 3 astrocyte clusters, 6 oligodendrocyte and oligodendrocyte precursor clusters, 2 microglial clusters, as well as 5 endothelial and vascular-related clusters (**Fig. 4b**). As cell clustering relies purely on gene expression and not spatial coordinates, we next verified that cell clustering correctly identified anatomical locations of the cell populations by plotting cells in XY space (**Fig. 4c**, with all 12 brains shown in **Supplemental Fig. 6b**). Cell counts for each of the 33 clusters per genotype were calculated (**Supplemental Fig. 7c**), then condensed into major clusters encompassing similar cell types (e.g., ODC1-5 were combined into a broader “oligodendrocyte” cluster). Top gene expression per major cluster are shown in **Supplemental Fig. 7b**. The proportion of cells in each major cluster were then plotted per genotype (**Fig. 4d**). We observe an increased proportion of cells in the disease associated astrocyte (DAA) and disease-associated microglia (DAM) clusters in both 5xFAD and 5xFAD;*ApoeCh* mice, although no change was found among the other cell types. Concordant with spatial proteomics, we observe increases in DAMs in 5xFAD;*ApoeCh* compared to 5xFAD mice, despite their reduced plaque load (**Fig. 4d**). Plotting the DAM cluster in XY space reveals that DAMs are distributed in 5xFAD and 5xFAD;*ApoeCh* slices in plaque-laden areas (i.e., the subiculum of the hippocampus, throughout the cortex; **Fig. 4e**). Visualization of cellular expression of the DAM gene *Cst7* (**Fig. 4f**) shows that *Cst7*-expressing cells cluster around plaques (yellow-green *Cst7* expressing cells surround a white plaque; **Fig. 4g**), as expected.

**Figure 4:**
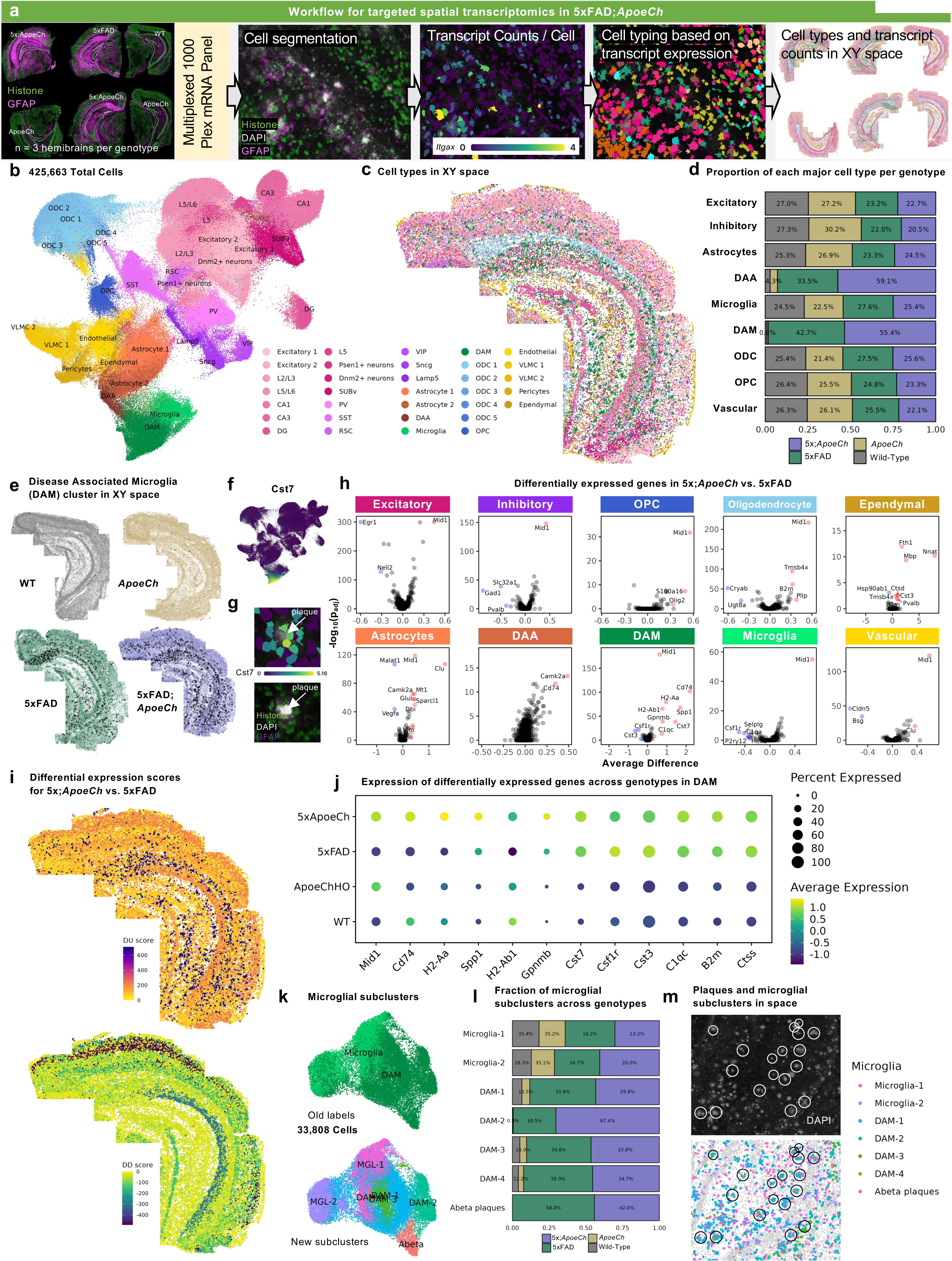
*ApoeCh* enhances microglial response to plaques confirmed by spatial transcriptomics. **a** Workflow for targeted 1000-plex single-cell spatial transcriptomics. FOVs were selected in hippocampus and cortex of each section, then imaged with DNA, rRNA, Histone, and GFAP markers for cell segmentation. Transcript counts for each gene were acquired per cell. **b** UMAP of 425,663 cells across 12 hemibrains (n=3/genotype). Clustering at 1.0 resolution yielded 33 clusters, which were annotated manually based on gene expression and anatomical location in space. **c** 33 clusters plotted in XY space. **d** Proportion of the number of cells in each major cell type, grouped by genotype. Percentages normalized for the total number of cells in each genotype. **e** DAM cluster (black dots) plotted in XY space for each genotype. **f** Feature plot of *Cst7* expression in UMAP space. **g** (top) *Cst7*-expressing cells (i.e., DAMs) surrounding an amyloid-beta plaque. (bottom) DAPI (grey) highlights the same plaque from top panel. Histone marker (green) highlights nucleosomes in cells surrounding the plaque. GFAP+ (purple) cell processes surround the plaque. **h** Volcano plots showing DEGs between 5xFAD;*ApoeCh* and 5xFAD among major cell types. **i** Differential upregulation (DU) and differential downregulation (DD) scores for 5xFAD;*ApoeCh* vs. 5xFAD in each cluster plotted in space in a 5xFAD;*ApoeCh* brain. **j** Pseudo-bulk expression of top DEGs between 5xFAD;*ApoeCh* and 5xFAD across genotypes. **k-m** Microglial subclustering analysis. **k** UMAP of 33,808 subsetted microglia. (top) UMAP annotated with old labels from **b**. (bottom) Cells were re-clustered at 0.2 resolution to yield 7 new subclusters. **l** Proportion of the number of cells in each microglial subcluster, grouped by genotype. **m** Microglial subclusters in XY space in a 5xFAD;*ApoeCh* brain. (top) Amyloid-beta dense core plaques, shown by the presence of DAPI+ aggregates (grey), circled in white. (bottom) Microglial subclusters plotted in space. Plaques from top panel circled in black.

To understand the impact of plaque deposition on the different cell types of the CNS, we next conducted differential gene expression (DGE) analysis on 5xFAD and WT brains. Volcano plots show changes in gene expression within each cell population between these two groups (**Supplemental Fig. 8a, c)**. Consistent with previous sequencing studies, all microglia (combined DAM and microglia clusters, as there are no DAMs in WT and *ApoeCh* genotypes) appear strongly impacted by plaques, as seen by the highest number of DEGs (with 48 DEGs, defined as p_adj_ < 0.05 and absolute average difference > 0.3) between 5xFAD and WT mice. In line with other studies, we observe an upregulation in DAM genes (i.e., *Cst7*, *Apoe*, *Ctsb*, *Ctsd*, *Ctss*, and *Psen1*) and downregulation in homeostatic genes (*P2ry12*, *Tmem119*, and *Csf1r)* in 5xFAD compared to WT mice [45–47]. In exploring the effects of plaques on neuronal-related genes, we see consistent downregulation of *Egr1*, involved in learning and memory, in 5xFAD compared to WT mice [48]. Our analysis does not detect many genetic changes in grouped excitatory or inhibitory neuronal clusters (4 DEGs in excitatory and 4 in inhibitory; **Supplemental Fig. 8c**) but does identify several larger genetic changes when examined in specific neuronal populations (i.e., 23 DEGs in CA1 and 13 DEGs in Subiculum (SUBv)); **Supplemental Fig. 8a**). These data illustrate the importance of performing non-aggregate DGE analyses, especially while examining genetic changes within neuronal populations.

Next, we investigated how the presence of *ApoeCh* impacted the brains’ response to plaques. Non-aggregate DGE analysis of 5xFAD;*ApoeCh* vs. 5xFAD brains was performed and revealed changes in gene expression across many of the 33 cell clusters (**Supplemental Fig. 8b**), as well as in the aggregate major cell clusters (**Fig. 4h**), particularly DAM and DAA. To visualize these genetic changes (across all cell types) with spatial resolution, we first generated a DEG score, which represents the magnitude of genetic change between any two animal groups for each cell cluster. DEG score was calculated by summing the product of the average difference in gene expression between two genotypes and the negative log 10 p-value for each gene. The calculation gave each cluster a value which is then plotted in XY space (DU score represents the DEG score of all upregulated genes and DD score represents the DEG score of all downregulated genes; **Fig. 4i**). Focusing on downregulated genes, visualization of the DD score implicates regional specificity of genetic changes induced by introduction of *ApoeCh* into 5xFAD mice within L2/L3 cortical and CA1 neurons (**Fig. 4i**). Replotting of DG, CA1, and L2/L3 clusters in XY space confirmed correct cell type clustering with high spatial specificity (**Supplemental Fig. 9a**). To visualize gross genetic changes across the different genotypes, we next plotted the DEGs from each of these clusters as pseudo-bulked expression values across the four animal groups (**Supplemental Fig. 9b**). For L2/L3, several genes were upregulated in 5xFAD mice relative to WT, and then downregulated with 5xFAD;*ApoeCh* (i.e., *Malat1*, *Olfm1*, *Nell2*, and *Snap25*). Many of these genes are involved in cell migration, and axon growth and guidance [49–52]. For CA1, most of the changes appeared in *ApoeCh* mice regardless of 5xFAD transgene array, with similar effects seen in both *ApoeCh* and 5xFAD;*ApoeCh* brains (i.e., *Nell2*, *Penk*, *Mid1*, *Cck*, *Dnm1*, *Nnat*, *Gap43*, and *Atp2b1*). Several of these genes implicate *ApoeCh* in modulating synaptic function and plasticity.

### Spatial transcriptomics confirms increased disease-associated gene expression in microglia in response to plaques with ApoeCh

APOE is involved in the regulation of DAMs and the microglial response to amyloid [46, 53–55]. Among all the cell types, microglia surrounding plaques (i.e., DAMs) exhibited the greatest differential upregulation (DU) score between 5xFAD;*ApoeCh* and 5xFAD, implicating an important role of *ApoeCh* in regulating the DAM response to plaques (**Fig. 4i**). Consistent with the protein expression data, DAM genes were further upregulated in DAMs in 5xFAD:*ApoeCh* (i.e., *Cd74*, *Cst7*, *Gpnmb*, *Spp1*, *H2-Aa*, and *H2-Ab1;* **Fig. 4h**) while homeostatic microglial genes were further downregulated (i.e., *Csf1r*, *Cst3*). Pseudo-bulk analysis of top DEGs between 5xFAD;*ApoeCh* and 5xFAD in DAMs was performed across the four genotypes, highlighting again that upregulated genes in 5xFAD relative to WT microglia are further upregulated with *ApoeCh* (**Fig. 4j**). To gain a deeper understanding of microglial populations, we captured all 33,808 microglia and DAM cells across the four groups and reclustered them into seven new subclusters (**Fig. 4k**). Among the subclustered microglial cells, we identified a subcluster that appeared to represent the epicenter or core of Aβ dense core plaques, as validated by aligning the coordinates of this subcluster with DAPI-positive plaques from CosMx cell segmentation imaging (**Fig. 4m**). We demonstrated that many “cells” in this cluster were anucleate and did not express histone markers (**Supplemental Fig. 10b**). As expected, these presumed plaque core subclusters were only present in 5xFAD;*ApoeCh* and 5xFAD genotypes (**Fig. 4l**). Compared to other microglial subclusters, this Aβ plaque core subcluster contained a high level of *Iapp*, which encodes the amyloidogenic hormone amylin, along with RNAs for genes involved in microglial activation (i.e., *Cast*, *Pde7b*, *Ltbp1;* **Supplemental Fig. 10c**).

Surrounding these plaque core subclusters in space were subclusters DAM-1 and DAM-2, representing plaque-associated microglia. DAM-1 is characterized by high expression of genes involved in antigen presentation and the complement system (i.e., *Ctss*, *Ctsb*, *Ctsd*, *C1qa*, *C1qb*). By contrast, DAM-2 shows high expression of classic DAM marker genes such as *Cst7*, *Trem2*, *Cd74*, and *Itgax* (**Supplemental Fig. 10c**). Cell proportions were calculated for each microglial subcluster across the four genotypes (**Fig. 4l**). While there was no difference in cell proportions of homeostatic microglia between 5xFAD;*ApoeCh* and 5xFAD, the number of cells (i.e., plaques) in the Aβ plaque core subcluster was reduced in 5xFAD;*ApoeCh* compared to 5xFAD, in line with our IHC findings that the presence of *ApoeCh* reduces plaque load in 5xFAD mice. Furthermore, while numbers of DAM-1 were unchanged between 5xFAD;*ApoeCh* and 5xFAD, DAM-2 (**Fig. 4l**) was significantly increased in 5xFAD;*ApoeCh* compared to 5xFAD, supporting the increased DAM response in 5xFAD;*ApoeCh* seen in our proteomics data. Together, these data show that the presence of *ApoeCh* in 5xFAD mice upregulates the response of a *Cst7*-expressing DAM population around plaques and reduces plaque load.

#### ApoeCh has minimal effects on phosphorylated tau accumulation in the PS19 model

Having shown that the presence of *ApoeCh* promotes a specific DAM response that is neuroprotective against plaque pathology, we next investigated the effect of *ApoeCh* on development of tau pathology. To do so, we crossed *ApoeCh* mice with the PS19 model of tauopathy along with additional crosses for controls, generating four animal groups: (i) WT, (ii) *ApoeCh* HO (*ApoeCh*), (iii) PS19 hemizygous (PS19), and (iv) *ApoeCh* HO; PS19 (PS19;*Apoe*Ch) and aged two cohorts to 5 or 9 mo of age (**Fig. 1d**). As anticipated, weight loss was observed in both PS19 and *ApoeCh*;PS19 mice at 9 mo of age while no difference was observed between *ApoeCh* and WT mice (**Supplemental Fig. 11a, b**). Scoring of hindlimb clasping reflex, a marker of neurodegenerative disease progression, was assessed with impairment recorded in both PS19 and PS19;*ApoeCh* mice at 9 mo of age (**Supplemental Fig. 11c, d**) [21]. Open field analyses showed no difference between genotypes at either 5 or 9 mo (**Supplemental Fig. 11e-h**), but both PS19 and PS19;*ApoeCh* mice exhibited impairment during elevated plus maze testing at both ages (**Supplemental Fig. 11i, j**). These data show that the *ApoeCh* variant was unable to prevent development of motor and anxiety-related impairments in PS19 mice, based on the assays used.

Following completion of behavioral testing, brains were extracted and processed for histology, biochemistry, and transcriptomic analyses. Immunostaining for AT8, which detects the phosphorylation of serine 202 threonine 205 of tau, revealed prominent AT8+ pathology in the dentate gyrus (DG) and piriform cortex (PIRI) of in the PS19 mouse model. However, no significant difference was observed between PS19 and PS19;*ApoeCh* mice at both 5 and 9 mo of age (**Fig. 5a-d, Supplemental Fig. 12a-c**) [35, 56–58].

**Figure 5:**
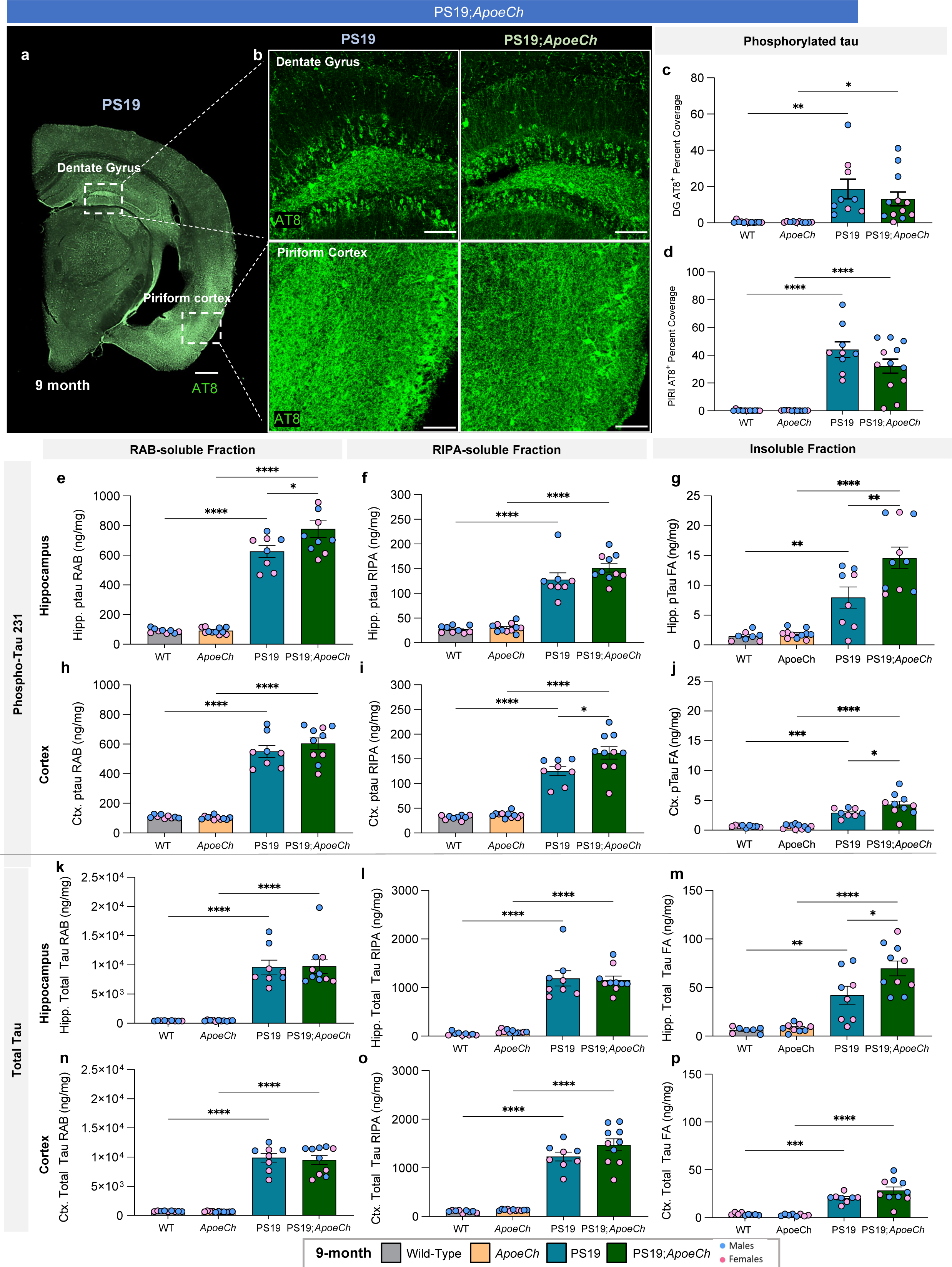
*ApoeCh* has minimal impact on phosphorylated tau accumulation. **a** Representative 10x image of hemispheric coronal brain image of a PS19 mouse immunolabeled for phosphorylated tau using AT8 (green) highlighting brain regions with high pathological manifestation, dentate gyrus (DG) and piriform cortex (PIRI). Scale bar=500µm **b** Representative 20x confocal images of dentate gyrus (top) and piriform cortex (bottom) from 9-mo old PS19 and PS19;*ApoeCh* mice immunolabeled for AT8 (green). Scale bar=100µm. **c-d** Percentage of AT8^+^ area coverage per FOVs of 9 mo-old WT, *ApoeCh*, PS19, and PS19;*ApoeCh* mice in the DG (**c**) and PIRI (**d**). **e-j** Phosphorylated tau T231 measured via MSD in RAB, RIPA, and formic acid fraction of the micro-dissected hippocampi (**e-g**) and cortices (**h-j**) of 9-mo-old WT, *ApoeCh*, PS19, and PS19;*ApoeCh* mice. **k-p** Measurement of total tau in RAB, RIPA, and formic acid fraction of the micro-dissected hippocampi (**k-m**) and cortices (**k-p**) of 9-mo-old WT, *ApoeCh*, PS19, and PS19;*ApoeCh* mice. n=3-6 mice/sex/genotype. Data are represented as mean ± SEM. Two-way ANOVA followed by Tukey’s post hoc tests to examine biologically relevant interactions. Statistical significance is denoted by *p<0.05, **p<0.01, ***p<0.001, ****p<0.0001.

Sequential biochemical extractions in high salt reassembly buffer (RAB), lysis radioimmunoprecipitation assay buffer (RIPA), and 70% formic acid (FA) were performed to obtain extracellular soluble, intracellular detergent soluble, and highly insoluble tau proteins, respectively. Utilizing these brain lysates, we quantified total and phosphorylated tau (Thr231) in 9-mo-old mice via MULTI-ARRAY assay (Meso Scale Discovery). Phospho-tau 231 was increased in all three fractions in both the hippocampus and cortex of PS19;*ApoeCh* mice compared to PS19 mice (**Fig. 5e-j**). Total tau was increased in the insoluble hippocampal fraction PS19;*ApoeCh* mice compared to PS19 mice. No significant change was detected between these mice in the cortex or other lysate fractions (**Fig. 5k-p**). Together, these data indicate that the presence of *ApoeCh* increases the phosphorylation of tau, at Thr231 but does not profoundly alter the accumulation of total tau in PS19 mice.

We next measured plasma NfL, as a surrogate for axonal damage, and as expected found a dramatic elevation in NfL in PS19 compared to WT mice at 9 mo of age, although no difference was found between PS19 and PS19;*ApoeCh* mice (**Supplemental Fig. 11k, l**).

### ApoeCh suppresses the glial response to tauopathy

Since the introduction of *ApoeCh* into 5xFAD mice promoted the microglial response to plaques, we next explored if this variant would have similar effects on the glial response to tauopathy. We focused our analyses on the 9-mo cohort as no overt pathology was seen at 5-mo (**Supplemental Fig. 12a-h**). Immunohistology of the dentate gyrus revealed a dramatic increase in tau-induced astrocytic reactivity visualized by GFAP in PS19 mice that was suppressed in PS19;*ApoeCh* mice (**Fig. 6a, b**). Similarly, immunostaining for microglia revealed increased total volume of IBA1+ microglia in PS19 mice relative to WT. In PS19;*ApoeCh* mice, IBA1+ microglia total volume was decreased, despite no change in overall microglia number (**Fig. 6c, d, Supplemental Fig. 12i**), suggesting an altered microglial response to the pathology. Thus, despite increases in phosphorylated tau T231, we observed a robust decrease in tau-induced glial responses in the PS19;*ApoeCh* mice. This contrasts with the effects observed of *ApoeCh* on microglia in the presence of plaques.

**Figure 6:**
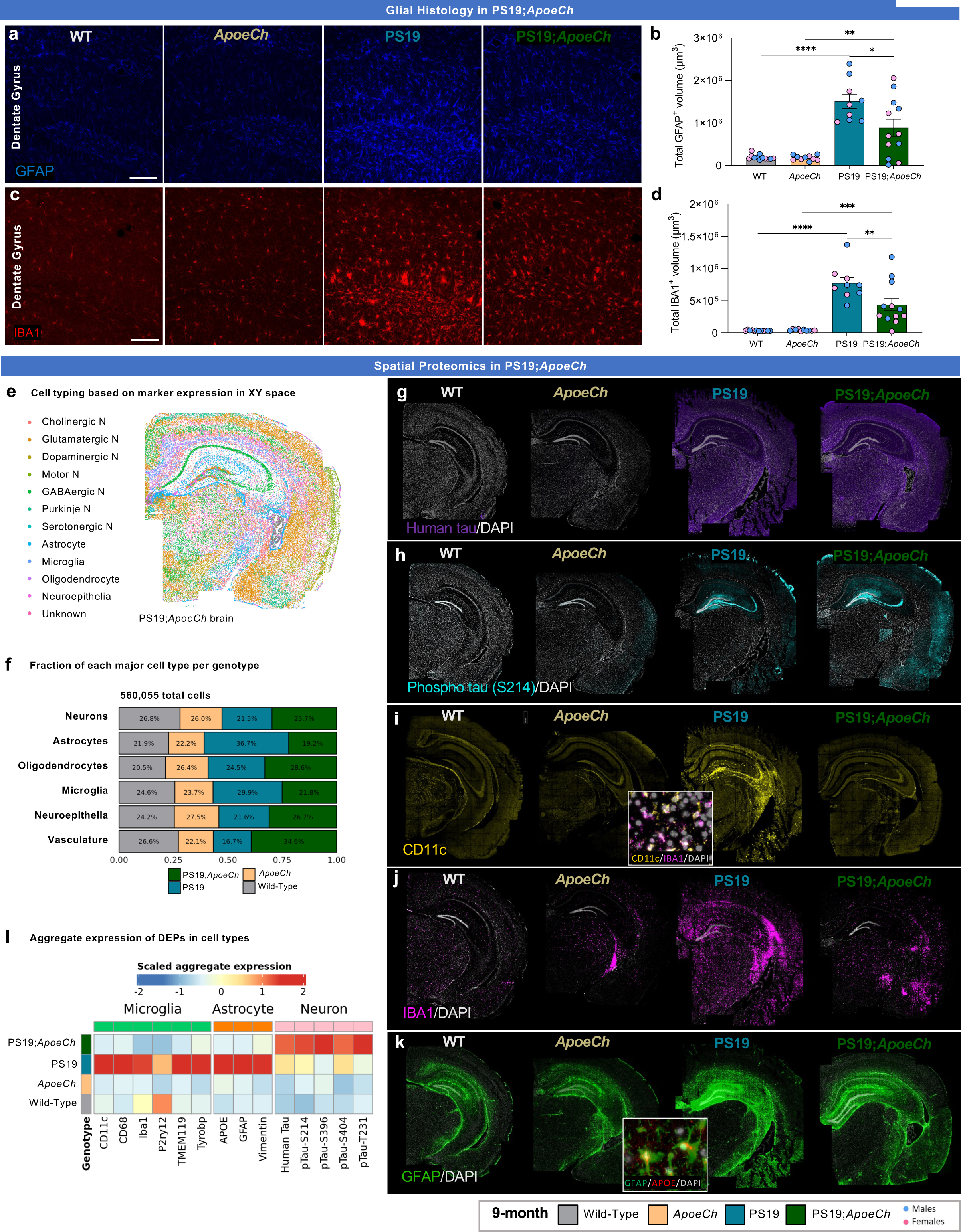
Reduced microglial and astrocytic response to tau induced by *ApoeCh*. **a, c** Representative confocal images of immunostained dentate gyrus for GFAP (blue, a) and IBA1 (red, c) of 9-mo-old WT, *ApoeCh*, PS19, and PS19;*ApoeCh* mice. Scale bar=100µm. **b,d** Quantification of total volume of GFAP^+^ cells (**b**) and IBA1^+^ cells (**d**). n=4-6 mice/sex/genotype. Data are represented as mean ± SEM. Two-way ANOVA followed by Tukey’s post hoc tests to examine biologically relevant interactions. Statistical significance is denoted by *p<0.05, **p<0.01, ***p<0.001, ****p<0.0001. **e** Cell types plotted in XY space. **f** Proportion of the number of cells in each major cell type, grouped by genotype. **g-k** Immunofluorescence images for Human Tau, Phospho-Tau (S214), CD11c, IBA1, and GFAP in representative brains of WT, *ApoeCh*, PS19, and PS19;*ApoeCh* mice. **l** Aggregate expression of differentially expressed proteins (DEPs) in microglia, astrocytes, and neurons across the four genotypes. Expression is normalized for the number of brains in each genotype.

Given the paradoxical observations that *ApoeCh* promotes the microglial response to plaques, but appears to suppress the microglial response to tauopathy, we used spatial proteomics to further characterize the cellular response. As with the 5xFAD study, we acquired data from 10 µm coronal hemibrain slices (9-mo-old mouse samples, n=3 for WT, PS19 and PS19;*ApoeCh* brains, and n=2 for *ApoeCh*) yielding a dataset of 560,055 cells. Cell segmentation was automated using histone, GFAP, and DAPI markers (**Supplemental Fig. 13a**), then automatically assigned to cell types using marker proteins. Plotting these cell types in XY space confirmed accurate cell identification (**Fig. 6e**; all imaged brains shown in **Supplemental Fig. 13b**). Here, the “microglia” category includes both homeostatic and disease-associated microglia. Quantification of cell proportions between genotypes revealed that microglia and astrocyte populations were expanded in PS19 brains, but unchanged in WT, *ApoeCh*, and PS19;*ApoeCh* brains. No change was observed in the proportions of neurons overall across the four genotypes suggesting no neuronal loss is observed in PS19 mice at this age (**Fig. 6f**).

Next, we plotted the aggregate expression of select proteins across microglia, astrocyte, and neuronal populations (**Fig. 6l**). Sample images from the proteomic analyses are shown for neuronal (human Tau and p-Tau S214), microglial (CD11c and IBA1), and astrocyte (GFAP) markers (**Fig. 6g-k**). In microglia, we observed a significant reduction in DAM-related proteins (i.e., CD11c, CD68, TYROBP) in PS19;*ApoeCh* compared to PS19, indicating that microglia have a reduced response to tauopathy in the presence of *ApoeCh*. Similarly, we observe that astrocyte-related proteins GFAP, APOE, and VIM are reduced in PS19;*ApoeCh* compared to PS19, supporting results of histology (**Fig. 6a, b**). Given the biochemical findings that levels of phosphorylated tau (pTau) were altered in hippocampal and cortical tissue lysates, we next focused on pTau expression in neurons. Aggregated protein expression data in neurons showed increased pTau-T231 in PS19;*ApoeCh* brains compared to PS19, consistent with our biochemical data (**Fig. 6l**). Additionally, we identified that aggregate expression of other forms of pTau (i.e., pTau-S396, pTau-S214, and pTau-S404) were also increased in PS19;*ApoeCh* brains compared to PS19 (**Fig. 6l**). These findings show that despite elevated phosphorylated tau accumulation in neurons, there is a marked suppression of both microglial and astrocytic responses in the presence of *ApoeCh*.

### Spatial transcriptomics reveals PS19 transgene-induced changes in gene expression in glial populations are prevented by ApoeCh

To further interrogate the role of *ApoeCh* in the tauopathy murine brain, we performed single-cell spatial transcriptomics analysis, imaging coronal hemibrain slices adjacent to the sections used for spatial proteomics, with data acquired from the cortex and hippocampus. 354,499 cells were processed, clustered, and manually annotated based on gene expression and spatial location to yield a total of 40 clusters, corresponding to 13 clusters of excitatory neurons, 11 inhibitory neurons, 4 astrocytes, 6 oligodendrocytes and oligodendrocyte precursors, 2 microglia, and 4 endothelial and vascular-related clusters (**Fig. 7a**). Sample cell segmentation is shown in **Supplemental Fig. 14a.** Plotting clusters in space again confirmed accurate cell typing (**Fig. 7b**, and all imaged brains are shown in **Supplemental Fig. 14b**). Top marker genes per major cell type were calculated to validate cell annotation (**Supplemental Fig. 15b**), and cell numbers per cluster per genotype are plotted in **Supplemental Fig. 15c**. Calculating the proportions of cells within each major cell type normalized by genotype revealed a marked decrease in the populations of DAAs and DAMs in PS19;*ApoeCh* mice compared to PS19, supporting the reduced glial response seen in PS19;*ApoeCh* from IHC and spatial proteomics data (**Fig. 7c**). Viewing the UMAP split by genotype similarly shows a prominent DAM cluster in PS19 mice that is absent in the other three genotypes (**Supplemental Fig. 15a**).

**Figure 7:**
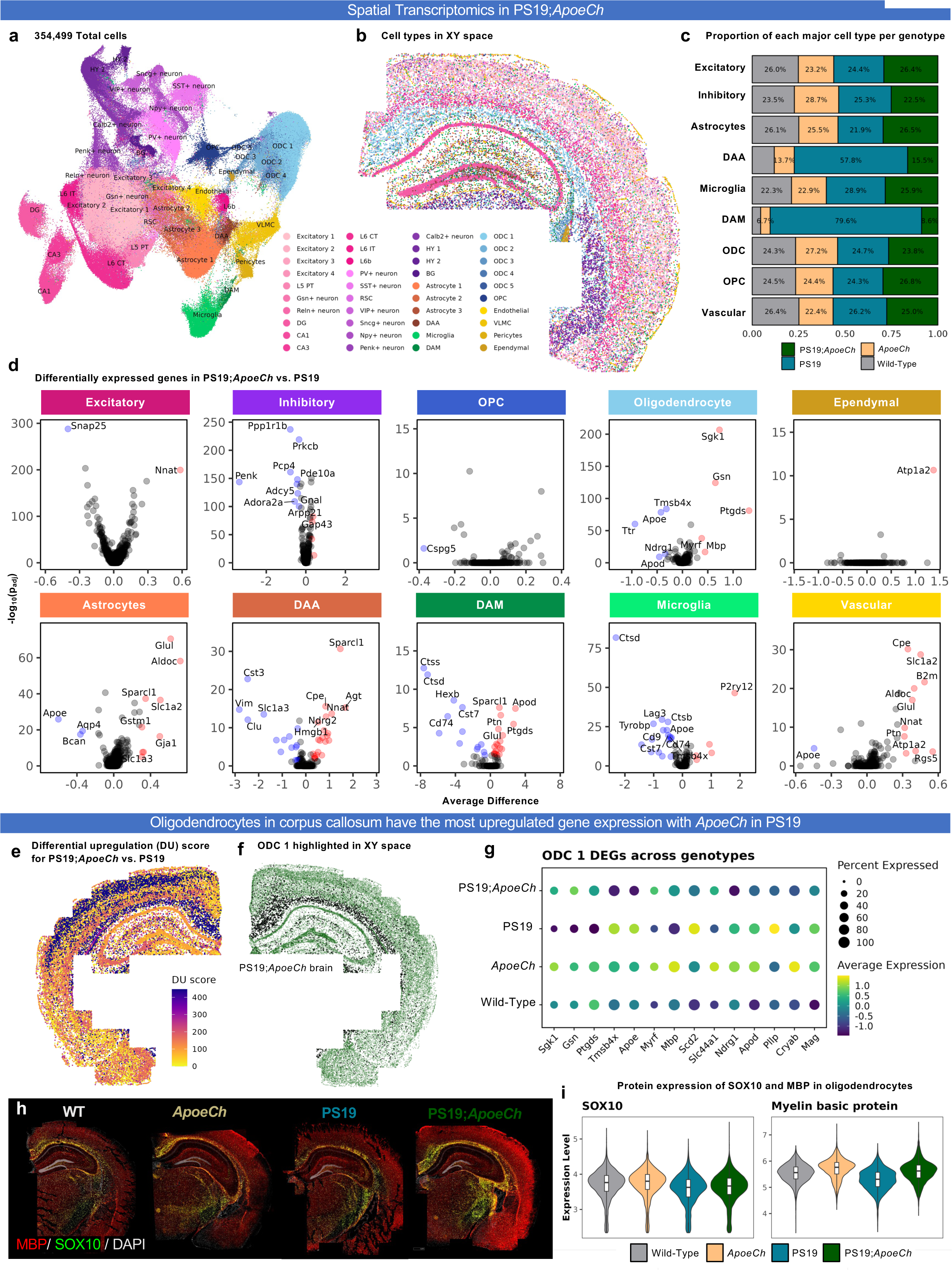
*ApoeCh* prevents tau-induced myelin loss via changes in oligodendrocyte transcriptomics. **a** UMAP of 354,499 cells across 11 hemibrains (n=3/genotype, except n=2 for *ApoeCh*). Clustering at 1.0 resolution gave rise to 40 clusters, which were then manually annotated based on gene expression and anatomical location in space. **b** 40 clusters plotted in XY space. Common legend between **a**, **b**. **c** Proportion of the number of cells in each major cell type, grouped by genotype. Percentages were normalized for the total number of cells in each genotype (i.e., to account for differences in the number of samples per genotype). **d** Volcano plots of DEGs between PS19;*ApoeCh* and PS19 for each major cell type. **e** DU score for PS19;*ApoeCh* vs. PS19 in a PS19;*ApoeCh* brain. **f** ODC 1 cluster highlighted (black) in XY space in a PS19;*ApoeCh* brain. **g** Pseudo-bulk expression of the top DEGs between PS19;*ApoeCh* and PS19 in the ODC 1 cluster across all four genotypes. **h** Immunofluorescence images of MBP (red) and SOX10 (green) in representative brains from each genotype from spatial proteomics dataset. DAPI nuclear stain shown in light grey. **i** Violin plots of normalized and scaled protein expression of SOX10 and MBP in oligodendrocytes across the four genotypes. Overlaid white box-and-whisker plot shows the median and the first and third quartiles, with whiskers representing up to 1.5 times the inter-quartile range (i.e., the distance between the first and third quartiles).

To understand how the presence of tau pathology affects the 40 cell clusters identified in the brain, we performed DGE analysis between PS19 and WT mice. Volcano plots for each of the cell types are shown in **Supplemental Fig. 16a**. Because DAAs and DAMs were not present in WT mice, these cells were grouped into “All astrocytes” and “All microglia” clusters for the PS19 vs. WT comparison. To further simplify interpretation of results, cells from individual clusters were grouped into major cell types and volcano plots generated for each major cell type (**Supplemental Fig. 16c**). Overall, we observed large numbers of significantly upregulated genes in inhibitory neurons (5 upregulated DEGs, defined as p_adj_ < 0.05 and average difference > 0.3), astrocytes (14 upregulated), and microglia (27 upregulated), while oligodendrocytes exhibited both up- (8 upregulated) and down-regulated (4 downregulated) genes. Of all clusters, we detected the most differences in microglia (32 total DEGs) between PS19 and WT brains, with PS19 showing an upregulation in inflammatory-related genes such as *Cd74*, *C1qa*, *C1qb*, *C1qc*, *Ctss*, *Ctsd*, *Cst7*, as well as *Apoe*. Astrocytes in PS19 mice also showed upregulation in reactive astrocytic markers *Vim*, *Clu*, *Aqp4*, and *Ctsb* compared to WT mice (**Supplemental Fig. 16a**). When grouped together, excitatory neurons showed few significant changes, and inhibitory neurons showed upregulation of *Gsk3b*, a kinase implicated in tau phosphorylation, along with *Penk*, *Pcp4*, *Ppp1r1b*, and *Adora2a* (**Supplemental Fig. 16c**). However, when analyzed separately, DGE analysis revealed significant changes in many neuronal populations, notably *Lamp5*+, *Calb2*+, L6b, and hypothalamic (HY) neurons, again reinforcing the utility of analyzing discrete neuronal clusters rather than aggregated data analysis.

Next, we performed DGE analysis between PS19;*ApoeCh* and PS19 mice to investigate the brains’ response to tau laden neurons in the presence of *ApoeCh*. Volcano plots for all 40 cell clusters are shown in **Supplemental Fig. 16b**, and broad collapsed cell clusters shown in **Fig. 7d**. PS19;*ApoeCh* mice exhibit downregulated genes in inhibitory neurons (9 downregulated), DAM (13 downregulated), and microglia (17 downregulated), while genes in astrocytes (9 upregulated), and vascular cells (10 upregulated) were mostly upregulated compared to PS19 mice. In PS19;*ApoeCh* mice, oligodendrocytes (5 upregulated and 5 downregulated) and DAA (22 upregulated and 15 downregulated) cells exhibit gene expression changes in the form of both up- and down-regulated genes. DGE analysis of DAMs revealed significant downregulation of genes related to lysosomal function, including *Ctss*, *Ctsd*, *Cst3*, and *Hexb* (**Fig. 6d**). Notably, many of the genes that were upregulated in microglia for the PS19 vs. WT comparison (i,e., *Ctss*,

*Ctsd*, *C1qa*, *C1qb*, *C1qc*, *Cst3*, *Apoe*, *Tyrobp*) were instead downregulated in the PS19;*ApoeCh* vs. PS19 comparison. A similar reciprocal pattern was observed in astrocytes, neurons, and oligodendrocytes. For example, genes that were upregulated in astrocytes in the PS19 vs. WT comparison, including reactive astrocytic markers *Vim* and *Clu*, were instead downregulated in the PS19;*ApoeCh* vs. PS19 comparison. These data provide evidence that some of the genetic changes induced in different cell populations by the PS19 transgene, such as those associated with glial activation, are reversed by the presence of the *ApoeCh* variant.

#### PS19-induced changes in oligodendrocyte gene expression in the corpus callosum and level of MBP are prevented by ApoeCh

To visualize patterns of differential gene expression in space, we plotted differential upregulation scores between PS19;*ApoeCh* and PS19. Most upregulation in genes in PS19;*ApoeCh* vs. PS19 mice occurs in white matter tracts (ODC 1) and L2/L3 neurons in cortex (Excitatory 1; **Fig. 7e**). Cluster ODC 1 is highlighted in a PS19;*ApoeCh* brain in **Fig. 7f** and localizes to the corpus callosum. We performed pseudo-bulk analysis of the top DEGs between PS19;*ApoeCh* and PS19 mice in the ODC 1 cluster, which are displayed as a dot plot across the four genotypes. These data reveal that several ODC 1 cluster genes altered by PS19 (compared to WT) show a reverse in directionality of expression upon introduction of the *ApoeCh* variant. These genes include *Sgk1*, *Gsn*, and *Ptgds* (downregulated in PS19 compared to WT and upregulated in PS19;*ApoeCh* compared to PS19)*, Tmsb4x*, *Apoe*, *Scd2*, and *Pllp* (upregulated in PS19 compared to WT and reduced in PS19;*ApoeCh*). We also found several ODC 1 genes were upregulated by *ApoeCh,* independent of PS19 background, including *Mbp*, *Myrf* and *Slc44a1*, of which *Mbp* was restored to WT levels in PS19;*ApoeCh* mice. Given the restoration of ODC 1 *Mbp,* and other oligodendrocyte-associated genes, we used the spatial proteomics dataset to analyze the level of myelin basic protein (MBP) and SOX10 in oligodendrocytes. Consistent with gene expression data, MBP is higher in *ApoeCh* and *PS19;ApoeCh* compared to WT and PS19 mice (**Fig. 7h, i**).

SOX10, an oligodendrocyte marker, is downregulated in PS19 mice compared to WT, and less downregulated in the presence of *ApoeCh*. Together, these data indicate that PS19 results in alterations to oligodendrocytes, including MBP expression, which are largely prevented by *ApoeCh*.

### PS19-induced changes in CA1 neuron gene expression and CA1 synaptic densities are prevented with ApoeCh

Having shown and explored the changes associated with differential gene upregulation, we next plotted differential downregulation scores between PS19;*ApoeCh* and PS19 (**Fig. 8a**). These data highlight DAM, microglia, DAA, and CA1 as the clusters with the most downregulated genes in PS19;*ApoeCh* compared to PS19 mice (DD scores were plotted in a PS19;*ApoeCh* brain, which has limited DAM). We focused initially on the changes in the CA1 cluster, which localized spatially exclusively in the CA1 (**Fig. 8b**). Pseudo-bulk analysis was performed for the top DEGs in the CA1 cluster across the four genotypes (**Fig. 8c**). These data show that many genes that were upregulated in PS19 relative to WT mice (i.e., *Tmsb4x*, *Camk2b*, *Hsp90ab1*, *Zbtb20*, *Atp2b1*, *Sstr4*) were subsequently downregulated in PS19;*ApoeCh* compared to PS19 mice. Conversely, genes that were downregulated in PS19 mice relative to WT (i.e., *Atp2a2*, *Slc8a1)*, genes involved in calcium homeostasis and neuronal signaling, were upregulated in PS19;*ApoeCh* compared to PS19 mice. There were also several genes (i.e., *Gria1*, *Grin2b*, *Wasf1*), involved in synaptic plasticity, synaptic transmission, and long-term potentiation that showed low expression in WT, *ApoeCh*, and PS19, but which were upregulated in PS19;*ApoeCh* (**Fig. 8c**) [59–62].

**Figure 8:**
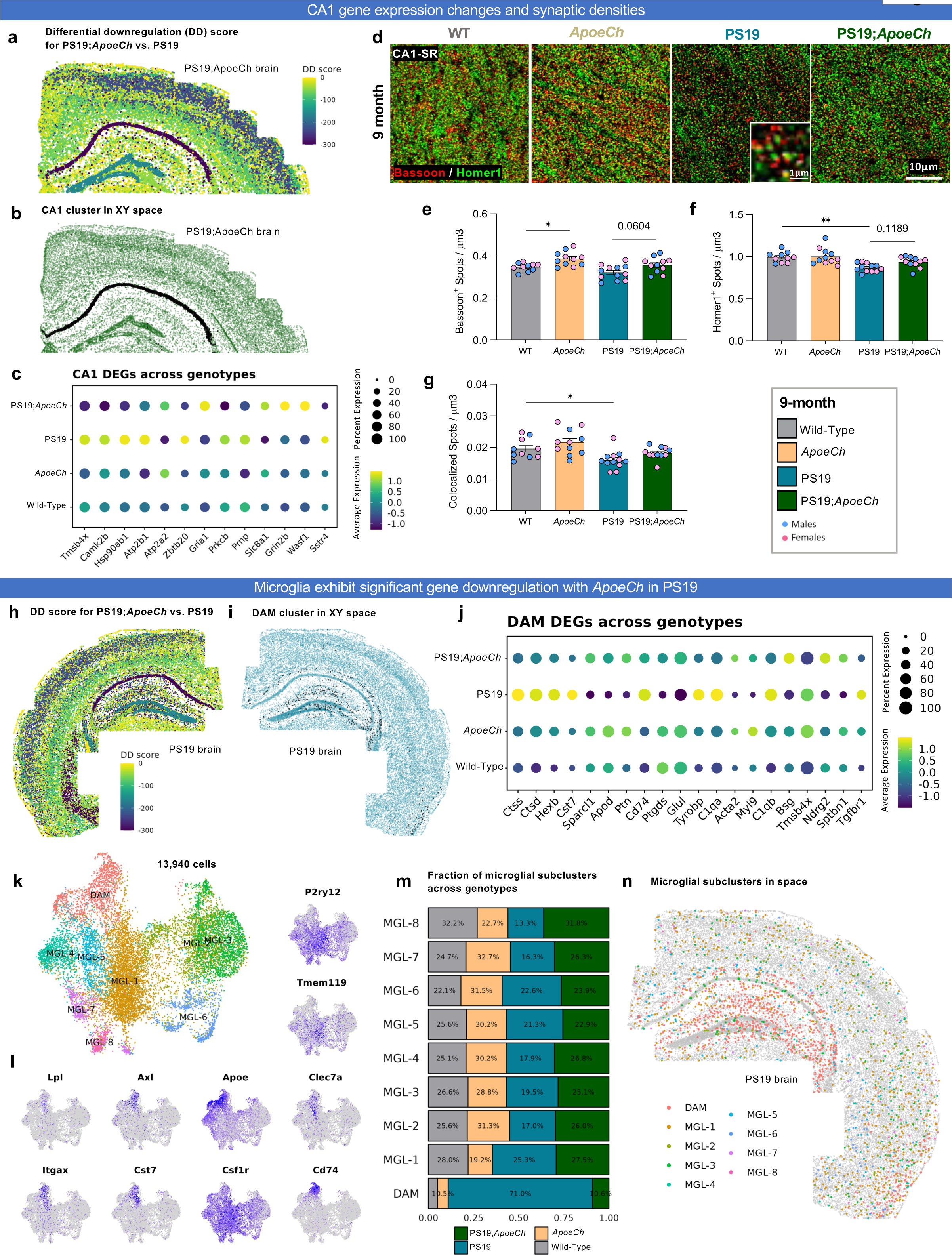
*ApoeCh* suppresses microglial response to tau and partially suppresses tau-induced synaptic loss in the PS19 mouse model. **a** DD score for PS19;ApoeCh vs. PS19 in a PS19;*ApoeCh* brain**. b** CA1 cluster highlighted (black) in XY space in a PS19;*ApoeCh* brain. **c** Pseudo-bulk expression of the top DEGs between PS19;*ApoeCh* and PS19 in the CA1 cluster across the four genotypes. **d** Representative super-resolution images of Bassoon and Homer1 synaptic markers for WT, *ApoeCh*, PS19, and PS19;*ApoeCh* mice at 9 mo of age. Scale bar = 10mm. Insert scale bar = 1mm **e-g** Quantification of Bassoon+ spots per mm^3^, Homer1+ spots per mm^3^ and colocalized Bassoon+/Homer1+ synaptic spots per mm^3^ showing decreased synaptic puncta in PS19 mice compared to WT mice. Three images per mouse and n=5-6 mice/sex/genotype. Data are represented as mean ± SEM. Two-Way ANOVA followed by Tukey’s post hoc test to examine biologically relevant interactions. *p<0.05, **p<0.001. **h** DD score for PS19;*ApoeCh* vs. PS19 in a PS19 brain. **i** DAM cluster highlighted (black) in XY space in a PS19 brain. **j** Pseudo-bulk expression of the top 20 DEGs between PS19;*ApoeCh* and PS19 in the DAM cluster across the four genotypes. **k-n** Microglial subclustering analysis. **k** UMAP of 13,940 microglial cells in space. Clustering at 0.4 resolution yielded 9 subclusters, excluding the smallest subcluster that had fewer than 100 cells. **l** Feature plots of common microglial and DAM marker genes in UMAP space. **m** Proportion of the number of cells in each microglial subcluster, grouped by genotype. Proportions are normalized for the total number of microglia in each genotype. **N** Microglial subclusters plotted in XY space in a PS19 brain.

To examine the functional changes occurring in CA1, we immunostained for presynaptic marker Bassoon and postsynaptic marker Homer1 in 9-mo-old mouse brains and imaged synaptic puncta via super resolution microscopy (**Fig. 8d**). Interestingly, the *ApoeCh* variant increased Bassoon presynaptic puncta independently of PS19 transgene (**Fig. 8e**). Significant reductions in both Homer1 postsynaptic puncta and colocalized Homer-Bassoon pre- and post-synaptic puncta were observed in PS19 mice compared to WT, but no reduction was observed between *ApoeCh* and PS19;*ApoeCh* brains (**Fig. 8f, g**), suggesting that the *ApoeCh* variant can partially suppress the synaptic loss seen in the PS19 mouse model.

### Spatial transcriptomics reveal a suppressed transition to disease-associated microglia in response to tauopathy with ApoeCh

DAMs and microglia had the greatest differential downregulation score in the PS19;*ApoeCh* vs. PS19 comparison (**Fig. 8h**; −336.2 for DAM, −378.5 for microglia, −279.8 for CA1). We plotted DAMs in a PS19 brain (**Fig. 8i**), showing that these cells localize to areas of the hippocampus and piriform cortex where tau pathology is seen. We performed pseudo-bulk analysis of the top 20 DEGs between PS19;*ApoeCh* and PS19 DAMs and plotted their expression levels across the four genotypes in DAMs (**Fig. 8j**). As expected, many genes upregulated in PS19 mice relative to WT (i.e., *Ctss, Ctsd, Hexb, Cst7, Cd74, Tyrobp, C1qa, C1qb,* and *Tgfbr1*) are not upregulated in PS19;*ApoeCh* mice. Similarly, several genes downregulated in PS19 compared to WT (i.e., *Sparcl1*, *Apod*, *Ptn*, *Ptgds*, *Glul*, *Acta2*, *Myl9*, *Bsg*, *Ndrg2*, *Sptbn1*) are expressed at WT level in PS19;*ApoeCh*.

To further investigate how microglial cells are impacted by the introduction of *ApoeCh* on the PS19 background, we clustered all the microglia and DAMs from the four genotypes (13,940 total cells) into 9 new subclusters (**Fig. 8k**). Feature plots of microglial marker genes are shown in **Fig. 8l**, separating DAMs from homeostatic microglia. We identified 8 subclusters of homeostatic microglia (MGL1-8) and one subcluster of disease-associated microglia (DAM). Quantification of the proportion of cells in each of these subclusters reveals the distinct changes in each of these subclusters due to PS19 and the introduction of *ApoeCh*. PS19 mice show a reduction in the proportion of several homeostatic microglial clusters compared to WT (MGL-2,3,4,7,8) and a marked increase in DAMs. All microglial cluster proportions appear to be at WT levels in PS19;*ApoeCh* mice (**Fig. 8m**). Plotting these microglial subclusters in space identifies that while homeostatic microglia are expressed throughout the brain, cells in the DAM cluster are localized to the hippocampus in PS19 mice (**Fig. 8n**).

### Comparison of gene expression effects of ApoeCh in 5xFAD and PS19 mice

We have shown that the *ApoeCh* variant has a differential impact on the microglial response dependent on disease pathology, enhancing the glial response to amyloid plaques in 5xFAD mice but tempering the response to tau pathology in PS19 mice. To gain insight into the molecules and underlying mechanisms of this differential impact of the *ApoeCh* variant on microglia, we first merged our spatial transcriptomic datasets from both the 5xFAD;*ApoeCh* and PS19;*ApoeCh* mice, yielding 41,790 Microglia/DAM cells (**Fig. 9a**). Next, we visualized microglial cell counts and proportions in each of the distinct animal lines (**Fig. 9b-c**). These data show the pronounced number and proportion of DAMs that make up the microglial population in 5xFAD (∼75%), which appears expanded in 5xFAD;*ApoeCh* (∼82%) mice, indicating that the *ApoeCh* variant encourages a DAM-like response or promotes the transition of more homeostatic microglia cells into DAMs in the presence of plaques, as we have described. In PS19 mice, we observe a much smaller number and proportion of DAMs (∼20%), which are nearly absent in PS19*;ApoeCh* mice (**Fig. 9b-c**). These data confirm the opposing effects on numbers of DAMs in response to either plaques or tau pathology in the presence of *ApoeCh*.

**Figure 9:**
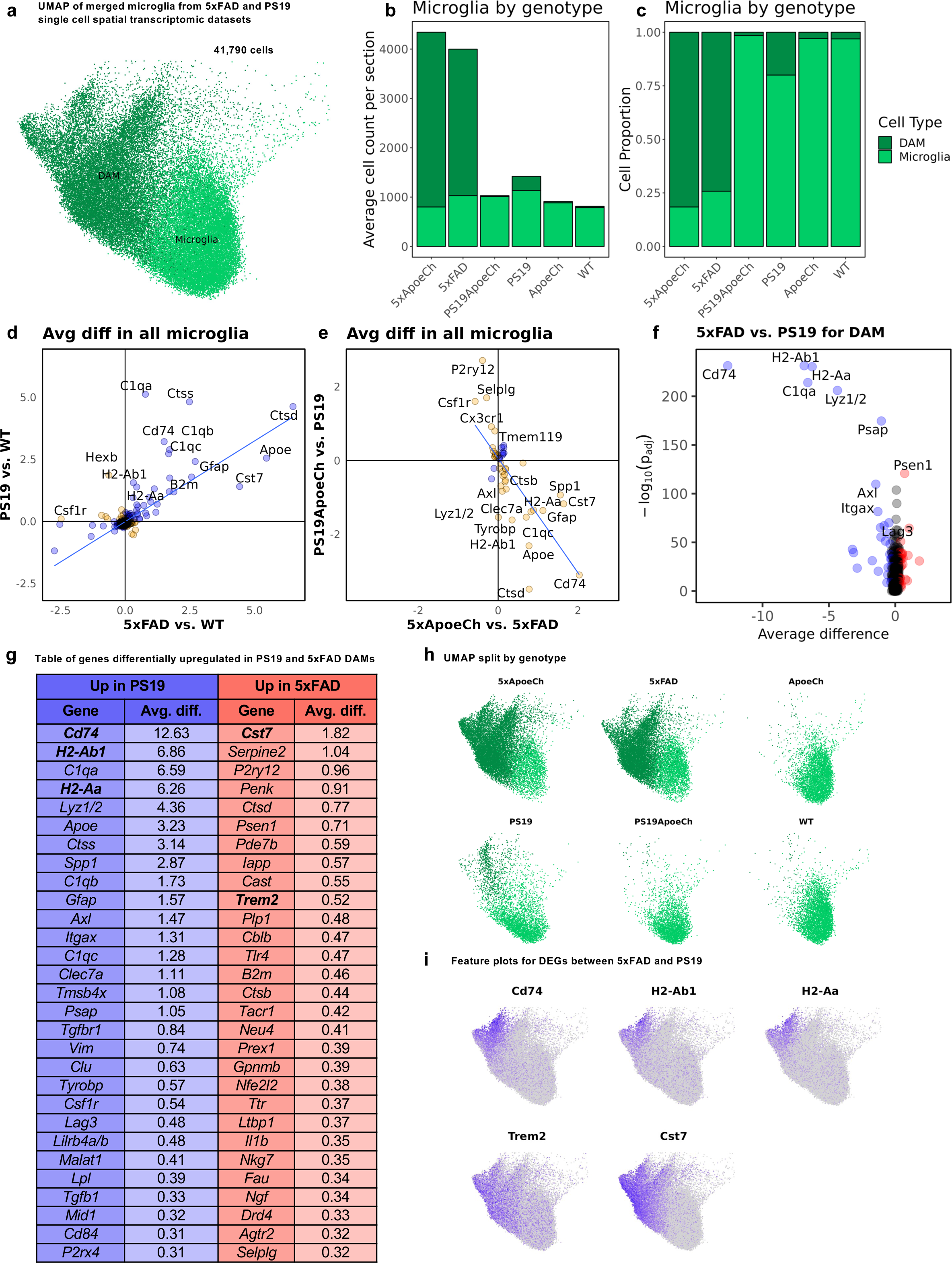
Microglia has differential responses to tau and plaque. **a** UMAP of 41,790 microglia merged and integrated across both 5xFAD;*ApoeCh* and PS19;*ApoeCh* datasets (n=3 for 5xFAD;*ApoeCh*, 5xFAD, PS19;*ApoeCh*, and PS19; n=5 for *ApoeCh*; n=6 for WT). Cells are labelled by their previous annotations from their original datasets prior to merging. **b** Average microglial cell counts per section across each of the six genotypes. Cell counts for each genotype were divided by the total number of samples in that genotype. **c** Proportion of the number of cells in each genotype, grouped by DAM versus homeostatic microglia. **d** Scatterplot of the average difference in all microglia for all significant genes (i.e., p_adj_ < 0.05) between 5xFAD vs. WT and PS19 vs. WT comparisons. Directly correlated genes (blue) occur in the same direction for both comparisons, while inversely correlated genes (orange) occur in opposite directions for each comparison. Linear regression line showing the relationship between the two comparisons plotted in blue. **e** Scatterplot of the average difference in all microglia for all significant genes between 5xFAD;*ApoeCh* vs. 5xFAD and PS19;*ApoeCh* vs. PS19. **f** Volcano plot of DEGs between 5xFAD and PS19 DAMs. **g** Full list of genes from **f**, in order of magnitude of average difference. Genes that are blue are upregulated in PS19 DAMs (i.e., downregulated in 5xFAD DAMs), while genes in red are upregulated in 5xFAD DAMs (i.e., downregulated in PS19 DAMs). **h** UMAP of merged microglia from 5xFAD;*ApoeCh* and PS19;*ApoeCh* datasets split by genotype. DAMs in PS19 genotype are localized to a. **i** Feature plots of DAM markers *Cd74*, *H2-Ab1*, *H2-Aa*, *Trem2*, and *Cst7* in all microglia.

To gain insight into the different genes and pathways involved in the differential *ApoeCh*-mediated microglial response to plaques and tauopathy, we compared how microglia respond to each of these pathologies. To capture this, all DEGs between 5xFAD vs. WT and PS19 vs. WT in all microglia and DAMs were plotted using the average difference of each gene among the two comparisons (**Fig. 9d**). Notably, many gene expression changes were shared and correlated between the two responses. In response to amyloid and tau, microglia exhibit down regulation in similar homeostatic genes, and an upregulation in many DAM genes (i.e., *Ctsd*, *Apoe*, *Cst7*, etc.). However, several genes showed differential regulation between the responses. For example, *C1qa-c, Ctss, Cd74, H2-Ab1, and H2-Aa* were more upregulated in the PS19 vs. WT response compared to the 5xFAD vs. WT response. Conversely, *Apoe* and *Cst7* were more upregulated in the 5xFAD vs. WT response. Homeostatic genes *Hexb* and *Csf1r* were downregulated in 5xFAD mice compared to WT, but unchanged or upregulated in PS19 vs. WT mice. Thus, while the overall response is similar (i.e., 5xFAD and PS19 microglia upregulate DAM genes in response to pathology), there are differences in related groups of genes in the magnitude of their response. For example, DAMs make up 74.2% of all microglia in 5xFAD brains compared to 20.0% in PS19 brains. Despite this difference, the overall response is remarkably similar in magnitude (i.e., average difference) in 5xFAD vs. WT mice compared to PS19 vs. WT mice, suggesting that microglia in PS19 have highly elevated DAM gene expression on a per-microglia basis compared to those in the 5xFAD brain. We plotted the average difference for all the DEGs from 5xFAD;*ApoeCh* vs. 5xFAD and PS19;*ApoeCh* vs. PS19 comparisons (**Fig. 9e**). As expected, gene expression changes were inversely correlated to each other, with most genes being upregulated in 5xFAD;*ApoeCh* vs. 5xFAD microglia/DAM, but downregulated in PS19;*ApoeCh* vs. PS19 microglia/DAM, reinforcing the opposing effects of *ApoeCh* on the microglial response to the two pathologies.

To directly compare the DAM response to either plaques or tauopathy, we plotted DEGs between DAMs in 5xFAD vs. PS19 mice (**Fig. 9f**). Consistent with fewer PS19 DAMs exhibiting the same magnitude in overall gene expression changes as the far greater number of 5xFAD DAMs, we observed that many genes were upregulated in PS19 DAMs compared to 5xFAD. Antigen presentation-associated genes were highly upregulated in PS19 compared to 5xFAD DAMs, including *Cd74, H2Ab1,* and *H2-Aa* (**Fig. 9g**). On the other hand, several genes were more upregulated in DAMs from 5xFAD brains compared to DAMs from PS19 brains, including *Cst7* and *Trem2* (**Fig. 9g**). Splitting the UMAP by genotype (**Fig. 9h**) reveals that PS19 mice only express a subset of DAMs that are found in 5xFAD mice. Visualization of *Cd74, H2-Ab1*, and *H2-Aa* on the UMAP show that DAMs in PS19 are specifically enriched in these genes, while *Trem2* and *Cst7* are enriched in DAMs found in 5xFAD mice (**Fig. 9i**). These results indicate that the responses of microglia to either plaques or tauopathy are fundamentally different from one another, providing a potential explanation as to how the *ApoeCh* variant can produce such contrasting effects.

## DISCUSSION

The recent identification of an ADAD PSEN1-E280A carrier who was homozygous for the rare APOE Christchurch variant (R136S, APOECh) and who was resistant to the onset of ADAD dementia until her 70’s was a remarkable discovery [12]. Due to the inherent nature of a single human case report, it was formally unknown whether the APOECh variant was responsible for the resilience to AD observed in this person. For this reason, we used genetically modified mouse models to investigate whether introduction of the Christchurch variant into mouse APOE could modify development of amyloid and tau-related pathologies. Using spatial transcriptomics and proteomics, we show that the mouse *ApoeCh* variant enhances plaque-associated microgliosis in 5xFAD mice but suppresses the glial response to tauopathy in PS19 mice. These results support that the APOECh variant could contribute to the resilience to AD observed in the PSEN1-E280A carrier in the case report and offer insight into the potential mechanisms for this effect. While this report was being prepared, independent studies reporting similar findings have very recently been published [32, 63, 64]. We discuss our results and conclusions in the context of those studies.

Mouse models are powerful systems to explore mechanisms underlying the development of pathology and the brains response to it. While no single mouse model effectively recapitulates human AD, the field routinely utilizes mice that have been engineered to develop either extensive amyloid pathologies (e.g., the 5xFAD model) or tauopathy (e.g., the PS19 model). Both models rely on transgene overexpression and familial mutations (such as ADAD mutations in 5xFAD mice and frontotemporal dementia mutations in PS19 mice) and are not well suited for studying the causative events that lead to the development of pathology in sporadic AD [34–37]. However, they are useful for examining the brain’s response to Aβ- and tau-associated pathology, in this instance in the context of the APOE Christchurch variant.

We decided to analyze the effect of introducing the Christchurch variant into the mouse APOE protein (mAPOE) rather than the human protein (hAPOE). Both mAPOE and hAPOE4 have arginine at the equivalent positions of 112 and 158. However, mAPOE and hAPOE have incomplete sequence identity between those two residues, including between AA136 to 150 (human numbering) that is the receptor region involved in binding to LDLR and HSPG. Modifying the mouse *Apoe* locus allows us to exclude the potential confound associated with interaction of hAPOE3/4 with murine receptors.

Three other APOE Christchurch mouse models have recently been described, all with Christchurch introduced into human sequences (including both hAPOE3 and hAPOE4) with some contrasting results [32, 63, 64]. Introduction of hAPOE3Ch into APP/PS1 mice (with and without injection of human AD-tau) resulted in decreased plaque load and AD-tau seeding and spreading, alongside increased microglial reactivity around plaques [32]. In PS19 mice, a homozygous R136S mutation rescued APOE4-dependent tau pathology and hippocampal atrophy, with a reduction in microgliosis [63]. In addition, the R136 mutation in PS19 mice (on a hAPOE3 background) resulted in protection against tau pathology and tau-induced synaptic deficits while suppressing interferon microglial activation [64]. Thus, the impact of the Christchurch variant appears to be dependent on the stimulus – i.e., plaques or tau-laden neurons. In mice homozygous for the Christchurch variant, these studies indicate that microglial activation states are suppressed in PS19 mice (tau-associated pathology), promoting a homeostatic or “disease protective” response, whereas they are enhanced in APP/PS1 mice (amyloid-associated pathology), promoting a “disease-associated” state. To dissect the differential microglial responses to pathology, we generated *ApoeCh* homozygous mice independently with both 5xFAD and PS19 lines, which enabled us to directly compare the impact of *ApoeCh* on the response to amyloid and tau pathology individually. The results we obtained complements, validates, and helps reconcile these previous findings.

Our study reveals that introduction of *ApoeCh* into the 5xFAD mouse model reduces plaque load and reactive astrocytic and microglial presence, as well as ameliorates plaque-induced neuritic and axonal damage, consistent with the study involving humanized *APOE3Ch* knock-in mice crossed to APP/PS1 mice [32]. Despite a reduction in glial numbers, spatial proteomics and transcriptomics reveal a significant upregulation of disease-associated microglia and astrocytes in 5xFAD;*ApoeCh* compared to 5xFAD mice. In agreement with Chen et al. [32] who showed increased myeloid cell phagocytosis in humanized APOE3Ch, we show that *ApoeCh* produces upregulation of several genes associated with autophagy/endocytosis and antigen presentation (*Mid1, Cd74, H2-Aa, H2-Ab1, Spp1, Gpnmb*, *Cst7*) in 5xFAD mice. A recent report provides evidence that autophagy plays an important role in regulating microglial activation (i.e., DAM phenotype induction) and that deficiency in autophagy leads to senescence, reduces DAMs, impairs Aβ clustering around plaques, and aggravates AD pathology [65]. Given the reduction in amyloid-associated damage, these data implicate a neuroprotective role of enhanced DAM/microglial reactivity to amyloid pathology with the introduction of *ApoeCh* in 5xFAD mice.

Crossing *ApoeCh* mice to PS19 mice reveals the divergent impact of *ApoeCh* on pathology and microglia in a mouse model of tauopathy. We show that introduction of *ApoeCh* reduces microglia and reactive astrocytes, similar to 5xFAD mice, but is accompanied by a reduction in DAM and DAA as well as associated proteins and genes, thereby promoting a more homeostatic state, and counter to changes in models of amyloidosis. Contrary to studies in humanized mice [63, 64], our study does not show that *ApoeCh* rescues tau pathology or functional deficits. Chen et al. [32] report no difference in tau with introduction of humanized *APOE3Ch* in the absence of amyloidosis. In line with humanized *APOECh* observations, we demonstrate a diminished DAM and astrocyte response to tau [63, 64]. Moreover, we provide evidence that *ApoeCh* results in the rescue of tau-induced synaptic and myelin loss in PS19, associated with the rescue of transcription changes in oligodendrocytes and neurons [63, 64]. Together, our results provide insight into the duality of the Christchurch variant, which enhances DAM reactivity against plaques while promoting a more homeostatic or “disease protective” response to tau.

Given the multifaceted roles of APOE and its receptors in AD pathologies, the protective effects conferred by the Christchurch variant likely involve multiple mechanisms. Previous studies provide evidence that the Christchurch variant reduces AD pathology and associated CNS damage at least in part through alterations in receptor binding to LRP1, LDLR and/or HSPGs [12, 32, 63]. Consistent with data indicating that *ApoeCh* decreases the binding affinity to these molecules [32, 66], our data shows that introduction of *ApoeCh* causes elevated plasma cholesterol. However, our data also indicates that the Christchurch variant may influence Aβ pathology by directly affecting microglial programming. 5xFAD;*ApoeCh* mice show changes in DAM-associated gene expression that are heavily implicated in inflammation, particularly autophagy and phagocytosis. Krasemann et al. showed that the DAM (DAM/MGnD) signature is dependent on the TREM2-APOE pathway [46]. Thus, triggering receptor expressed on myeloid cells 2 (TREM2), a microglia cell surface receptor involved in phagocytosis and regulation of inflammation, could also in part contribute to protective *ApoeCh*-associated amyloid changes. APOE-TREM2 interactions have been found to facilitate phagocytosis of apoptotic neurons and increased uptake of Aβ [67–70].

HSPGs are responsible for neuronal uptake of tau and the R136S mutation causes impaired binding affinity of APOE to HSPGs, leading to reduced tau pathology [12, 32, 63, 71, 72]. We show that introduction of *ApoeCh* in PS19 mice leads to a downregulation in DAM and disease associated astrocytes (DAA), coupled with the upregulation of oligodendrocyte-related genes and rescue of myelin loss. Shi et al. [73] report that overexpression of LDLR in PS19 mice suppresses microglial activation, enlarges pools of oligodendrocyte progenitor cells, and enhances preservation of myelin integrity. Thus, introduction of *ApoeCh* in tauopathy murine models might also promote or alter LDLR signaling. These data also suggest that *ApoeCh* may also modulate the interactions or crosstalk between microglia, astrocytes, and oligodendrocytes. Activated microglia, via C1q release, have been implicated as potential mediators of neurotoxic effects, resulting in reactive astrocyte-mediated neuronal and oligodendrocyte death [74]. Notably, we also observed reduction in *C1q* expression in PS19;*ApoeCh* mice. Our spatial transcriptomics data also reveals that PS19 mice exhibit an upregulation in several MHC II-antigen presentation-related genes, which is dampened by *ApoeCh*. A prior investigation has highlighted the higher inflammatory reactivity (i.e., microglial MHC II+ proteins and T cells) associated with tau-induced neurodegeneration, but not those with amyloid deposition [75]. Moreover, *Apoe* deletion rescued brain atrophy and altered microglia and T cells, tying together the role of APOE and the innate and adaptive immune responses to tau vs. plaques via microglia [75]. In line with this, we report that *ApoeCh* produces an overall decrease in inflammatory response to tau, suggesting the impaired binding of APOE to its receptors plays an important role in neuroinflammation.

The role of microglia in the development and progression of AD is a hotly debated topic [76–79]. Genetic association data implicate microglial expressed genes, and hence microglia, with altered risk for developing the disease, but the underlying mechanisms are not yet fully understood [80–82]. Further, the microglial responses to plaques, tangles, and dying neurons are prominent in the human AD brain as well as in animal models, while aberrant microglial activity is associated with synaptic and neuronal loss [83–85]. The current consensus is that the microglial response to plaques confers protection. Indeed, approaches are underway to further induce this microglial response through TREM2 agonists and other approaches [86, 87]. Conversely, the microglial response to tauopathy appears to drive neurodegeneration. For instance, microglial elimination from PS19 mice or suppression of inflammation prevents tau-induced neurodegeneration [88–90], while APOE2 or *Apoe* KO protects against, and APOE4 exacerbates, tauopathy-induced neurodegeneration [24]. Understanding how changes in microglial biology, triggered by different stimuli or pathological states, can induce either protection or damage should yield insights for the timing of therapeutic interventions. Introduction of the *ApoeCh* variant represents a therapeutic strategy that presents a nuanced approach to modulating distinct microglial responses in the presence of amyloid vs. tau pathology. Understanding the interactions of APOE and how the R136S variant alters those interactions may provide small molecule opportunities for targeted glial therapeutics. Outside of small molecules that target the R136 region of APOE, introduction of APOE3Ch into humans may offer another avenue for lifelong protection, whether through genome editing or other means. In this study, we employed single-cell spatial transcriptomics imaging to examine the transcriptional changes induced by the introduction of the *ApoeCh* variant. This technique offers a powerful advantage by providing both transcriptomic and spatial information, which is lacking in traditional single-cell/single-nucleus approaches. However, it is limited by the pre-selected mouse neuroscience 1000-plex probe set, which constrains the level of exploratory or unbiased investigation compared to the aforementioned techniques. Although the current probe set broadly covers major biological pathways, it restricts in-depth investigations due to the limited selection of RNA targets. Additionally, while crossing the *ApoeCh* mice with amyloidosis and tauopathy models in parallel provides important insights into how the variant independently affects Aβ plaques and tau, it is important to investigate the effect of the *ApoeCh* variant in a mouse model that exhibits both Aβ and tau pathologies, ideally not produced via classical transgenes. This approach should offer a more comprehensive understanding of the role and mechanism of APOECh in Alzheimer’s disease.

## CONCLUSIONS

The R136S mutation occurs within a conserved HSPG/LDLR binding region of APOE, and its introduction in the murine genome/proteome allows us to study its function in its natural interactome and complement findings from humanized APOECh mouse models. By independently crossing with a model of amyloidosis and a model of tauopathy, we show that the *ApoeCh* variant effects the microglial response to both pathologies, but in opposite directions. *ApoeCh* promotes the response of microglia to plaques, but suppresses the response to tauopathy, seemingly providing protection in both situations. Thus, *ApoeCh* appears to provide a nuanced and context-dependent modulation of the microglial response, highlighting the importance of microglial biology and APOE to the development and progression of AD.

## LIST OF ABBREVIATIONS

AD: Alzheimer’s disease
ADAD: Autosomal Dominant Alzheimer’s Disease
APOE: Apolipoprotein E
ApoeCh: Apoe Christchurch
APP: Amyloid precursor protein
Aβ: Amyloid-beta
CNS: Central nervous system
DAA: Disease-associated astrocyte
DAM: Disease-associated microglia
DEGs: Differentially expressed genes
DGE: Differential gene expression
EOAD: Early-Onset Alzheimer’s Disease
hAPOE: human APOE protein
HSPGs: Heparan sulfate proteoglycans
LDLR: Low-density lipid receptor
LOAD: Late-onset Alzheimer’s Disease
LRP1: Lipoprotein-related receptor 1
mAPOE: Mouse APOE protein
MERFISH: multiplexed error-robust fluorescence in situ hybridization
NfL: Neurofilament light chain
PBS: Phosphate buffered saline
PFA: Paraformaldehyde
PSEN: Presenilin
UMAP: Uniform Manifold Approximation and Projection
WT: Wild-type

## ADDITIONAL FILES

**Supplemental Table 1 - Location of potential off-target sites for crRNA TMF1648 on mouse chromosome 7**

The desired target site within *Apoe* locus is listed, plus the sequence of each of the 12 potential off-target sites on mouse chromosome 7 (GRCm38/mm10 nucleotide numbering). Green text denotes the 11-nucleotide seed region proximal to the PAM site. Mismatches to the guide are indicated by lowercase bold letters and the number of mismatches (including the number within the seed region) is shown. No difference was found in sequence between the C57BL/6J WT and *Apoe^em1Aduci^* alleles at the six potential off-target sites analyzed (Supplementary Fig. 1b).

**Supplementary Table 2: Primers for PCR amplification and sequencing for off-target analysis.** Forward (For) and reverse (Rev) primer sequences are listed for each potential off-target site. The off-target code corresponds to the panels in Supplementary Fig. 1.

**Supplemental Figure 1: Sequence of the *Apoe^em1Aduci^* (*ApoeCh*) allele and off-target site analysis for crRNA TMF1648 on mouse chromosome 7.** (**a**) Amino acid (aa) sequence alignment between human and mouse APOE and DNA sequence of the wildtype and *ApoeCh* alleles in the region of the APOE R136S (R128S in mouse mature APOE). Red-colored nucleotides denote the missense codon while the green nucleotides denote synonymous base changes introduced to prevent recutting of the targeted site. (**b**) Chromatograms of DNA sequence in wildtype and *ApoeCh* heterozygous mice, with colored asterisks corresponding to the colored nucleotides in the above sequence. (**c-i**) Chromatograms of N3F2 wildtype and *ApoeCh* homozygous offspring at six potential off-target sites on mouse chromosome 7. No difference was found in sequence between the B6J WT and *ApoeCh* homozygotes at each of the six potential off-target sites analyzed. The black underline denotes the crRNA target sequence while the blue underline denotes the NGG PAM site.

**Supplemental Figure 2: Behavioral analysis of 4- and 12-mo-old WT, ApoeCh, 5xFAD, 5xFAD;ApoeCh mice. a,b** Weight of 4-mo-old (**a**) and 12-mo-old (**b**) mice taken at euthanizing day. **c,e** Total time mice spent in the center of open field behavioral assay of 4-mo-old (**c**) and 12-mo-old (**e**) in the 5 min recording time. **d,f** Mean velocity mice traveled in the center of the open field in 5 min of 4-mo (**d**) and 12-mo-old mice (**f**). **g,h** Total time 4-mo (**g**) and 12-mo-old (**h**) mice spent in open arms of the elevated plus maze behavior test. n=4-6 mice/sex/genotype. Data are represented as mean ± SEM. Two-way ANOVA followed by Tukey’s post hoc tests to examine biologically relevant interactions. Statistical significance is denoted by *p<0.05, **p<0.01, ***p<0.001, ****p<0.0001. # denotes trending significance.

**Supplemental Figure 3: Quantification of insoluble and soluble Aβ in micro-dissected hippocampi and cortices of 4-mo-old 5xFAD, 5xFAD;*ApoeCh* mice. a, d** Cortical and hippocampal soluble Ab40 (**a,b**) and Ab42 (**c,d**) level measured via MSD of 4-mo-old mice. **e-h** Cortical and hippocampal soluble Ab42 (**e,f**) and Ab42 (**g,h**) level measured via MSD of 4-mo-old mice. n=4-6 mice/sex/genotype. Data are represented as mean ± SEM. Two-way ANOVA followed by Tukey’s post hoc tests to examine biologically relevant interactions. Statistical significance is denoted by *p<0.05, **p<0.01, ***p<0.001, ****p<0.0001.

**Supplemental Figure 4: Quantification of astrocyte and microglia in 4-mo-old WT, *ApoeCh*, 5xFAD, 5xFAD;*ApoeCh* mice. a, c** Representative 20x confocal images of the subiculum stained for dense-core plaque with AmyloGlo (green) and immunolabeled for astrocyte with GFAP (red, **a**) and IBA1 for microglia (red, **c**). Scale bar=100µm. **b, d** Total GFAP+ (**b**) and IBA1+ volume (**d**) in the subiculum per FOV of 4-mo-old WT, *ApoeCh*, 5xFAD, 5xFAD;*ApoeCh* mice. **e, f** Total number of IBA1+ cells per FOV of 4 mo (**e**) and 12 mo (**f**) WT, *ApoeCh*, 5xFAD, 5xFAD;*ApoeCh* mice in the subiculum.

**Supplemental Figure 5: Cell segmentation and annotation in 5xFAD;*ApoeCh* spatial proteomics data. a** Representative images of cell segmentation (teal outline) in cortex, dentate gyrus (i.e., high cell density region), and white matter tracts for 5xFAD;*ApoeCh* spatial proteomics dataset. DAPI nuclear stain is shown in light grey. White dotted line demarcating anatomical landmarks for dentate gyrus and white matter tracts. Scale bar = 50 µm. **b** 12 CELESTA cell types in XY space for all 12 hemibrains (n=3/genotype).

**Supplemental Figure 6: Cell segmentation and annotation in 5xFAD;*ApoeCh* spatial transcriptomics data. a** Representative images of cell segmentation (teal outline) in white matter tracts, CA1, and cortex for 5xFAD;*ApoeCh* spatial transcriptomics dataset. DAPI nuclear stain shown in light grey, and histone marker shown in green. White dotted lines demarcating anatomical landmarks for white matter tracts and CA1 of hippocampus. Scale bar = 50 µm. **b** 33 cell types in XY space for all 12 sections (n=3/genotype).

**Supplemental Figure 7: Distribution of cell types across genotypes in 5xFAD;*ApoeCh* dataset. a** UMAP of cells from 5xFAD;*ApoeCh* spatial transcriptomics dataset split by genotype (5xFAD;*ApoeCh*, 5xFAD, *ApoeCh*, WT). DAM (dark green) and DAA (dark orange) clusters are primarily present in 5xFAD;*ApoeCh* and 5xFAD genotypes, but not *ApoeCh* and WT genotypes. **b** Heatmap of the top five marker genes per major cell type. DGE analysis was performed between each major cell type compared to all other major cell types to identify the top 5 genes expressed for each major cell type. **c** Stacked bar plot of cell counts of all 33 clusters, split by genotype. Clusters are grouped by their major cell type (i.e., ODC 1-5 and OPC are grouped together on the bar plot), and then within each major cell type plotted by size in descending order.

**Supplemental Figure 8: Differential gene expression analysis between 5xFAD vs. WT and 5xFAD;*ApoeCh* vs. 5xFAD across major cell types and all cell types. a** Volcano plots showing DEGs between 5xFAD and WT across each cluster. Because DAMs and DAAs are present in 5xFAD but not in WT mice, DAMs and microglia were grouped together into “All microglia”, and DAAs and astrocytes were grouped together into “All astrocytes” for the 5xFAD vs. WT comparison. Each gene represents a point on the volcano plot. Red and blue colored points represent significantly up- and downregulated genes, respectively (i.e., p_adj_ < 0.05 and absolute average difference > 0.3). Black points represent unsignificant genes (i.e., p_adj_ > 0.05 or absolute average difference < 0.3). Only the top 20 genes are labelled for legibility. **b** Volcano plots showing DEGs between 5xFAD;*ApoeCh* and 5xFAD across each cluster. Dotted red outlines highlight clusters CA1 and L2/L3, which are shown to have the greatest (most negative) DD score in the 5xFAD;*ApoeCh* vs. 5xFAD comparison (Fig. 4i). **c** Volcano plots showing DEGs between 5xFAD and WT across each major cell type. DAMs and microglia are grouped into “All microglia”, and DAAs and astrocytes into “All astrocytes”.

**Supplemental Figure 9: Differential gene expression changes in L2/L3 of cortex and CA1 of hippocampus. a** Examples of high spatial specificity for dentate gyrus, CA1, and L2/L3 clusters in a 5xFAD;*ApoeCh* section. **b** Dot plot showing pseudo-bulk expression of the top DEGs in the 5xFAD;*ApoeCh* vs. 5xFAD comparison across all four genotypes in L2/L3 and CA1 clusters.

**Supplemental Figure 10: Microglial subclustering in 5xFAD;*ApoeCh* dataset. n a** Feature plots for common microglial and DAM marker genes. **b** Example of cell segmentation and cell annotation surrounding an amyloid beta plaque in a 5xFAD;*ApoeCh* brain. (top) White arrows pointing to two cells annotated as amyloid beta plaques. Many cells in this plaque cluster were anucleate (i.e., did not contain histone marker). DAPI nuclear stain (light grey), histone marker (green). (bottom) Cell type annotation for all microglia. Cells annotated as Abeta plaques (red) are surrounded by 5 DAM-2 cells (turquoise), 2 DAM-1 cells (blue), and a Microglia-1 (magenta). **c** Heatmap of the top 10 marker genes for each microglial subcluster. DGE analysis was performed for each microglial subcluster compared to all other microglial subclusters to obtain the top 10 genes expressed in each microglial subcluster.

**Supplemental Figure 11: Behavioral analysis of 5- and 9-mo-old WT, *ApoeCh*, PS19, PS19;*ApoeCh* mice. a, b** Weight of 5-mo-old (**a**) and 9-mo-old (**b**) mice taken at euthanizing day. **c, d** Violin plot of hindlimb clasping score of 4-mo-old (c) and 12-mo-old (d) mice. **e, g** Total time mice spent in the center of open field behavioral assay of 5-mo-old (**e**) and 9-mo-old (**g**) in the 5 min recording time. **f, h** Mean velocity mice traveled in the center of the open field in 5 mins of 5-mo (**f**) and 9-mo-old mice (**h**). **i, j** Total time 5-mo (**i**) and 12-mo-old (**j**) mice spent in open arms of the elevated plus maze behavior test. **k, l** Measurement of plasma NfL level in WT, *ApoeCh*, PS19, and PS19;*ApoeCh* mice at 5- (**k**) and 9-mo (**l**). n=4-10 mice/sex/genotype. Data are represented as mean ± SEM. Two-way ANOVA followed by Tukey’s post hoc tests to examine biologically relevant interactions. Statistical significance is denoted by *p<0.05, **p<0.01, ***p<0.001, ****p<0.0001. # denotes trending significance.

**Supplemental Figure 12: Immunohistochemical analysis of 5-mo-old WT, *ApoeCh*, PS19, PS19;*ApoeCh* mice. a** Representative 20x confocal images of the dentate gyrus (top) and piriform cortex (bottom) immunolabeled with AT8 for phosphorylated tau (green). Scale bar = 100µm. **b, c** Percent coverage of AT8+ area per FOV in the dentate gyrus (b) and piriform cortex (c). **d, f** Representative 20x confocal images of the dentate gyrus immunolabeled with IBA1 for microglia (red, d) and GFAP for astrocytes (blue, f). Scale bar = 100µm. **e, g** Total volume of IBA1+ (**e**) and GFAP+ (**g**) cells per FOV of the dentate gyrus. **h, i** IBA1+ microglia number per FOV of 5-mo-old (h) and 9-mo-old (i) mice in the dentate gyrus. n=4-6 mice/sex/genotype. Data are represented as mean ± SEM. Two-way ANOVA followed by Tukey’s post hoc tests to examine biologically relevant interactions. Statistical significance is denoted by *p<0.05, **p<0.01, ***p<0.001, ****p<0.0001.

**Supplemental Figure 13: Cell segmentation and annotation in PS19;*ApoeCh* spatial proteomics data. a** Representative images of cell segmentation (teal outline) in dentate gyrus, choroid plexus, and cortex for PS19;*ApoeCh* spatial proteomics dataset. DAPI nuclear stain is shown in light grey. White dotted line demarcates anatomical landmarks for dentate gyrus. Scale bar = 50 µm. **b** CELESTA cell types in XY space for 11 hemibrains (n=3/genotype, except n=2 for *ApoeCh*).

**Supplemental Figure 14: Cell segmentation and annotation in PS19;*ApoeCh* spatial transcriptomics data. a** Representative images of cell segmentation (teal outline) in cortex, dentate gyrus, and white matter tracts for PS19;*ApoeCh* spatial transcriptomics dataset. DAPI nuclear stain (light grey), histone marker (green). White dotted line demarcates anatomical landmarks for dentate gyrus, white matter tracts, and CA1 of hippocampus. Scale bar = 50 µm. **b** 40 clusters in XY space for all 11 sections (n=3/genotype, except n=2 for *ApoeCh*).

**Supplemental Figure 15: Distribution of cell types across genotypes in PS19;*ApoeCh* dataset. a** UMAP of cells from PS19;*ApoeCh* spatial transcriptomics dataset split by genotype (PS19;*ApoeCh*, PS19, *ApoeCh*, and WT). DAM (dark green) cluster is present in PS19 genotypes, but not in PS19;*ApoeCh*, *ApoeCh*, or WT genotypes. **b** Heatmap of the top 5 marker genes per major cell type. DGE analysis was performed between each major cell type compared to all other major cell types to identify the top 5 genes expressed for each major cell type. **c** Stacked bar plot of each of the 40 clusters, split by genotype. Clusters are grouped by their major cell type (e.g., Astrocyte 1-3 and DAA are grouped together) and plotted in order from largest to smallest cluster.

**Supplemental Figure 16: Differential gene expression analysis between PS19 vs. WT and PS19;*ApoeCh* vs. PS19 across major cell types and all cell types. a** Volcano plots of DEGs between PS19 and WT for all clusters. Because DAA and DAM clusters are present in very low numbers in WT mice, DAA and astrocytes are combined into “All astrocytes”, and DAM and microglia are combined into “All microglia”. **b** Volcano plots of DEGs between PS19;*ApoeCh* and PS19 for all 40 clusters. **c** Volcano plots of DEGs between PS19 and WT for each major cell type. All astrocytes and all microglia are grouped together.

## DECLARATIONS

### Ethics approval and consent to participate

All experiments involving mice were approved by the UC Irvine Institutional Animal Care and Use Committee and were conducted in compliance with all relevant ethical regulations for animal testing and research. All experiments involving mice comply with the Animal Research: Reporting of *in Vivo* Experiments (ARRIVE-10) guidelines.

### Consent for publication

Not applicable

### Availability of data and materials

Protocols, data, and results are available via the AD Knowledge Portal (https://adknowledgeportal.synapse.org). The AD Knowledge Portal is a platform for accessing data, analyses, and tools generated by the Accelerating Medicines Partnership (AMP-AD) Target Discovery Program and other National Institute on Aging (NIA)-supported programs to enable open-science practices and accelerate translational learning. The data, analyses and tools are shared early in the research cycle without a publication embargo on secondary use. Data is available for general research use according to the following requirements for data access and data attribution (https://adknowledgeportal.org/DataAccess/Instructions). Data can be accessed in an interactive matter at UCI Mouse Mind Explorer (admodelexplorer.org).

The *ApoeCh* model is available from The Jackson Laboratory (Stock # 39301) without restrictions on its use by both academic and commercial users. The content is solely the responsibility of the authors and does not necessarily represent the official view of the National Institutes of Health.

### Authors’ contributions

KMT, NK, AG-A, SK, CM, DC, CDC, SW, SC, AW, K-XS, JAA, and JN performed experiments and analyzed data. KMT, NK, LAH, GRM and KNG analyzed and reviewed all data and wrote the manuscript with contributions from all authors. VS, AJT, FL, GRM, and KNG designed the study, managed the project, and contributed to the manuscript. All authors read and approved the final manuscript.

## Acknowledgments

The authors acknowledge the support of the Transgenic Mouse Facility shared resource supported in part by the National Cancer Institute of the National Institutes of Health under award number (P30CA062203) at the University of California, Irvine.

## REFERENCES

1. Braak, H. and E. Braak, Neuropathological stageing of Alzheimer-related changes. Acta Neuropathol, 1991. 82(4): p. 239–59.

2. Tanzi, R.E., The genetics of Alzheimer disease. Cold Spring Harb Perspect Med, 2012. 2(10).

3. Sherrington, R., et al., Alzheimer’s disease associated with mutations in presenilin 2 is rare and variably penetrant. Hum Mol Genet, 1996. 5(7): p. 985–8.

4. Levy-Lahad, E., et al., A familial Alzheimer’s disease locus on chromosome 1. Science, 1995. 269(5226): p. 970–3.

5. Rogaev, E.I., et al., Familial Alzheimer’s disease in kindreds with missense mutations in a gene on chromosome 1 related to the Alzheimer’s disease type 3 gene. Nature, 1995. 376(6543): p. 775–8.

6. Chartier-Harlin, M.C., et al., Early-onset Alzheimer’s disease caused by mutations at codon 717 of the beta-amyloid precursor protein gene. Nature, 1991. 353(6347): p. 844–6.

7. Goate, A., et al., Segregation of a missense mutation in the amyloid precursor protein gene with familial Alzheimer’s disease. Nature, 1991. 349(6311): p. 704–6.

8. Jansen, I.E., et al., Genome-wide meta-analysis identifies new loci and functional pathways influencing Alzheimer’s disease risk. Nat Genet, 2019. 51(3): p. 404–413.

9. Serrano-Pozo, A., et al., APOEepsilon2 is associated with milder clinical and pathological Alzheimer disease. Ann Neurol, 2015. 77(6): p. 917–29.

10. Barger, S.W. and M.P. Mattson, Isoform-specific modulation by apolipoprotein E of the activities of secreted beta-amyloid precursor protein. J Neurochem, 1997. 69(1): p. 60–7.

11. Huang, Y.A., et al., ApoE2, ApoE3, and ApoE4 Differentially Stimulate APP Transcription and Abeta Secretion. Cell, 2017. 168(3): p. 427–441 e21.

12. Arboleda-Velasquez, J.F., et al., Resistance to autosomal dominant Alzheimer’s disease in an APOE3 Christchurch homozygote: a case report. Nat Med, 2019. 25(11): p. 1680–1683.

13. Fuentealba, R.A., et al., Low-density lipoprotein receptor-related protein 1 (LRP1) mediates neuronal Abeta42 uptake and lysosomal trafficking. PLoS One, 2010. 5(7): p. e11884.

14. Kanekiyo, T., et al., Heparan sulphate proteoglycan and the low-density lipoprotein receptor-related protein 1 constitute major pathways for neuronal amyloid-beta uptake. J Neurosci, 2011. 31(5): p. 1644–51.

15. Kanekiyo, T. and G. Bu, The low-density lipoprotein receptor-related protein 1 and amyloid-beta clearance in Alzheimer’s disease. Front Aging Neurosci, 2014. 6: p. 93.

16. Kanekiyo, T., et al., Neuronal clearance of amyloid-beta by endocytic receptor LRP1. J Neurosci, 2013. 33(49): p. 19276–83.

17. Rauch, J.N., et al., LRP1 is a master regulator of tau uptake and spread. Nature, 2020. 580(7803): p. 381-385.

18. Kloske, C.M. and D.M. Wilcock, The Important Interface Between Apolipoprotein E and Neuroinflammation in Alzheimer’s Disease. Front Immunol, 2020. 11: p. 754.

19. Rebeck, G.W., et al., Apolipoprotein E in sporadic Alzheimer’s disease: allelic variation and receptor interactions. Neuron, 1993. 11(4): p. 575–80.

20. Forner, S., et al., Systematic phenotyping and characterization of the 5xFAD mouse model of Alzheimer’s disease. Sci Data, 2021. 8(1): p. 270.

21. Guyenet, S.J., et al., A simple composite phenotype scoring system for evaluating mouse models of cerebellar ataxia. J Vis Exp, 2010(39).

22. Javonillo, D.I., et al., Systematic Phenotyping and Characterization of the 3xTg-AD Mouse Model of Alzheimer’s Disease. Front Neurosci, 2021. 15: p. 785276.

23. Tran, K.M., et al., A Trem2(R47H) mouse model without cryptic splicing drives age- and disease-dependent tissue damage and synaptic loss in response to plaques. Mol Neurodegener, 2023. 18(1): p. 12.

24. Shi, Y., et al., ApoE4 markedly exacerbates tau-mediated neurodegeneration in a mouse model of tauopathy. Nature, 2017. 549(7673): p. 523-527.

25. Yanamandra, K., et al., Anti-tau antibodies that block tau aggregate seeding in vitro markedly decrease pathology and improve cognition in vivo. Neuron, 2013. 80(2): p. 402–414.

26. Hao, Y., et al., Dictionary learning for integrative, multimodal and scalable single-cell analysis. Nat Biotechnol, 2024. 42(2): p. 293–304.

27. Finak, G., et al., MAST: a flexible statistical framework for assessing transcriptional changes and characterizing heterogeneity in single-cell RNA sequencing data. Genome Biol, 2015. 16: p. 278.

28. Wickham, H., ggplot2: Elegant Graphics for Data Analysis. 2016: Springer Publishing Company, Incorporated.

29. Zhang, W., et al., Identification of cell types in multiplexed in situ images by combining protein expression and spatial information using CELESTA. Nat Methods, 2022. 19(6): p. 759–769.

30. Fortea, J., et al., APOE4 homozygozity represents a distinct genetic form of Alzheimer’s disease. Nat Med, 2024. 30(5): p. 1284–1291.

31. Bae, S., J. Park, and J.S. Kim, Cas-OFFinder: a fast and versatile algorithm that searches for potential off-target sites of Cas9 RNA-guided endonucleases. Bioinformatics, 2014. 30(10): p. 1473–5.

32. Chen, Y., et al., APOE3ch alters microglial response and suppresses Abeta-induced tau seeding and spread. Cell, 2024. 187(2): p. 428–445 e20.

33. Vidal, M., et al., Tissue-specific control elements of the Thy-1 gene. EMBO J, 1990. 9(3): p. 833–40.

34. Moechars, D., et al., Expression in brain of amyloid precursor protein mutated in the alpha-secretase site causes disturbed behavior, neuronal degeneration and premature death in transgenic mice. EMBO J, 1996. 15(6): p. 1265–74.

35. Yoshiyama, Y., et al., Synapse loss and microglial activation precede tangles in a P301S tauopathy mouse model. Neuron, 2007. 53(3): p. 337–51.

36. Takeuchi, H., et al., P301S mutant human tau transgenic mice manifest early symptoms of human tauopathies with dementia and altered sensorimotor gating. PLoS One, 2011. 6(6): p. e21050.

37. Maruyama, M., et al., Imaging of tau pathology in a tauopathy mouse model and in Alzheimer patients compared to normal controls. Neuron, 2013. 79(6): p. 1094–108.

38. Condello, C., et al., Microglia constitute a barrier that prevents neurotoxic protofibrillar Abeta42 hotspots around plaques. Nat Commun, 2015. 6: p. 6176.

39. Bacioglu, M., et al., Neurofilament Light Chain in Blood and CSF as Marker of Disease Progression in Mouse Models and in Neurodegenerative Diseases. Neuron, 2016. 91(1): p. 56–66.

40. Piwecka, M., N. Rajewsky, and A. Rybak-Wolf, Single-cell and spatial transcriptomics: deciphering brain complexity in health and disease. Nat Rev Neurol, 2023. 19(6): p. 346–362.

41. Williams, C.G., et al., An introduction to spatial transcriptomics for biomedical research. Genome Med, 2022. 14(1): p. 68.

42. Jovic, D., et al., Single-cell RNA sequencing technologies and applications: A brief overview. Clin Transl Med, 2022. 12(3): p. e694.

43. Hammond, T.R., et al., Single-Cell RNA Sequencing of Microglia throughout the Mouse Lifespan and in the Injured Brain Reveals Complex Cell-State Changes. Immunity, 2019. 50(1): p. 253–271 e6.

44. Marsh, S.E., et al., Dissection of artifactual and confounding glial signatures by single-cell sequencing of mouse and human brain. Nat Neurosci, 2022. 25(3): p. 306–316.

45. Keren-Shaul, H., et al., A Unique Microglia Type Associated with Restricting Development of Alzheimer’s Disease. Cell, 2017. 169(7): p. 1276–1290 e17.

46. Krasemann, S., et al., The TREM2-APOE Pathway Drives the Transcriptional Phenotype of Dysfunctional Microglia in Neurodegenerative Diseases. Immunity, 2017. 47(3): p. 566–581 e9.

47. Deczkowska, A., et al., Disease-Associated Microglia: A Universal Immune Sensor of Neurodegeneration. Cell, 2018. 173(5): p. 1073–1081.

48. Gallo, F.T., et al., Immediate Early Genes, Memory and Psychiatric Disorders: Focus on c-Fos, Egr1 and Arc. Front Behav Neurosci, 2018. 12: p. 79.

49. Sheng, X.F., et al., Long non-coding RNA MALAT1 modulate cell migration, proliferation and apoptosis by sponging microRNA-146a to regulate CXCR4 expression in acute myeloid leukemia. Hematology, 2021. 26(1): p. 43–52.

50. Nakaya, N., et al., Olfactomedin 1 interacts with the Nogo A receptor complex to regulate axon growth. J Biol Chem, 2012. 287(44): p. 37171–84.

51. Wang, Y., et al., m6A demethylase FTO induces NELL2 expression by inhibiting E2F1 m6A modification leading to metastasis of non-small cell lung cancer. Mol Ther Oncolytics, 2021. 21: p. 367–376.

52. Huang, Q., et al., SNAP25 Inhibits Glioma Progression by Regulating Synapse Plasticity via GLS-Mediated Glutaminolysis. Front Oncol, 2021. 11: p. 698835.

53. Shi, Y. and D.M. Holtzman, Interplay between innate immunity and Alzheimer disease: APOE and TREM2 in the spotlight. Nat Rev Immunol, 2018. 18(12): p. 759–772.

54. Yin, Z., et al., APOE4 impairs the microglial response in Alzheimer’s disease by inducing TGFbeta-mediated checkpoints. Nat Immunol, 2023. 24(11): p. 1839–1853.

55. Liu, C.C., et al., Cell-autonomous effects of APOE4 in restricting microglial response in brain homeostasis and Alzheimer’s disease. Nat Immunol, 2023. 24(11): p. 1854–1866.

56. Goedert, M., R. Jakes, and E. Vanmechelen, Monoclonal antibody AT8 recognises tau protein phosphorylated at both serine 202 and threonine 205. Neurosci Lett, 1995. 189(3): p. 167–9.

57. Shi, Y., et al., Microglia drive APOE-dependent neurodegeneration in a tauopathy mouse model. J Exp Med, 2019. 216(11): p. 2546–2561.

58. Gratuze, M., et al., Impact of TREM2R47H variant on tau pathology-induced gliosis and neurodegeneration. J Clin Invest, 2020. 130(9): p. 4954–4968.

59. Zamanillo, D., et al., Importance of AMPA receptors for hippocampal synaptic plasticity but not for spatial learning. Science, 1999. 284(5421): p. 1805–11.

60. Kutsuwada, T., et al., Impairment of suckling response, trigeminal neuronal pattern formation, and hippocampal LTD in NMDA receptor epsilon 2 subunit mutant mice. Neuron, 1996. 16(2): p. 333–44.

61. Brigman, J.L., et al., Loss of GluN2B-containing NMDA receptors in CA1 hippocampus and cortex impairs long-term depression, reduces dendritic spine density, and disrupts learning. J Neurosci, 2010. 30(13): p. 4590–600.

62. Soderling, S.H., et al., A WAVE-1 and WRP signaling complex regulates spine density, synaptic plasticity, and memory. J Neurosci, 2007. 27(2): p. 355–65.

63. Nelson, M.R., et al., The APOE-R136S mutation protects against APOE4-driven Tau pathology, neurodegeneration and neuroinflammation. Nat Neurosci, 2023. 26(12): p. 2104–2121.

64. Naguib, S., et al., APOE3 R136S mutation confers resilience against tau pathology via cGAS-STING-IFN inhibition. bioRxiv, 2024: p. 2024.04.25.591140.

65. Choi, I., et al., Autophagy enables microglia to engage amyloid plaques and prevents microglial senescence. Nat Cell Biol, 2023. 25(7): p. 963–974.

66. Lalazar, A., et al., Site-specific mutagenesis of human apolipoprotein E. Receptor binding activity of variants with single amino acid substitutions. J Biol Chem, 1988. 263(8): p. 3542–5.

67. Bailey, C.C., L.B. DeVaux, and M. Farzan, The Triggering Receptor Expressed on Myeloid Cells 2 Binds Apolipoprotein E. J Biol Chem, 2015. 290(43): p. 26033–42.

68. Atagi, Y., et al., Apolipoprotein E Is a Ligand for Triggering Receptor Expressed on Myeloid Cells 2 (TREM2). J Biol Chem, 2015. 290(43): p. 26043–50.

69. Jendresen, C., et al., The Alzheimer’s disease risk factors apolipoprotein E and TREM2 are linked in a receptor signaling pathway. J Neuroinflammation, 2017. 14(1): p. 59.

70. Yeh, F.L., et al., TREM2 Binds to Apolipoproteins, Including APOE and CLU/APOJ, and Thereby Facilitates Uptake of Amyloid-Beta by Microglia. Neuron, 2016. 91(2): p. 328–40.

71. Mahley, R.W., Y. Huang, and S.C. Rall, Jr., Pathogenesis of type III hyperlipoproteinemia (dysbetalipoproteinemia). Questions, quandaries, and paradoxes. J Lipid Res, 1999. 40(11): p. 1933–49.

72. Holmes, B.B., et al., Heparan sulfate proteoglycans mediate internalization and propagation of specific proteopathic seeds. Proc Natl Acad Sci U S A, 2013. 110(33): p. E3138–47.

73. Shi, Y., et al., Overexpressing low-density lipoprotein receptor reduces tau-associated neurodegeneration in relation to apoE-linked mechanisms. Neuron, 2021. 109(15): p. 2413–2426 e7.

74. Liddelow, S.A., et al., Neurotoxic reactive astrocytes are induced by activated microglia. Nature, 2017. 541(7638): p. 481–487.

75. Chen, X., et al., Microglia-mediated T cell infiltration drives neurodegeneration in tauopathy. Nature, 2023. 615(7953): p. 668–677.

76. Gao, C., et al., Microglia in neurodegenerative diseases: mechanism and potential therapeutic targets. Signal Transduct Target Ther, 2023. 8(1): p. 359.

77. Hansen, D.V., J.E. Hanson, and M. Sheng, Microglia in Alzheimer’s disease. J Cell Biol, 2018. 217(2): p. 459–472.

78. Guan, Y.H., et al., The role of microglia in Alzheimer’s disease and progress of treatment. Ibrain, 2022. 8(1): p. 37–47.

79. Miao, J., et al., Microglia in Alzheimer’s disease: pathogenesis, mechanisms, and therapeutic potentials. Front Aging Neurosci, 2023. 15: p. 1201982.

80. Mosher, K.I. and T. Wyss-Coray, Microglial dysfunction in brain aging and Alzheimer’s disease. Biochem Pharmacol, 2014. 88(4): p. 594–604.

81. Bellenguez, C., et al., New insights into the genetic etiology of Alzheimer’s disease and related dementias. Nat Genet, 2022. 54(4): p. 412–436.

82. Efthymiou, A.G. and A.M. Goate, Late onset Alzheimer’s disease genetics implicates microglial pathways in disease risk. Mol Neurodegener, 2017. 12(1): p. 43.

83. Rajendran, L. and R.C. Paolicelli, Microglia-Mediated Synapse Loss in Alzheimer’s Disease. J Neurosci, 2018. 38(12): p. 2911–2919.

84. Zhan, Y., et al., Deficient neuron-microglia signaling results in impaired functional brain connectivity and social behavior. Nat Neurosci, 2014. 17(3): p. 400–6.

85. Paolicelli, R.C., et al., Synaptic pruning by microglia is necessary for normal brain development. Science, 2011. 333(6048): p. 1456–8.

86. Fassler, M., C. Benaim, and J. George, TREM2 Agonism with a Monoclonal Antibody Attenuates Tau Pathology and Neurodegeneration. Cells, 2023. 12(11).

87. Zhao, P., et al., A tetravalent TREM2 agonistic antibody reduced amyloid pathology in a mouse model of Alzheimer’s disease. Sci Transl Med, 2022. 14(661): p. eabq0095.

88. Asai, H., et al., Depletion of microglia and inhibition of exosome synthesis halt tau propagation. Nat Neurosci, 2015. 18(11): p. 1584–93.

89. Wang, C., et al., Microglial NF-kappaB drives tau spreading and toxicity in a mouse model of tauopathy. Nat Commun, 2022. 13(1): p. 1969.

90. Yoshiyama, Y., et al., Anti-inflammatory action of donepezil ameliorates tau pathology, synaptic loss, and neurodegeneration in a tauopathy mouse model. J Alzheimers Dis, 2010. 22(1): p. 295–306.

