## Supplementary Figures for "APOE Christchurch enhances a disease-associated microglial response to plaque but suppresses response to tau pathology"

#### Supplemental Table 1 - Location of potential off-target sites for crRNA TMF1648 on mouse chromosome 7

| Genomic DNA (11 base seed) | Position | Strand | Mismatches (in seed) | Nearest gene | Off-target code | Analyzed ? |
|---|---|---|---|---|---|---|
| GCACAGAGGAGATACGGGCG\|CGG | 19696877 | - | 0 | ***Apoe* exon 4 - correct target** | | |
| GCACAGAG**c**AGAT**ga**GGG**a**G\|GGG | 45859181 | - | 4 (3) | *Grin2d* - intron 2 | **A** | **YES** |
| G**g**ACA**a**AGGAGATAC**c**GGC**c**\|TGG | 51963361 | - | 4 (2) | *Gas2* - intron 7 | **B** | **YES** |
| **a**CA**t**AGAGGA**a**ATA**a**GGGCG\|AGG | 70552648 | - | 4 (2) | *Gm35842* lncRNA | **C** | **YES** |
| GC**g**CAGAGGAGA**cg**CGG**c**CG\|GGG | 81859324 | + | 4 (3) | *Tm6sf1 - 5' UTR* | **D** | **YES** |
| **a**CACAGAG**t**AGATAC**t**GG**t**G\|TGG | 88598839 | + | 4 (2) | intergenic - non-conserved | **E** | no |
| G**t**ACAGAGGAGATAC**t**GG**g**G\|TGG | 100149385 | - | 3 (2) | intergenic - non-conserved | **F** | no |
| GCACAGAGGAGAT**gg**G**t**G**g**G\|GGG | 122836707 | - | 4 (4) | intergenic - non-conserved | **G** | no |
| G**g**A**g**AGAGGAGATA**g**GGGC**t**\|AGG | 125326739 | + | 4 (2) | *4933440M02Rik* | **H** | **YES** |
| **a**CACAGAGGAGA**a**A**g**GGG**a**G\|GGG | 132361823 | + | 4 (3) | *Fgfr2*-217 - intron 2 (2.9Mb) | **I** | **YES** |
| **c**CACAGAGGAGA**ac**CGGGC**a**\|AGG | 136762831 | + | 4 (3) | intergenic - non-conserved | **J** | no |
| GCACA**c**AGGAG**c**TAC**a**GGC**a**\|AGG | 137604947 | - | 4 (3) | intergenic - non-conserved | **K** | no |
| **t**CACAGAGGAGA**c**A**g**GGG**a**G\|GGG | 137681077 | - | 4 (3) | intergenic - non-conserved | **L** | no |

**Supplementary Table 2: Primers for PCR amplification and sequencing for off-target analysis.** Forward (For) and reverse (Rev) primer sequences are listed for each potential off-target site. The off-target code corresponds to the panels in **Supplementary Fig. 1**.

| **Off-target code** | **Primer Name** | **Primer Sequence (5' - 3')** | **Product size** | **Chromosome 18 locus** |
|---|---|---|---|---|
| A | A For | AACCCATCTCTCATCTTGAACC | 616 bp | *Grin2d*, intron 2 |
| | A Rev | ATGGAGCTGAGGCTAGAAGAC | | |
| B | B For | CACAGACTAACGTCCTTAGCAC | 945 bp | *Gas2*, intron 7 |
| | B Rev | CGTATCACCAAGGGAGCTATG | | |
| C | C For | CCTCACCAACTTTCCTCTACAC | 442 b | *Gm35842*, lncRNA |
| | C Rev | CAAACTCTCAGGTGGCTGTAAC | | |
| D | D For | CCATTAAAGCAAGCGTCCTG | 467 bp | *Tm6sf1*, 5' UTR |
| | D Rev | GAAGCTACCCAGTCCCAAAG | | |
| H | H For | AGTGTCCACCTATCCTTTATCG | 1044 bp | *4933440M02Rik* , lncRNA |
| | H Rev | TTTCTGGCTCTCAGCACTAC | | |
| I | I For | AGTCCGTACACAAACCCAAG | 829 bp | *Fgfr2-217* - intron 2 (2.9Mb) |
| | I Rev | GCAGCATGGTGGATGAATTTG | | |

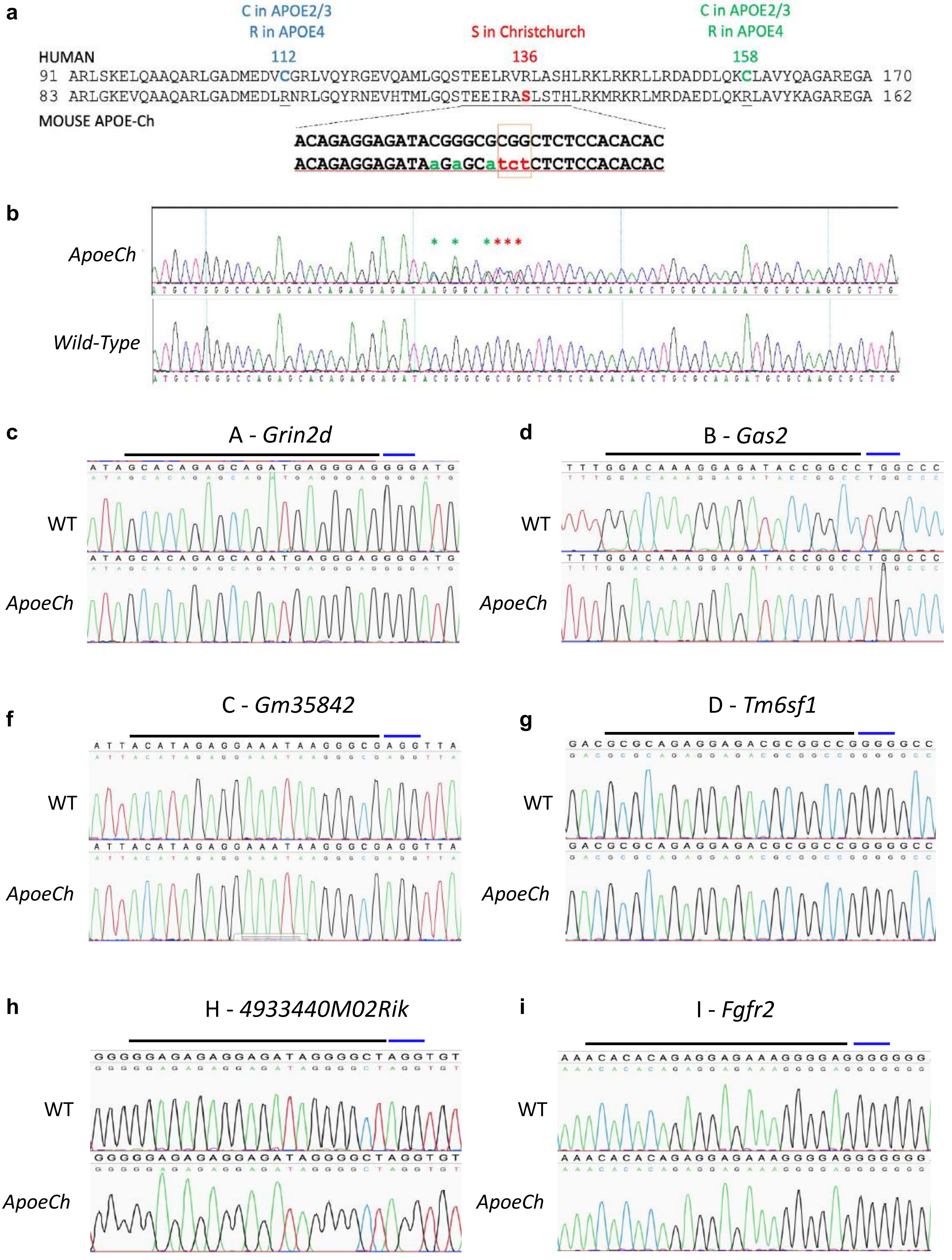


**Supplemental Figure 1**

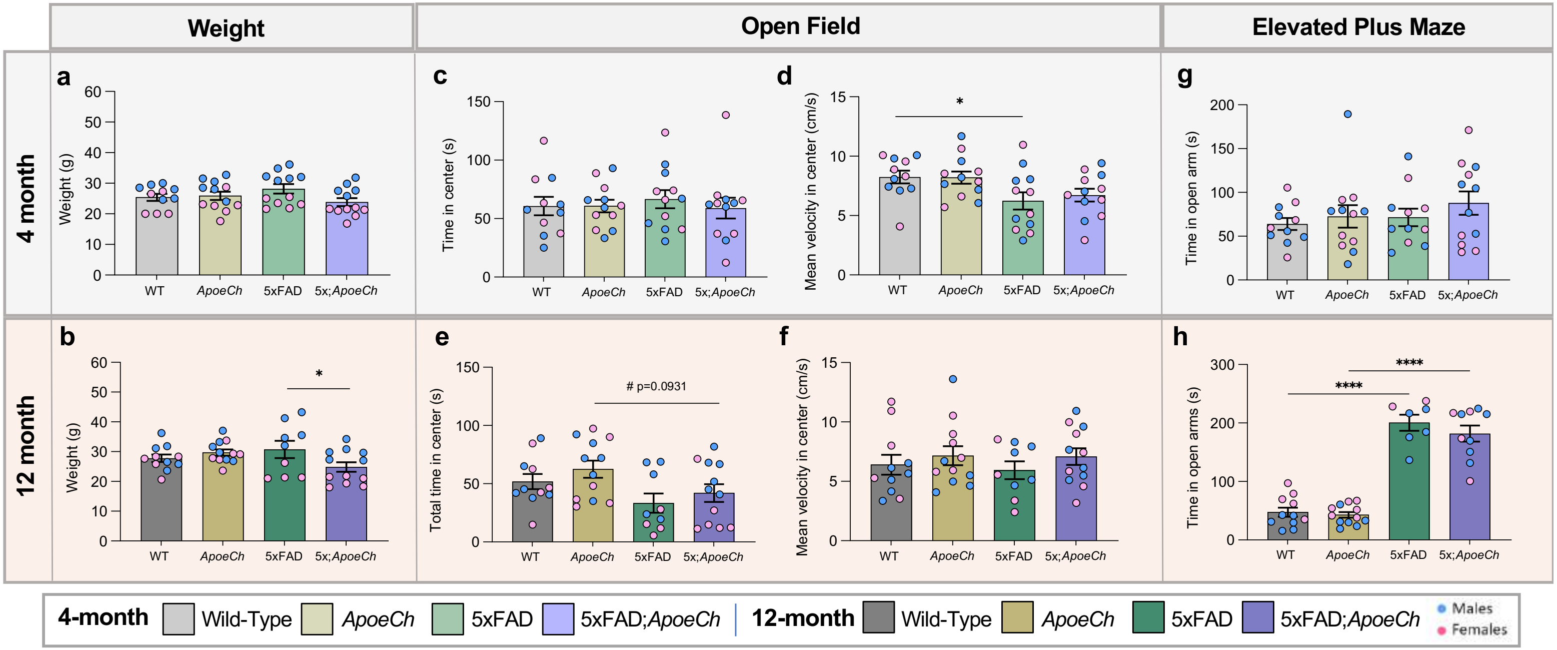


Supplemental Figure 2

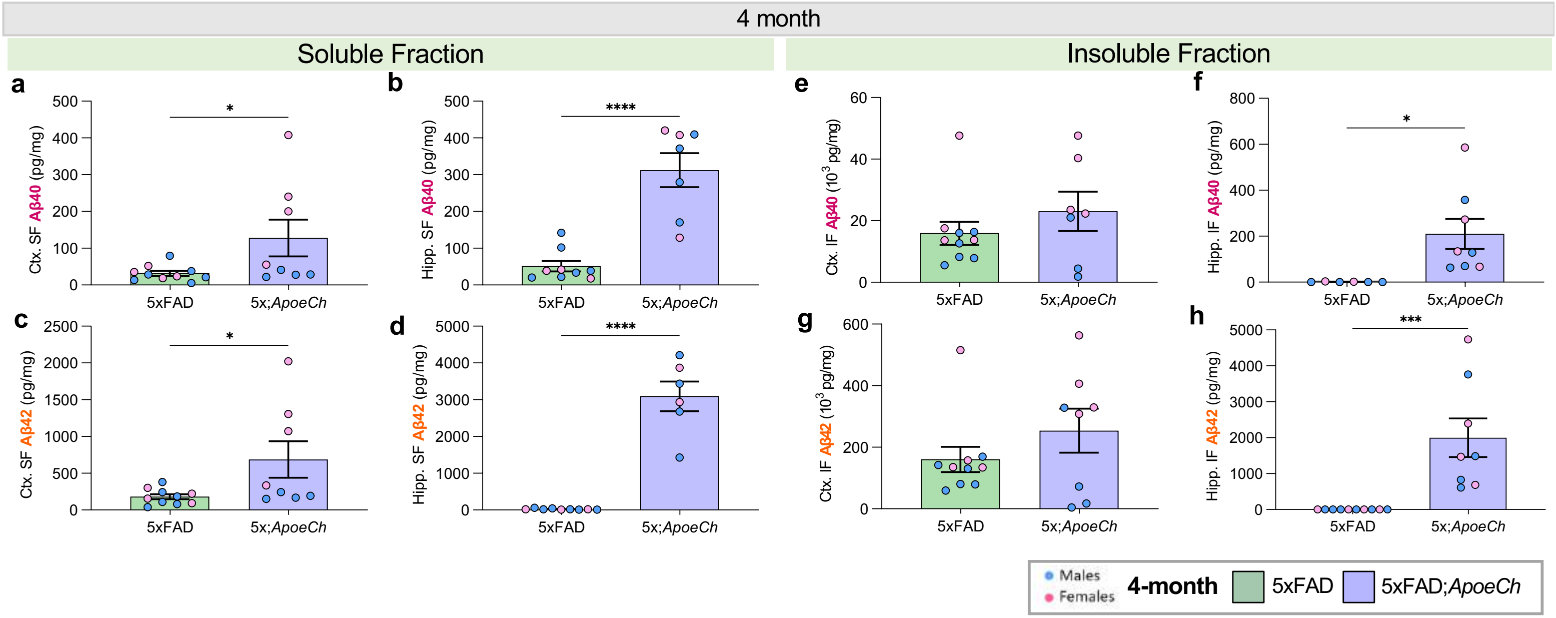


**Supplemental Figure 3**

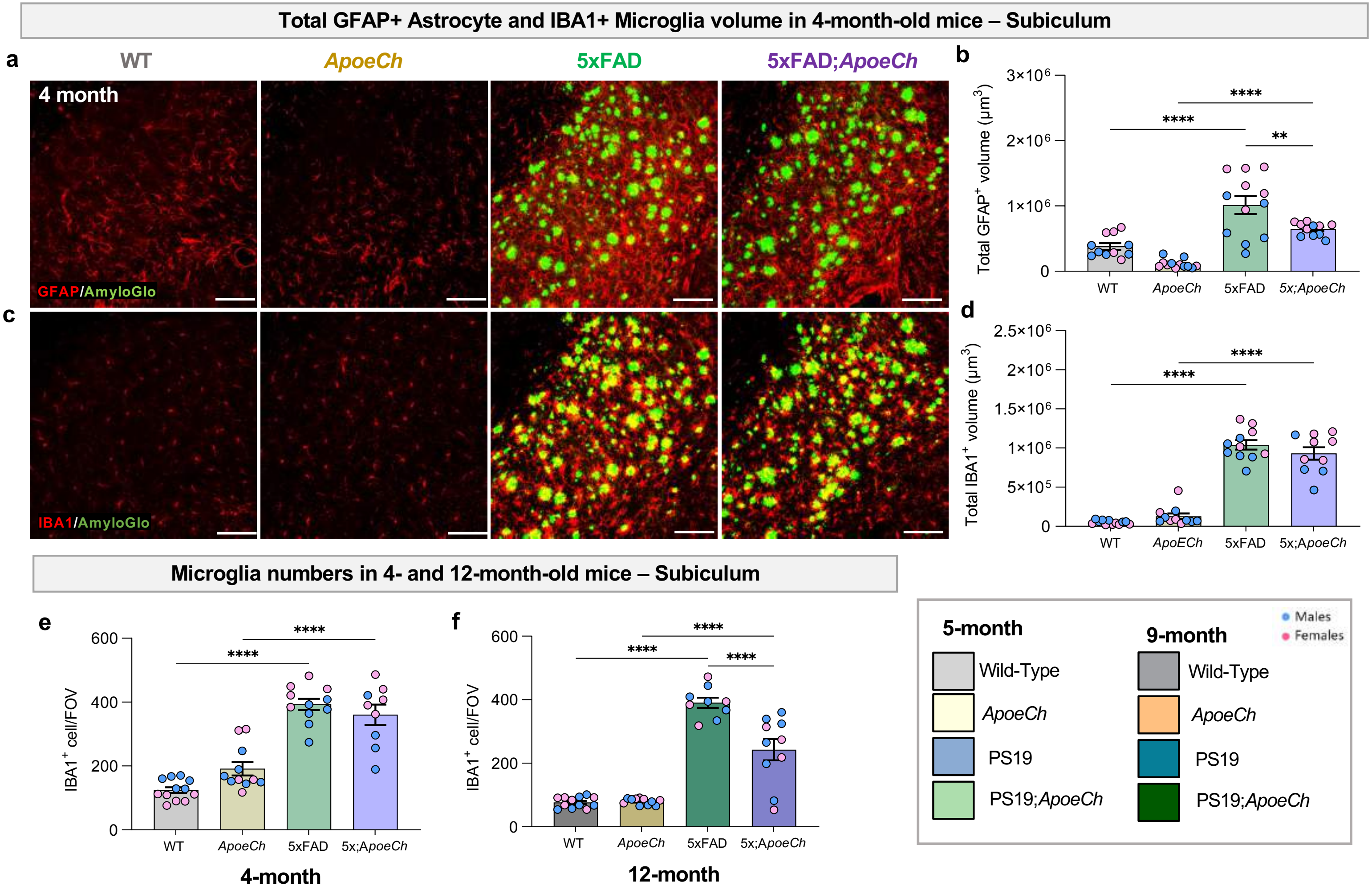


**Supplemental Figure 4**

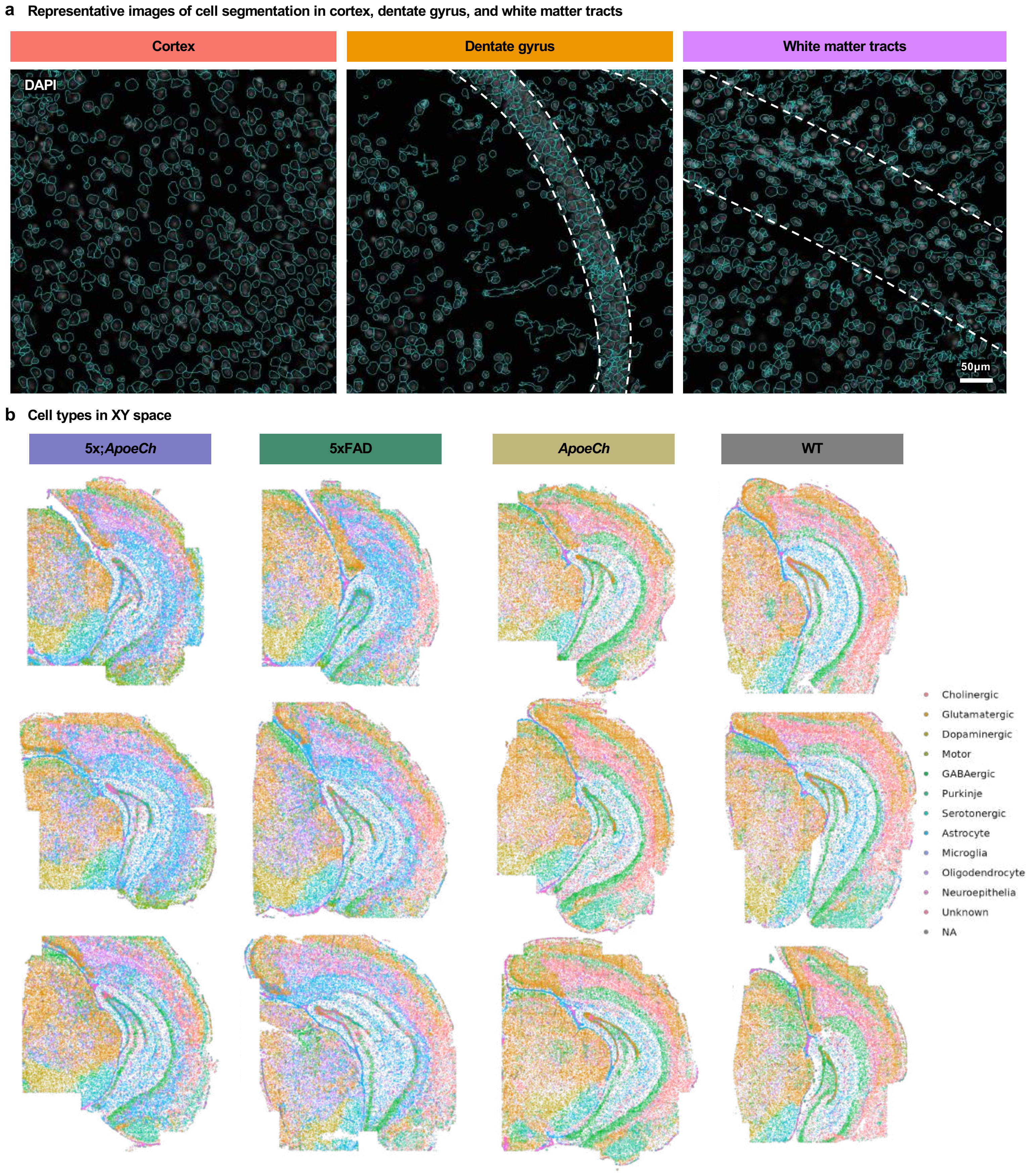


**Supplemental Figure 5**

**a** Representative images of cell segmentation in white matter tracts, CA1, and cortex

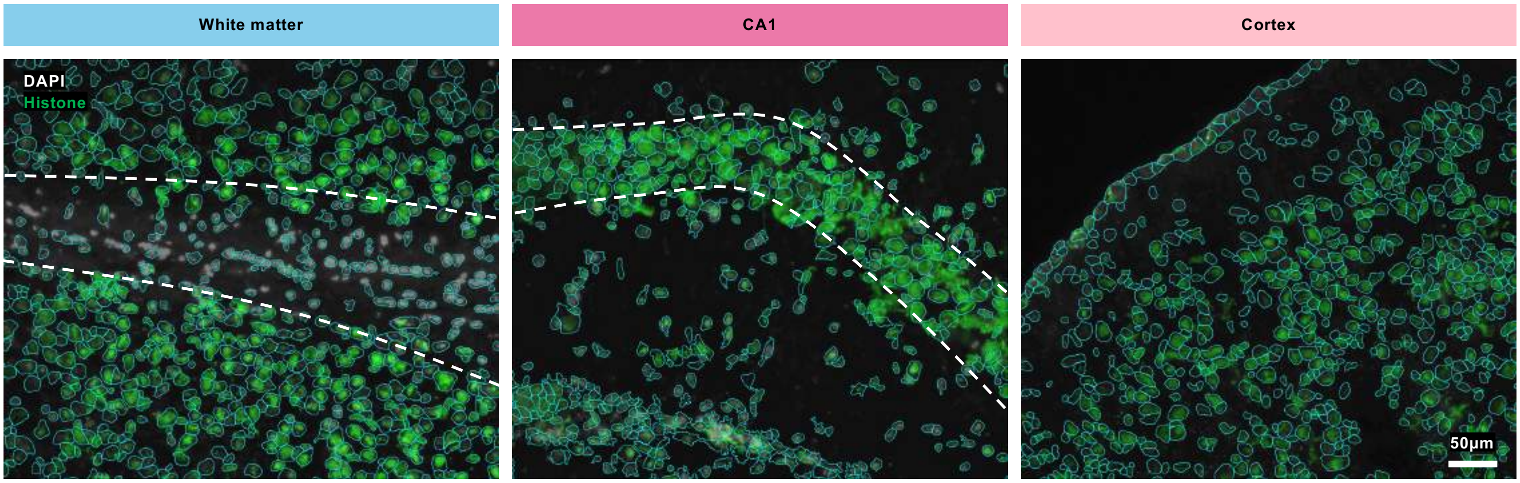


**b** Cell types in XY space

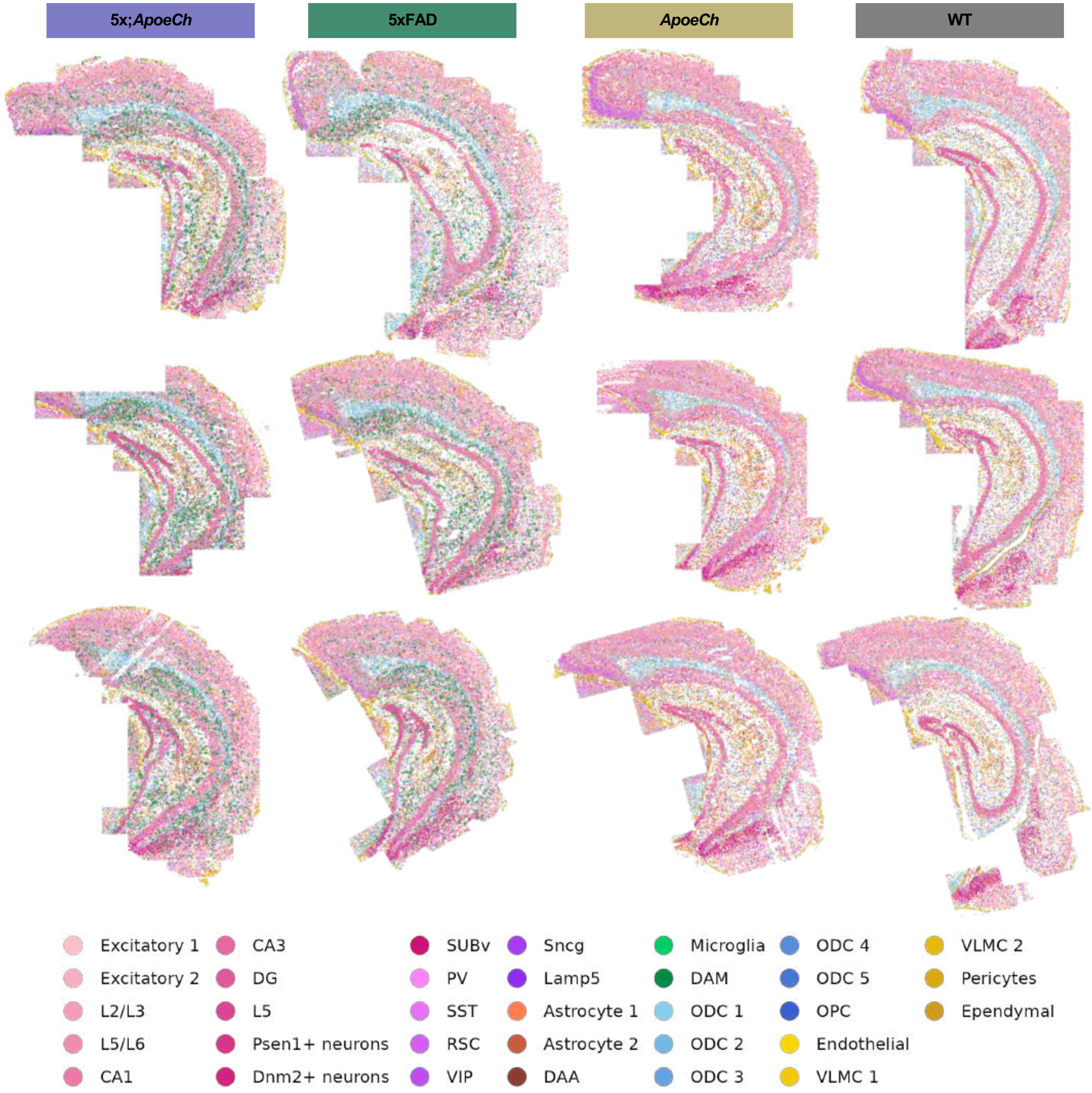


**Supplemental Figure 6**

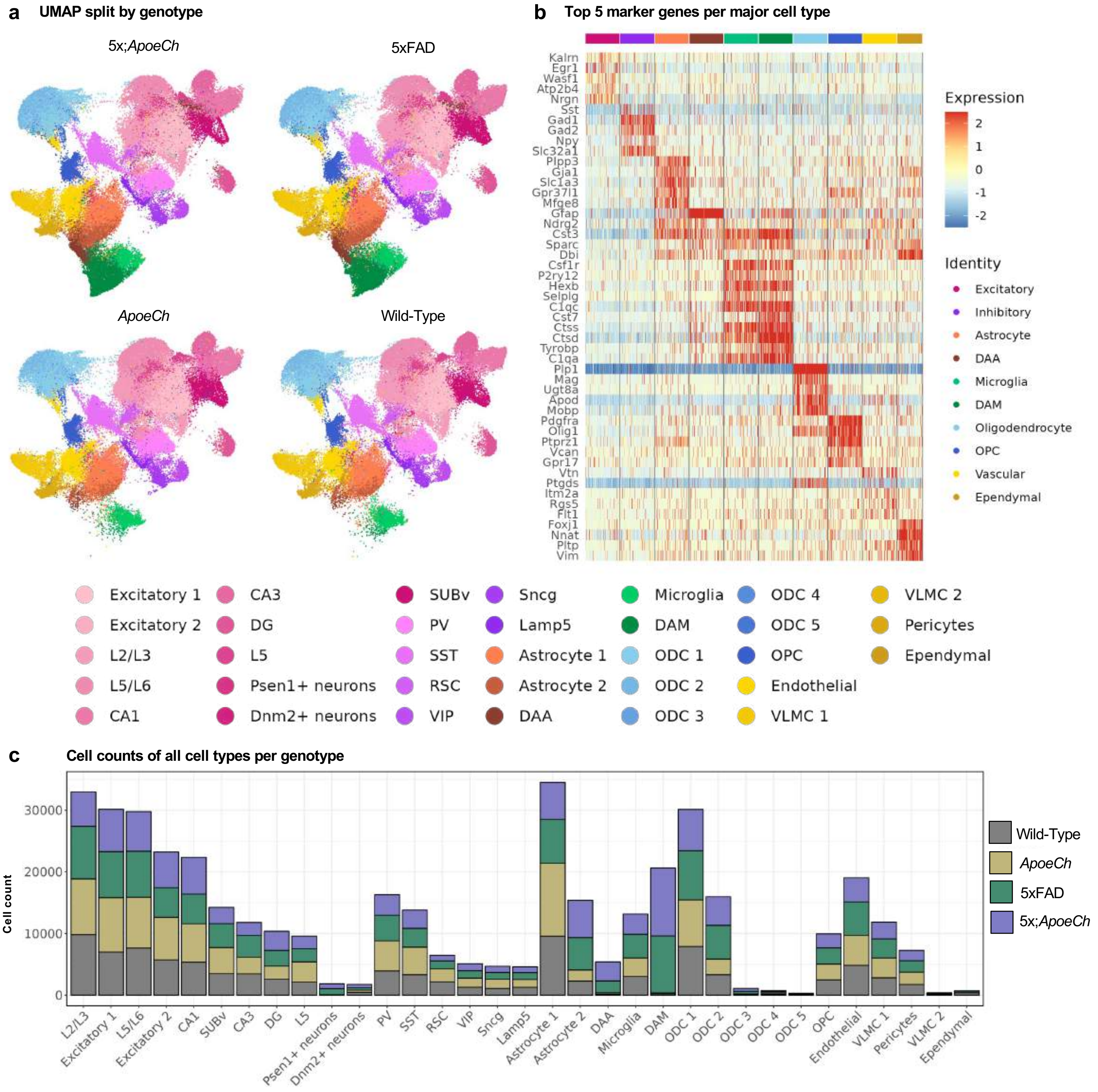


**Supplemental Figure 7**

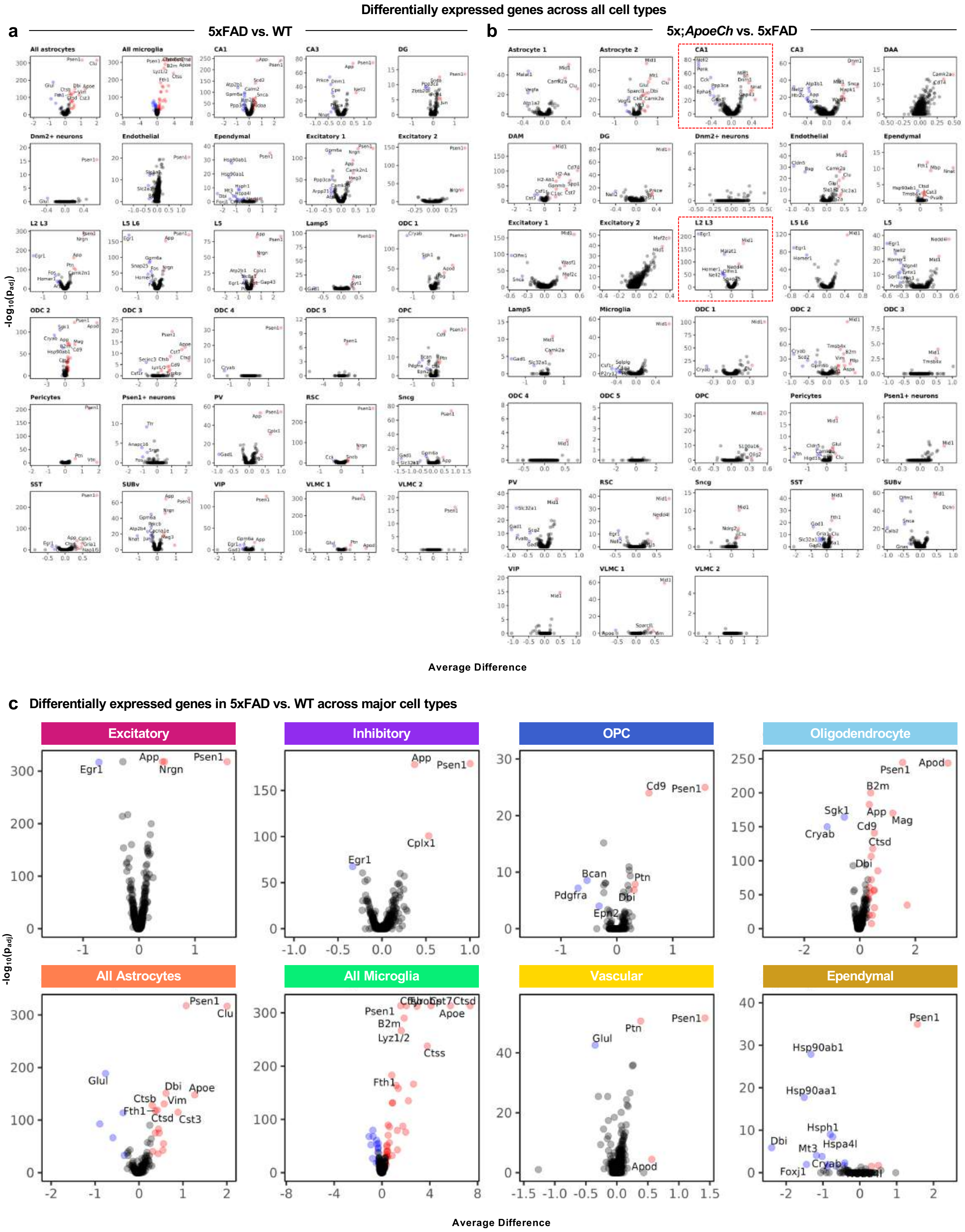


Supplemental Figure 8

#### L2/L3 and CA1 gene expression changes

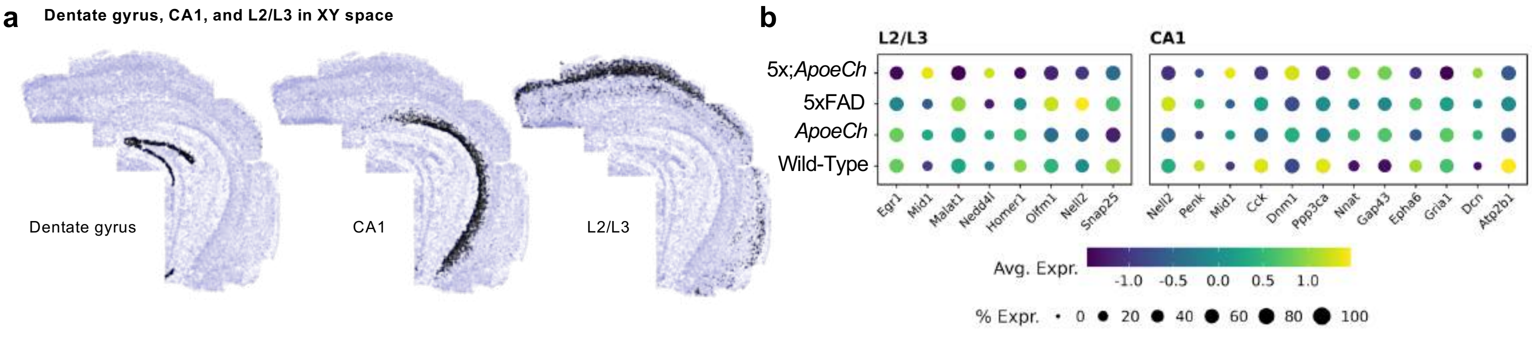


**Supplemental Figure 9**

### Microglial subclustering

**a** Feature plots for microglial and DAM marker genes

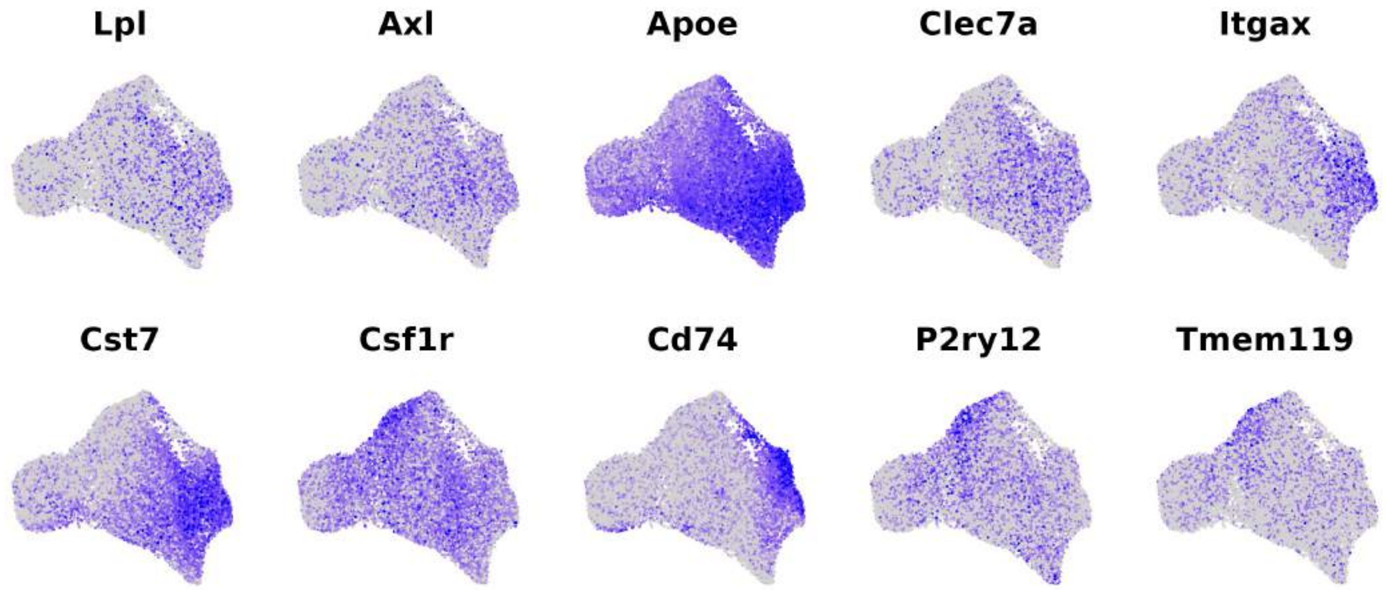


**b** Cell segmentation of plaque surrounded by microglia

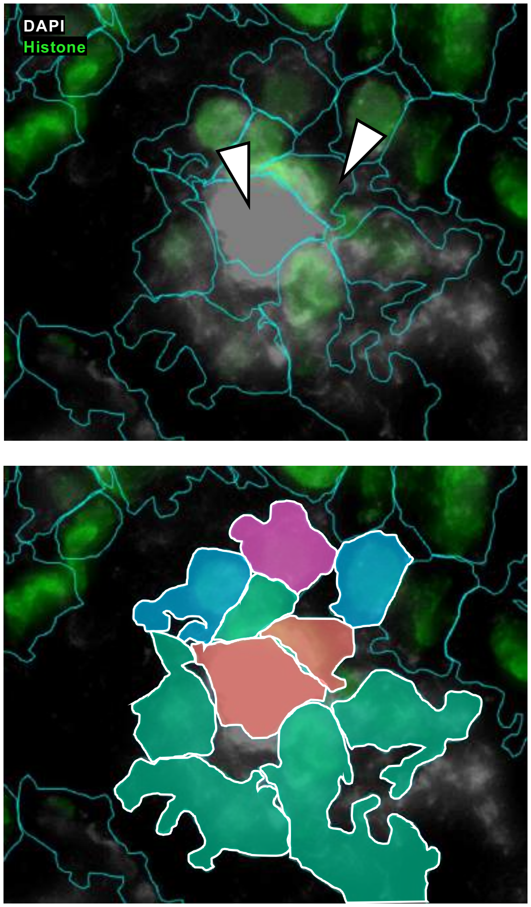


**c** Heatmap of the top 10 marker genes in each microglial subcluster

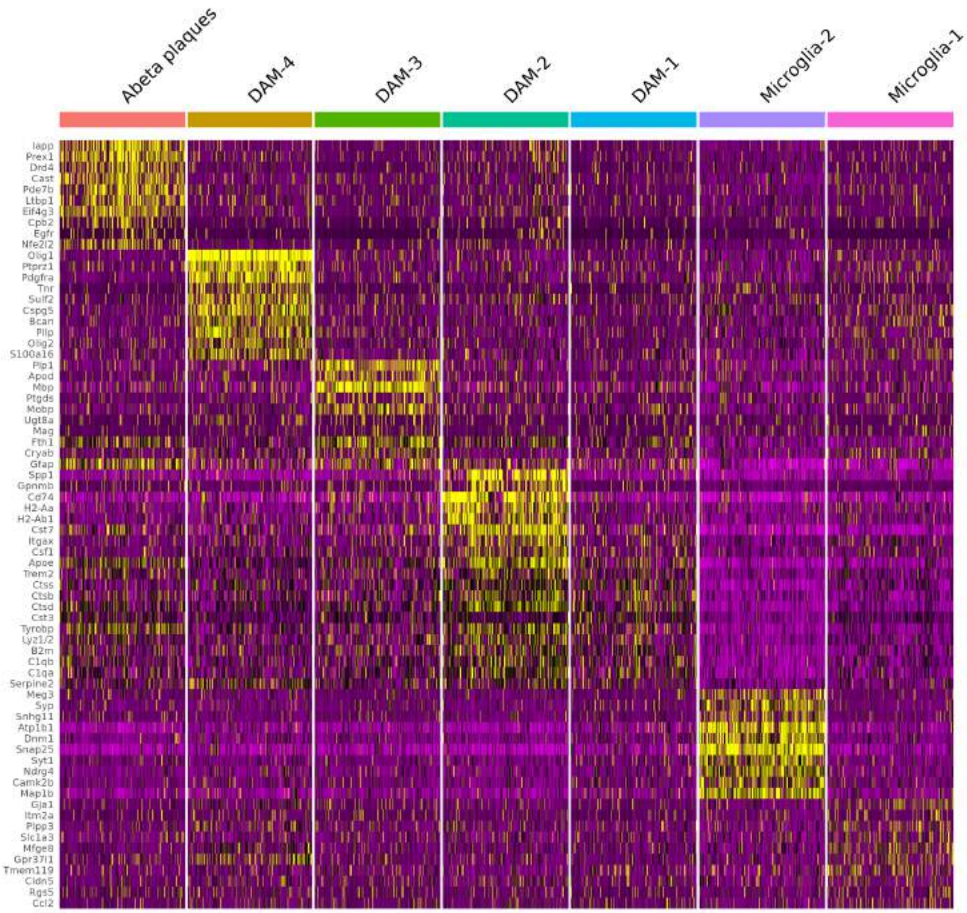


**Supplemental Figure 10**

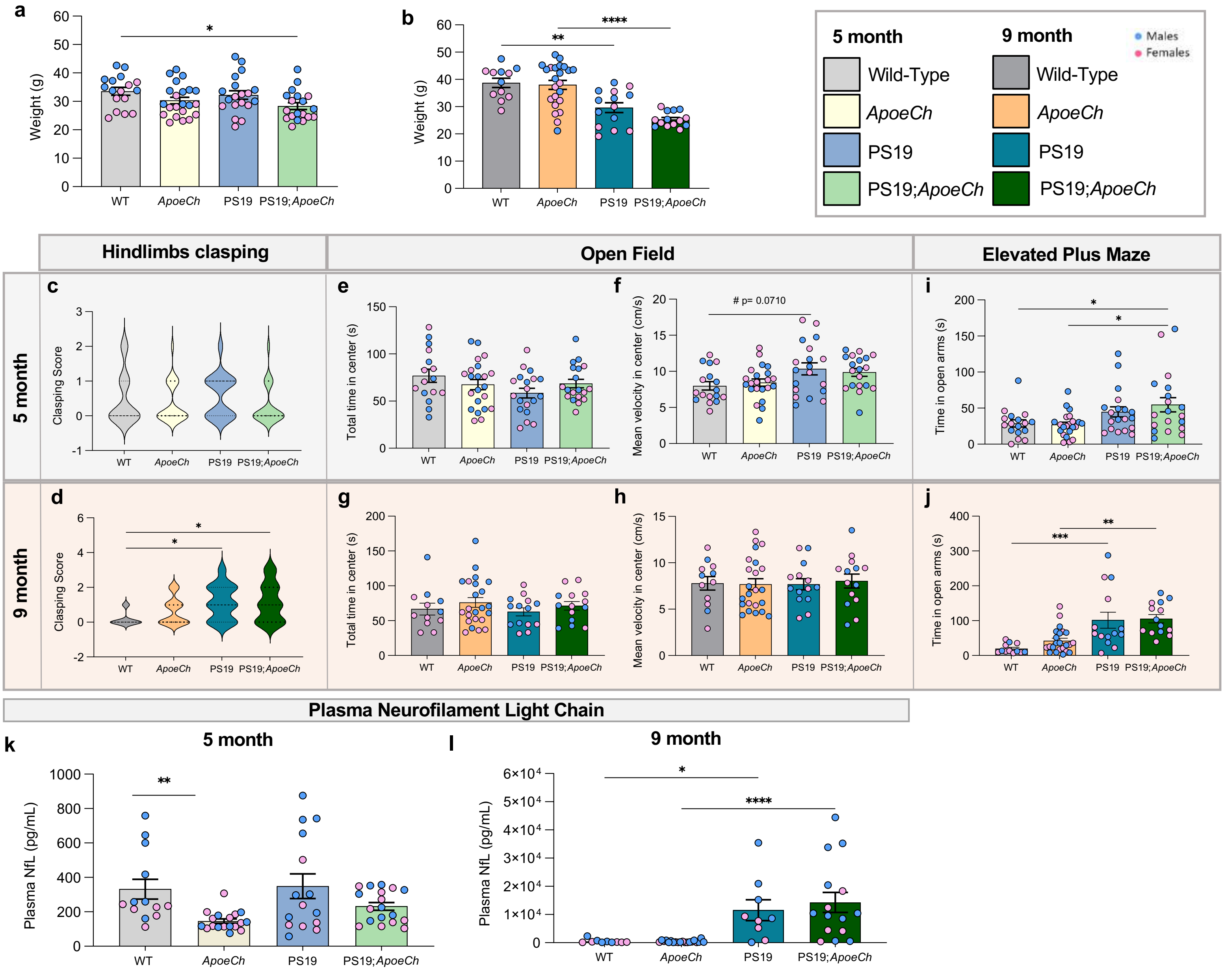


Supplemental Figure 11

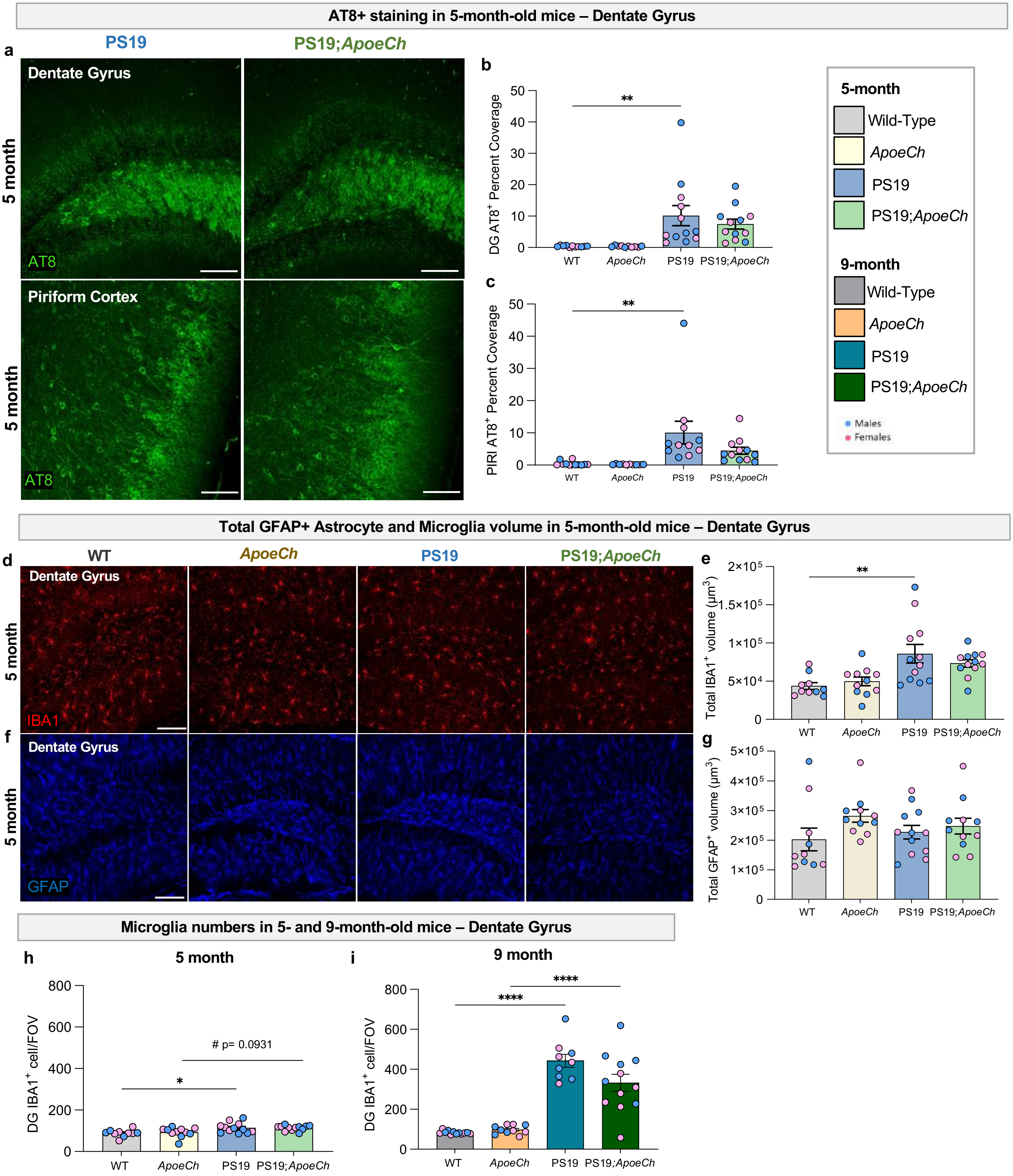


Supplemental Figure 12

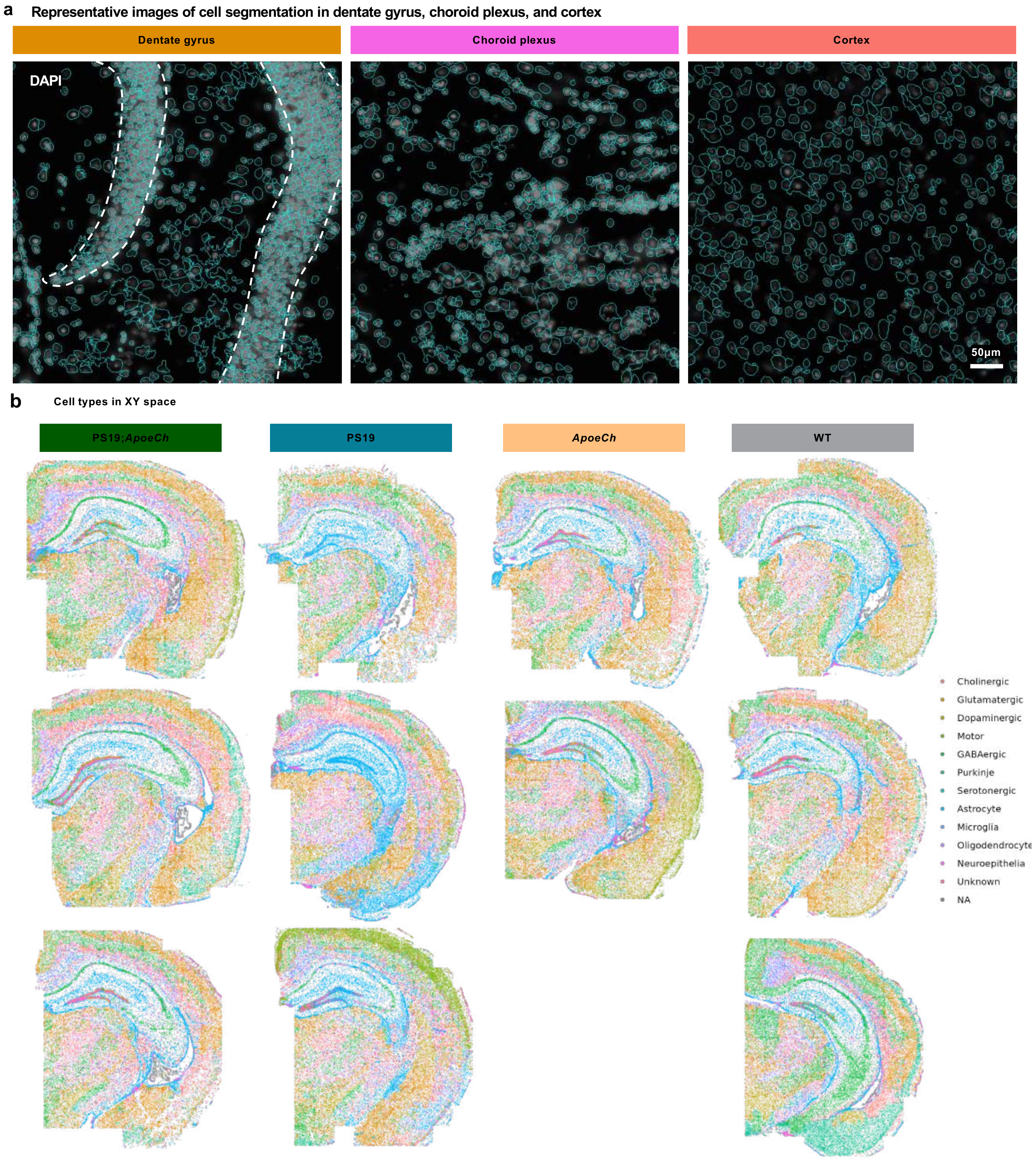


Supplemental Figure 13

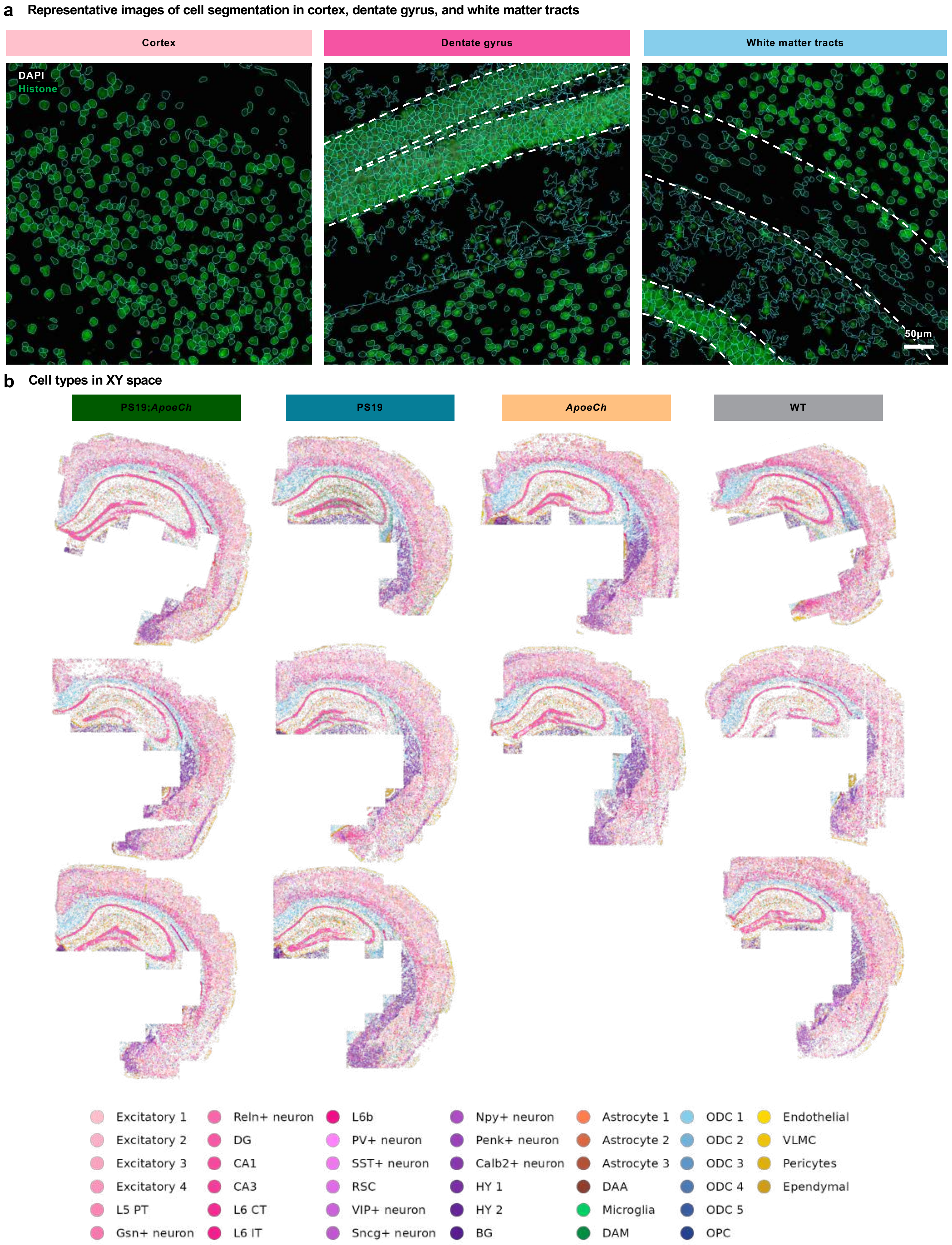


**Supplemental Figure 14**

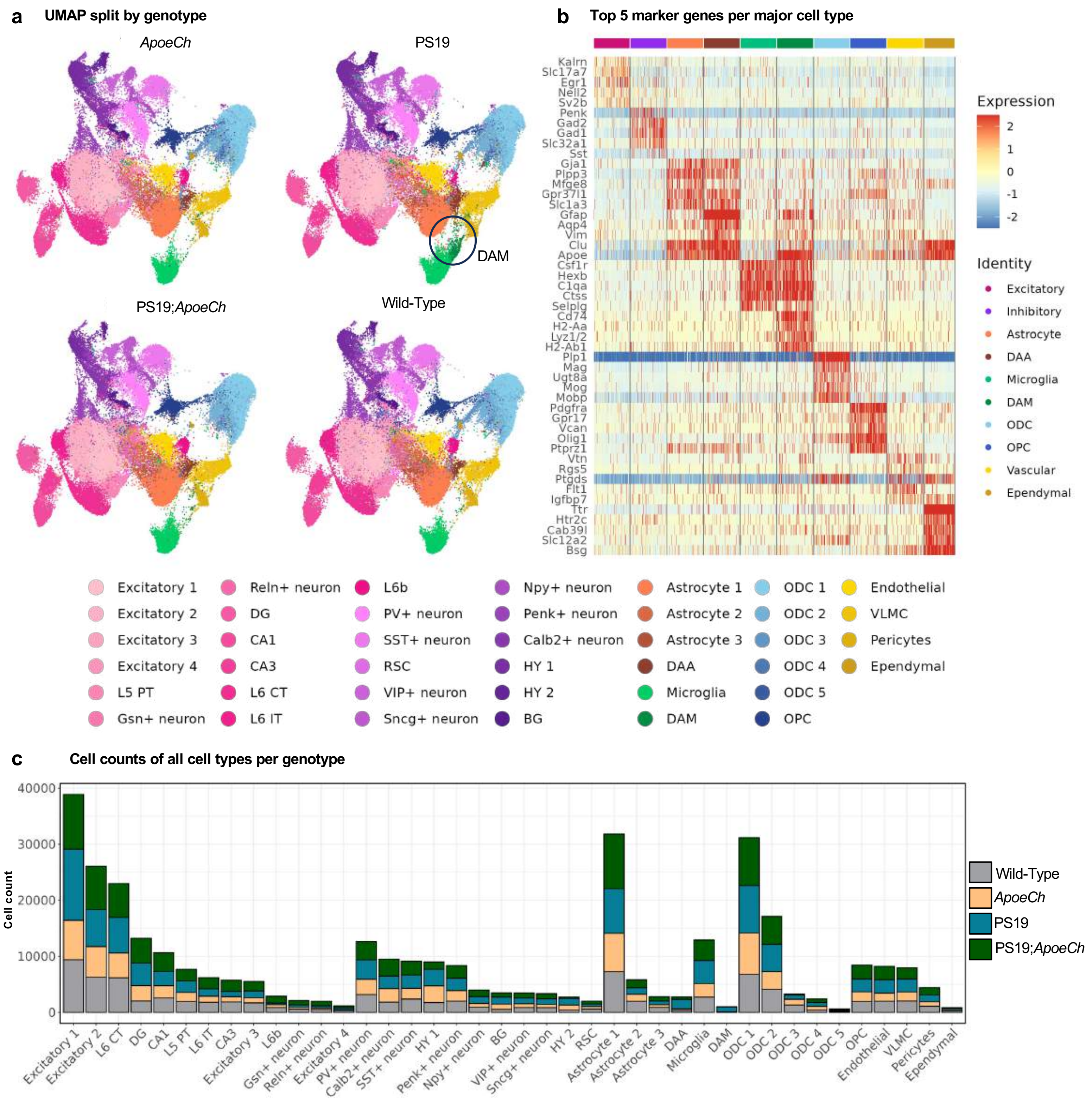


**Supplemental Figure 15**

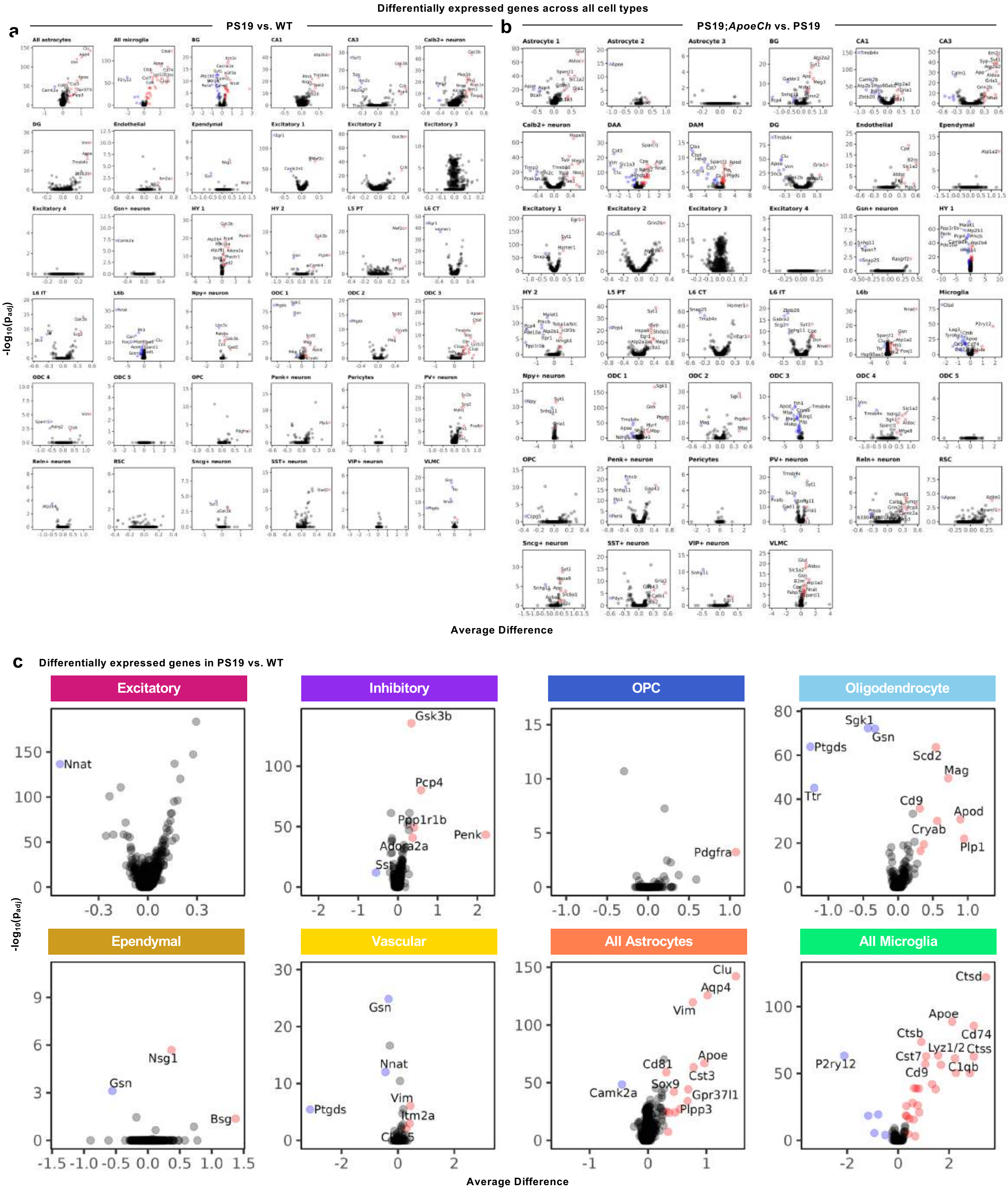


Supplemental Figure 16
